# Genetic control couples TGF*β*-independent pluripotent stem cell renewal to deterministic forward programming across germ layers

**DOI:** 10.64898/2026.10.25.761636

**Authors:** Sveva Bottini, Francesca Amodeo, Divyashree Avnoor, Mario De Gregorio, Maddalena Arigoni, Fabian Bachinger, Philip Barrett, David L. Mack, Paolo E. Porporato, Paola Defilippi, Ludovic Vallier, Alec S. T. Smith, Elisa Balmas, Alessandro Bertero

## Abstract

Reproducible and scalable manufacture of human induced pluripotent stem cell (hiPSC) derivatives requires control over expansion and differentiation. Transcription factor-mediated forward programming can place hiPSC differentiation under genetic control, but expansion still relies on extracellular growth-factor supplementation, increasing process complexity and operator dependence. Here we introduce GERALT (**G**enetically **E**nhanced **R**enewal and **A**utonomous **L**ineage **T**ransdifferentiation), a push–pull genetic circuit assembled through targeted safe-harbour integration that couples NANOG-supported, TGF*β*-independent renewal to doxycycline-inducible lineage programming and simultaneous NANOG silencing. GERALT maintained pluripotency during extended TGF*β* withdrawal in both adherent and suspension culture, yet remained programmable towards myogenic, neuronal, and hepatic fates. Compared with the established OPTi-OX platform in myogenesis, GERALT enabled differentiation in simplified medium while canalising and accelerating commitment. GERALT also supported direct myogenic programming in suspension and enabled direct casting of hiPSCs into hydrogel followed by in-tissue programming, yielding electrically responsive engineered muscle tissues. These results extend genetic control upstream into stem cell renewal, facilitating deployment of new approach methodologies and other biomanufacturing applications.

---

Reliable manufacture of defined mammalian cell states is becoming an enabling capability across biotechnology. Human cell-based new approach methodologies (NAMs), regenerative medicine, and cellular agriculture serve distinct purposes, but all rely on cellular inputs with appropriate identity and function that can be generated reliably, at scale, and with consistent performance across manufacturing sites^1–3^. The need is particularly immediate for NAMs, which are being developed to improve the human relevance of safety and efficacy testing. Regulatory agencies are expanding routes to evaluate, qualify, and incorporate NAMs into regulatory decision-making for defined contexts of use^4,5^, while best-practice frameworks for *in vitro* and stem cell research increasingly emphasise controlled test systems, reproducibility, and transferability^6,7^. As these models become more structurally and functionally sophisticated, manufacture of their cellular components increasingly becomes part of the assay-design problem itself.

Human pluripotent stem cells (hPSCs), including patient-derived human induced pluripotent stem cells (hiPSCs), offer a renewable source for generating diverse differentiated cell types while retaining donor-specific genetic backgrounds^8^. Yet lineage potential alone does not provide a robust manufacturing process. Conventional directed differentiation recreates developmental signalling through timed combinations of morphogens, growth factors, and small molecules^9^. Such developmental recapitulation is particularly valuable when intermediate states or developmental mechanisms are themselves the object of study. For applications in which the priority is instead reproducible production of a defined endpoint cell state, however, these protocols can require prolonged culture, complex media changes, and careful control of starting density, signalling exposure, and intermediate states^10,11^. These process sensitivities are compounded by intrinsic differences in differentiation competence across hPSC lines^12^, propagating variability across batches and complicating process transfer^13^. For such applications, the challenge is therefore not simply to generate more cells, but to produce cells of defined identity, purity, and function through procedures compatible with reproducible manufacture.

Forward programming partially addresses this problem by inducing lineage-defining transcription factors in hPSCs and thereby placing part of the differentiation programme under cell-intrinsic control^14,15^. Compared with transient or randomly integrated transgene-delivery approaches, platforms such as OPTi-OX have refined forward programming through targeted safe-harbour integration and doxycycline-controlled expression, enabling rapid and efficient generation of diverse cell types, including neurons, skeletal myocytes, oligodendrocytes, microglia, and hepatocytes^16–18^. This strategy reduces reliance on sequential extracellular developmental cues during differentiation, but leaves the preceding expansion phase dependent on conventional media-derived signals. Cells are only subsequently exposed to a programme that must overcome the pluripotent state. Expansion and differentiation are therefore controlled by operationally separate systems, even though the signalling and gene-regulatory networks that sustain pluripotency can oppose differentiation, and *vice versa*.

Primed human PSC renewal is maintained by interacting FGF and TGF*β*/Activin–Nodal signalling^19–22^. Among their downstream effects, SMAD2/3 signalling sustains expression of the core pluripotency regulator NANOG, whereas NANOG depletion promotes differentiation^23,24^. This relatively direct relationship offers an opportunity to replace a selected extracellular input with genetic control. Such reinforcement, however, creates a reciprocal requirement: a factor that stabilises renewal during expansion must be removed efficiently once differentiation begins. An integrated architecture that couples these opposing operations could therefore make the transition from pluripotency to a programmed lineage more deterministic.

Here we introduce **G**enetically **E**nhanced **R**enewal and **A**utonomous **L**ineage **T**ransdifferentiation (**GERALT**), a human PSC platform that couples NANOG-supported, TGF*β*-independent renewal to inducible lineage programming and simultaneous NANOG silencing. NANOG reinforcement establishes a stable, although transcriptionally distinct, pluripotent state during TGF*β* withdrawal, while its inducible silencing permits efficient forward programming. The same core architecture can accommodate lineage drivers directed towards derivatives of all three germ layers. In a myogenic setting, GERALT supports prolonged self-renewal in TGF*β*-free medium in both adherent and agitated suspension culture, followed by efficient differentiation, accelerates myogenic programming in simplified FGF2-free medium relative to an isogenic OPTi-OX comparator, and enables hPSCs to be cast into engineered muscle tissues before programming is initiated. These findings provide a strategy for extending cell-intrinsic genetic control across a greater fraction of PSC manufacture, with an immediate proof of concept in NAM-oriented tissue production.

## Results

### Engineering a TGF*β*-independent pluripotent state

GERALT was designed as a push–pull genetic circuit in which constitutive reinforcement of pluripotency supports cell expansion, while doxycycline induction couples activation of a lineage-specifying factor with silencing of the pluripotency module (Fig. 1a). We first engineered and validated the renewal arm of this system. Starting from WTC11 *TTN*-mEGFP reporter hiPSCs (E), we integrated constitutively expressed Tet-On-3G at the *hROSA26* genomic safe-harbor locus to generate ET cells. We then targeted the *CLYBL* locus with a CAG-driven cassette encoding *NANOG* and mCherry, obtaining monoallelic ETN_1_ and biallelic ETN_2_ lines (Fig. 1a; Extended Data Fig. 1a). Site-specific integration without detectable donor-backbone incorporation was confirmed by junctional, loss-of-allele and backbone PCRs (Extended Data Figs. 1b–1c). Both lines homogeneously expressed mCherry, with higher fluorescence in ETN_2_ consistent with its biallelic configuration, and ETN_2_ showed no newly acquired large-scale copy-number alterations (Extended Data Figs. 1d–1e).

**Figure 1.**
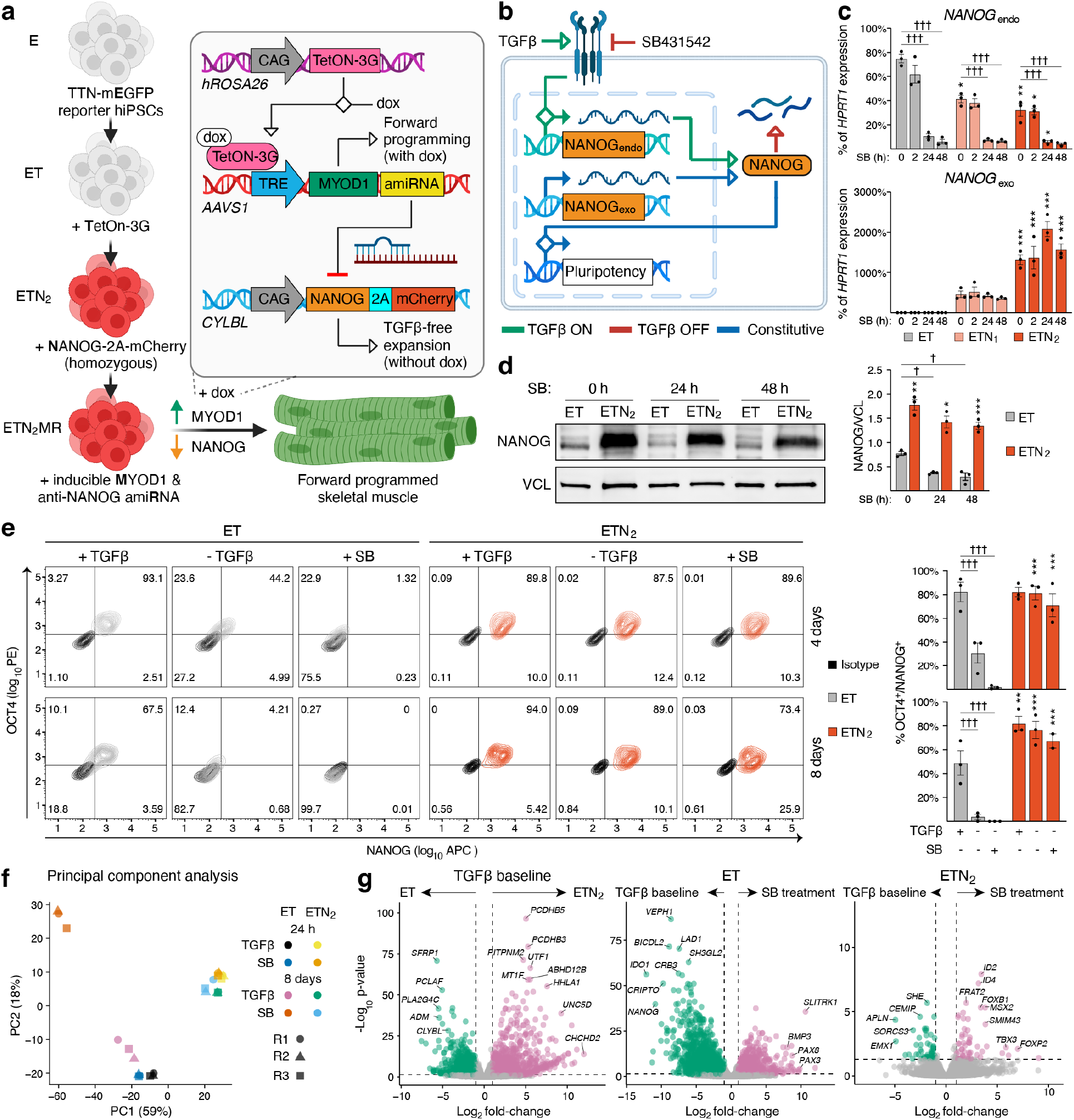
Engineering a TGF*β*-independent pluripotent state. **a**, Overview of GERALT for myogenic programming. Constitutive *NANOG* supports TGF*β*-independent pluripotency, whereas doxycycline (dox) induces *MYOD1* and artificial microRNA (amiRNA)-mediated *NANOG* silencing. The inducible arm is characterized in Fig. 3. Only key elements are shown (not to scale); complete cassette architectures are reported in Extended Data Fig. 1a. **b**, Design rationale for TGF*β*-independent pluripotency. TGF*β* signaling sustains endogenous (endo) *NANOG* transcription through SMAD2/3^23^. SB431542 (SB) inhibits this pathway. In contrast, transgenic (exo) *NANOG* expression is driven constitutively by the CAG promoter, independently of TGF*β* signaling. **c**, RT-qPCR analysis of endogenous (*NANOG*_endo_) and transgenic (*NANOG*_exo_) transcripts in ET, ETN_1_, and ETN_2_ hiPSCs. ETN_1_ and ETN_2_ carry monoallelic and biallelic transgene integration at the *CLYBL* locus, respectively. Cells were maintained in TGF*β*-containing Essential 8 medium (E8; 0 h) or transferred to TGF*β*-free Essential 7 medium (E7) supplemented with SB for 2–48 h. N = 3 independent experiments, each represented by the mean of two parallel cultures. For endogenous *NANOG*, repeated-measures two-way ANOVA followed by Dunnett’s tests compared each ETN line with ET at matched time points (asterisks) and each SB treatment time with 0 h within the same line (daggers). For transgenic *NANOG*, the same model compared ETN_2_ with ETN_1_ at matched time points (asterisks). Here and in panels d and e, *, p < 0.05; **, p < 0.01; ***, p < 0.001; p > 0.05 not reported. **d**, Western blot analysis of NANOG under the conditions in panel c at selected time points. A representative blot and densitometry from N = 3 independent experiments are shown. VCL, loading control. Statistical analysis and notation as in panel c. **e**, Flow cytometry analysis of NANOG and OCT4 expression in ET and ETN_2_ hiPSCs cultured for 4 or 8 days in E8 (+TGF*β*), E7 (-TGF*β*), or E7 supplemented with SB. Representative density plots with isotype controls in black and quantification of NANOG^+^OCT4^+^ cells from N = 2–3 independent experiments are shown. At each time point, a mixed-effects model followed by Dunnett’s tests compared genotypes within each condition (asterisks) and each E7-based condition with E8 within genotype (daggers); notation as in panel c. **f**, Principal component analysis of bulk RNA-seq profiles after 24 h or 8 days under the E8 and E7 plus SB conditions defined in panel c. **g**, Volcano plots of differential gene expression from the experiment in panel f. Left, genotype effect under TGF*β*-containing baseline conditions. Middle and right, treatment-by-time interactions within ET and ETN_2_, respectively. N = 3 independent experiments. Differentially expressed genes were defined by Benjamini–Hochberg-adjusted p < 0.05 and absolute log_2_ fold change > 1.

TGF*β* signaling sustains endogenous *NANOG* transcription through SMAD2/3^23^. We therefore reasoned that constitutive CAG-driven expression of a *NANOG* transgene could bypass this regulatory dependence (Fig. 1b). To test whether this design decoupled NANOG expression from TGF*β* signaling, ET, ETN_1_ and ETN_2_ cells were transferred from Essential 8 medium (E8) to TGF*β*-free Essential 7 medium (E7) containing the ALK4/5/7 inhibitor SB431542 (SB). Endogenous *NANOG* expression was lower in ETN_1_ and ETN_2_ than in ET under E8 conditions, consistent with established negative autoregulation of the endogenous *NANOG* locus^25,26^. SB rapidly reduced endogenous *NANOG* in all three lines, whereas transgenic *NANOG* expression remained high in ETN_1_ and ETN_2_ and increased with transgene dosage (Fig. 1c). Consistently, ETN_2_ maintained substantially higher NANOG protein than ET throughout a 48 h treatment (Fig. 1d). Suppression of *NODAL* and *LEFTY1* confirmed inhibition of the TGF*β* transcriptional program, while *POU5F1/OCT4* was selectively maintained in the engineered lines (Extended Data Fig. 2a).

We next cultured the cells for 4 or 8 days in E8, E7 or E7 supplemented with SB. ET cells progressively lost NANOG and OCT4 expression, with almost complete depletion of the double-positive population following sustained receptor inhibition. In contrast, ETN_1_ and ETN_2_ retained predominantly NANOG^+^OCT4^+^ populations across conditions (Fig. 1e; Extended Data Fig. 2b). RT-qPCR confirmed sustained *POU5F1/OCT4* and transgenic *NANOG* expression in ETN_2_, whereas the early neuroectodermal markers *PAX6, SOX1* and *GBX2* were strongly induced in ET but remained low in ETN_2_ (Extended Data Fig. 2c). Given its biallelic configuration and stronger transgene expression, ETN_2_ was selected for subsequent experiments.

To determine whether marker retention reflected broader preservation of cell state, we performed bulk RNA sequencing after 24 h or 8 days in E8 or E7 supplemented with SB. Principal component analysis separated ET and ETN_2_ along PC1, which accounted for 59% of variance, with a further displacement of longterm SB-treated ET cells (Fig. 1f). Biological replicates remained closely correlated within each condition (Extended Data Fig. 2d). Gene-loading analysis linked the negative PC1 direction to neural-developmental processes, whereas ETN_2_-associated loadings included pluripotency and epithelial-state genes such as *POU5F1/OCT4, CDH1, TDGF1* and *UTF1* (Extended Data Figs. 2e–2f).

Differential expression analyses further distinguished immediate pathway inhibition from progressive changes in cell identity. At 24 h, SB suppressed canonical TGF*β* targets in both genotypes. By day 8, however, ET cells displayed a broader SB-associated neurodevelopmental response, including induction of *PAX3, PAX8, SLITRK1* and *SIX3*, whereas ETN_2_ retained higher expression of *UTF1* and *DPPA5* and did not induce *SIX3* (Fig. 1g; Extended Data Figs. 2g–2h). Thus, constitutive NANOG maintained core pluripotency features despite TGF*β* pathway inhibition while suppressing overt neuroectodermal differentiation.

### NANOG overexpression sustains pluripotency during prolonged TGF*β*-free culture

Having established that NANOG overexpression preserved pluripotency during acute TGF*β* withdrawal or inhibition, we next tested whether this state remained stable during repeated expansion. Three independent ET and ETN_2_ cultures were transferred from E8 to E7 and serially passaged (Fig. 2a). ET cultures underwent marked differentiation and could not be maintained beyond passage 3. At this stage, almost no ET cells co-expressed NANOG and OCT4. Conversely, approximately 90% of ETN_2_ cells remained double positive at passage 3, and substantial double-positive populations persisted through passages 5 and 10, corresponding to approximately 40 days of continuous TGF*β*-free culture (Fig. 2b).

**Figure 2.**
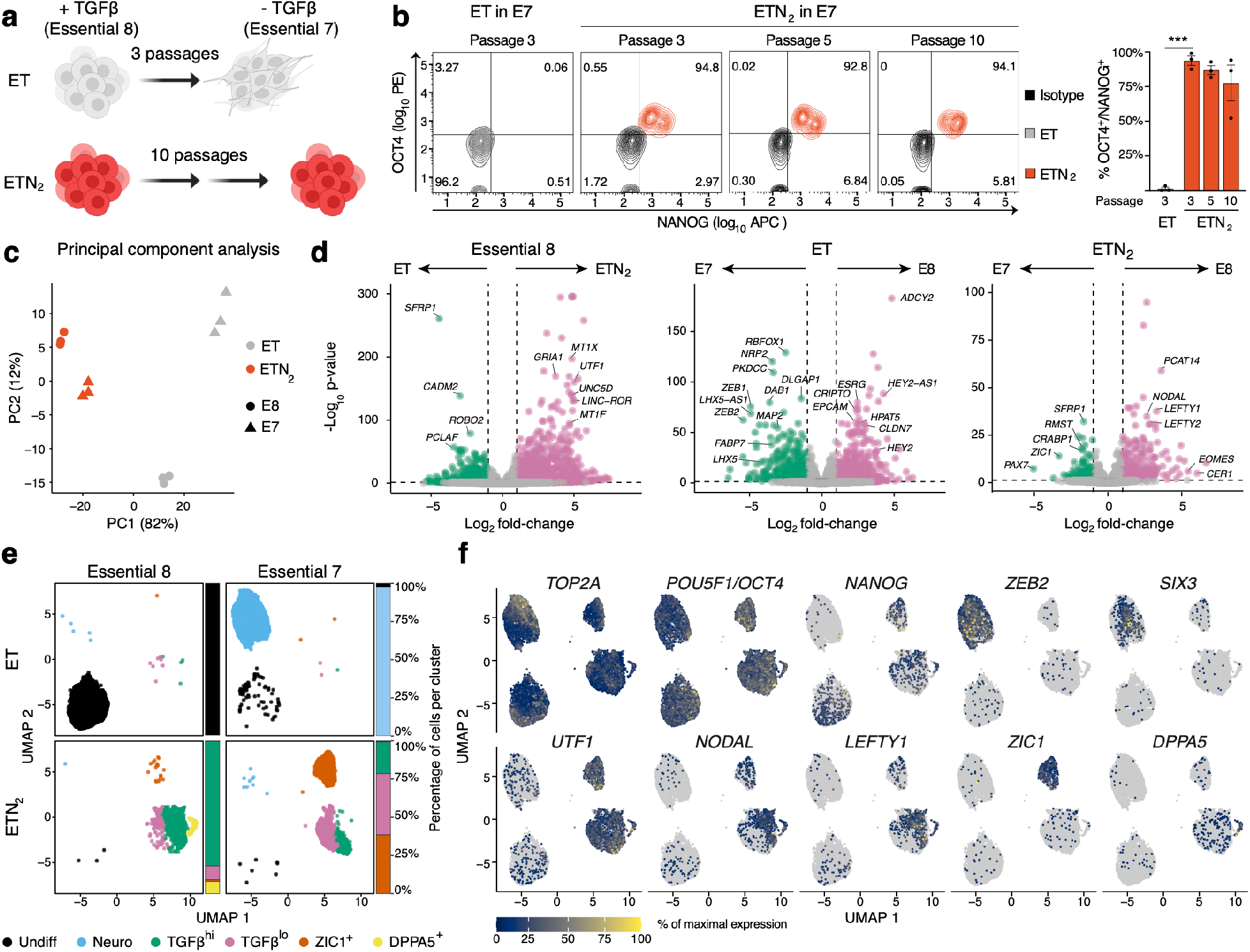
NANOG overexpression sustains pluripotency during prolonged TGF*β*-free culture. **a**, Experimental strategy for long-term culture in TGF*β*-free medium. ET and ETN_2_ hiPSCs were transferred from TGF*β*-containing E8 to TGF*β*-free E7. ET cultures could not be maintained beyond passage 3, whereas ETN_2_ cultures were maintained for 10 passages. **b**, Flow cytometry analysis of NANOG and OCT4 during long-term culture in E7. Representative density plots for ET at passage 3 and ETN_2_ at passages 3, 5 and 10, with isotype controls in black, and quantification of NANOG^+^OCT4^+^ cells from N = 3 independent experiments. Mixed-effects model followed by Šidák’s multiple-comparison test between ET and ETN_2_ at passage 3; *** p < 0.001. **c**, Principal component analysis (PCA) of pseudobulk expression generated by aggregating single-cell RNA-sequencing data for each biological replicate. Cells were analyzed in E8 or after adaptation to E7 for 3 passages (ET) or 10 passages (ETN_2_). N = 3 independent experiments. **d**, Volcano plots of pseudobulk differential gene expression from the experiment in panel c. Differentially expressed genes were defined by Benjamini–Hochberg-adjusted p < 0.05 and absolute log_2_ fold change ≥ 1. **e**, Uniform Manifold Approximation and Projection (UMAP) of single-cell transcriptomes from the experiment in panel c, split by geno-type and culture medium and colored by transcriptional cluster. Cluster labels were assigned from marker expression and condition distribution: Undiff, undifferentiated hiPSCs; Neuro, neuroectoderm-like cells; TGF*β*^hi^ and TGF*β*^lo^, states with higher or lower expression of TGF*β*-responsive genes; and *ZIC1*^+^ and *DPPA5*^+^, clusters defined by the corresponding markers. Bars show cluster composition within each condition. Supporting annotation analyses are reported in Extended Data Fig. 3. **f**, Feature plots of cell-cycle (*TOP2A*), pluripotency (*POU5F1/OCT4, NANOG*), neuroectodermal (*ZEB2, SIX3*), genotype-associated (*UTF1*), TGF*β*-responsive (*NODAL, LEFTY1*), and cluster-defining (*ZIC1, DPPA5*) genes. Expression is scaled independently for each gene.

To define the resulting cell states at transcriptomic resolution, we performed single-cell RNA sequencing on the three independently adapted cultures together with genotype-matched E8 controls. Pseudobulk principal component analysis separated the genotypes along PC1, which accounted for 82% of the variance. The E8-to-E7 transition was substantially larger in ET than in ETN_2_, and biological replicates clustered closely (Fig. 2c). Under E8 conditions, ETN_2_ remained distinguished by higher expression of genes including *UTF1*, whereas ET expressed higher levels of *SFRP1*, consistent with the bulk RNA-seq analysis. E7 adaptation in ET induced a broad neuroectodermal program marked by *RBFOX1, NRP2, ZEB1, ZEB2, MAP2* and *LHX5*, together with reduced expression of epithelial and pluripotency-associated genes. By contrast, ETN_2_ adaptation was characterized primarily by reduced *NODAL, LEFTY1, LEFTY2, EOMES* and *CER1*, accompanied by induction of *ZIC1, RMST* and *CRABP1*, suggesting limited engagement of neural- and retinoid-associated transcription without progression to a comparable broad neuroectodermal state (Fig. 2d).

Unsupervised clustering resolved six transcriptional states (Fig. 2e). ET cells in E8 predominantly formed an undifferentiated (Undiff) cluster, whereas ET cells adapted to E7 largely entered a neuroectoderm-like (Neuro) cluster characterized by loss of *POU5F1/OCT4* and *NANOG* and induction of *ZEB2* and *SIX3* (Fig. 2f).

ETN_2_ cells instead retained expression of core pluripotency markers and occupied two major states distinguished by high or low expression of TGF*β*-responsive genes, termed TGF*β*^hi^ and TGF*β*^lo^. E7 also generated an ETN_2_-specific *ZIC1*^+^ population, whereas a smaller *DPPA5*^+^ population was restricted to ETN_2_ in E8. These states were reproduced across the three independent adaptations, although one ET replicate displayed more extensive loss of *POU5F1/OCT4* and induction of *ZEB2* than the others (Extended Data Fig. 3a; Extended Data Fig. 3d). Cell-cycle phase was distributed across clusters and did not explain their segregation (Extended Data Fig. 3b), while cluster-marker analysis independently supported the cluster annotations (Extended Data Fig. 3c).

Pseudobulk comparison of the Neuro and Undiff clusters confirmed reciprocal upregulation of neuroectodermal and pluripotency-associated programs (Extended Data Fig. 3e). Genes induced in the Neuro cluster were strongly enriched for Gene Ontology terms related to neuronal differentiation, axonogenesis, forebrain development, synapse organization and neuron-projection guidance (Extended Data Fig. 3f). In contrast, the *ZIC1*^+^ and TGF*β*^hi^ ETN_2_ populations were transcriptionally distinct but both retained core features of undifferentiated cells (Extended Data Fig. 3g).

To position these ETN_2_ states within the broader spectrum of human pluripotency, we transferred labels from a reference atlas that resolves conventional, mesoderm-biased and neuroectoderm-biased hiPSC states^27^. After excluding the differentiated Neuro cluster, ETN_2_-derived populations were preferentially assigned to neuro-primed pluripotent references. This association was most pronounced for the mutually exclusive *ZIC1*^+^ and *DPPA5*^+^ states, which emerged in E7 and E8, respectively (Extended Data Fig. 3h). Finally, Milo differential-abundance analysis^28^ resolved these changes across local transcriptional neighborhoods while accounting for biological replicates. In particular, E7 culture increased the abundance of *ZIC1*^+^ neighborhoods while depleting TGF*β*^hi^ and *DPPA5*^+^ neighborhoods within ETN_2_ (Extended Data Fig. 3i).

Together, these analyses show that constitutive NANOG expression enables prolonged hiPSC expansion without exogenous TGF*β* and largely suppresses the broad neuroectodermal differentiation observed in control cells, while redistributing cells among distinct medium-dependent pluripotent states.

### Push–pull programming couples pluripotency exit to deterministic forward programming

Having established a renewal state sustained by constitutive *NANOG*, we next engineered the reciprocal differentiation arm of GERALT. We first tested whether a doxycycline-inducible miR-F cassette^29^ could provide rapid post-transcriptional silencing in hiPSCs. In ET cells, miR-F hairpins targeting the surface proteins CD151 or CD46 selectively depleted their cognate proteins in dox-induced EGFP^+^ cells, whereas an anti-RLuc control did not (Extended Data Figs. 4a–4b). We then transferred the same architecture to ETN_2_, where an anti-NANOG miR-F reduced NANOG abundance specifically in induced cells and extinguished the mCherry reporter linked to the constitutive *NANOG*_exo_ cassette (Extended Data Figs. 4c–4d). These experiments established an inducible silencing module capable of opposing the renewal arm of GERALT.

We next coupled this pluripotency “pull” to the “push” of a lineage-specifying factor by integrating dox-inducible *MYOD1* at *AAVS1*. The first generation ETN_2_MR_N-UTR_ design placed the anti-NANOG miR-F in the 3’ UTR of the *MYOD1* transcript, while an otherwise matched ETN_2_MR_C-UTR_ control carried an anti-CD151 miR-F. We also generated two alternative architectures in which the anti-NANOG miR-F was separated from *MYOD1* using a bidirectional promoter (ETN_2_MR_N-BID_) or embedded within an intron of the *MYOD1* transcript (ETN_2_MR_N-INT_) (Fig. 3a). Targeted integration was verified by genomic PCR across the circuit configurations (Extended Data Fig. 4e); SNP-array profiling largely recapitulated known parental-background alterations and additionally identified copy-number-neutral LoH on chromosomes 2 and 12, together with a small lower-confidence CNV call on chromosome 16 in two clones (Extended Data Fig. 4f).

**Figure 3.**
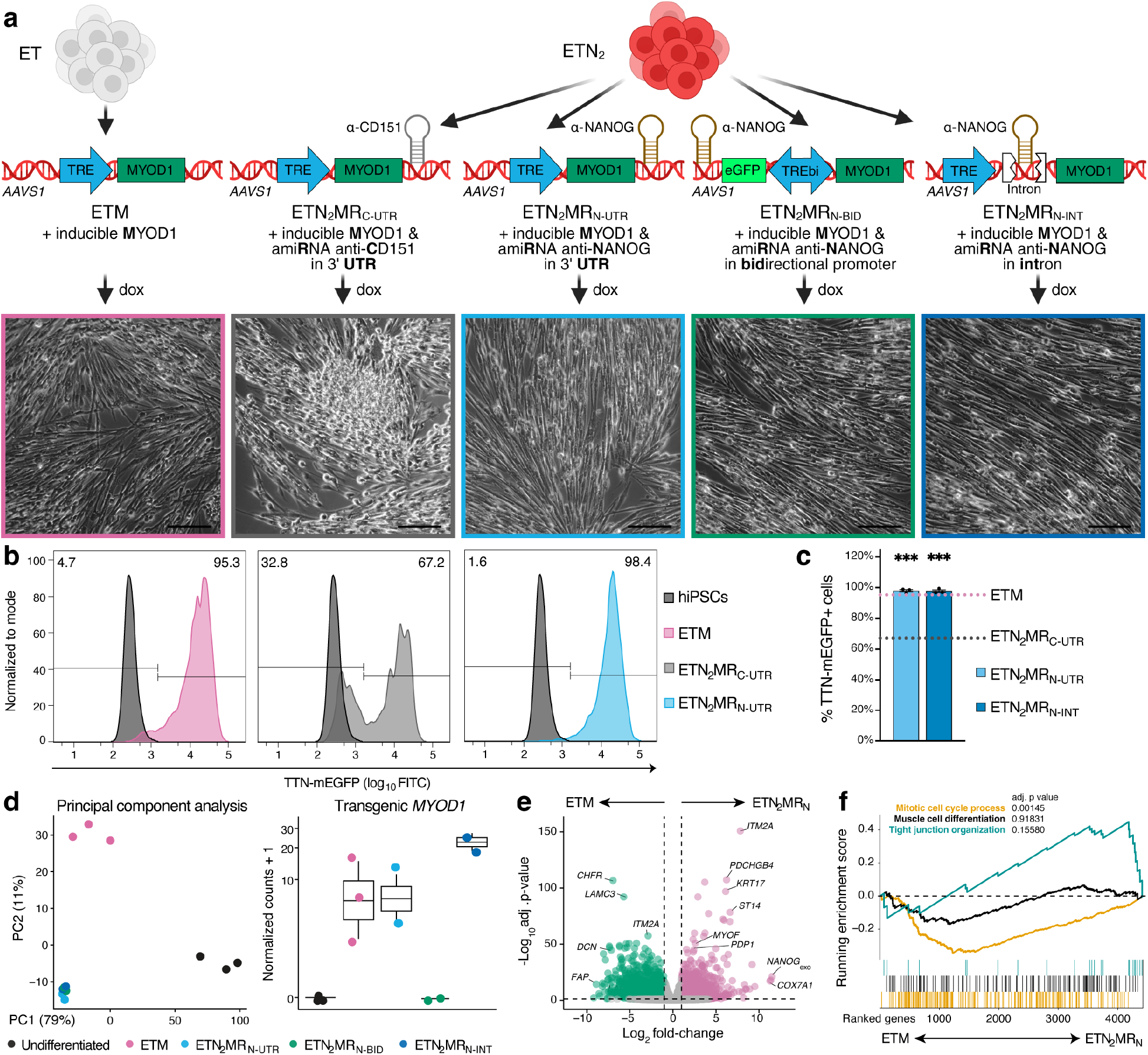
Push–pull programming couples pluripotency exit to deterministic forward programming. **a**, Schematic of the inducible myogenic circuits and representative phase-contrast images after 7 days of dox-induced, *MYOD1*-driven forward programming. ETM carries inducible *MYOD1* alone; ETN_2_MR variants additionally encode the indicated amiRNA (anti-NANOG or anti-CD151 control) in the 3’ UTR of *MYOD1*, from a bidirectional promoter, or within a *MYOD1* intron. Scale bars, 500 µm. **b**, Representative flow-cytometry histograms of the TTN-mEGFP myogenic reporter for the indicated conditions in panel a. Uninduced hiPSCs are shown in dark grey; percentages denote gated fractions. **c**, Quantification of TTN-mEGFP^+^ cells in independently generated ETN_2_MR_N-UTR_ and ETN_2_MR_N-INT_ clones after 7 days of forward programming. Dotted lines indicate the ETM and ETN_2_MR_C-UTR_ reference values from panel b. The N-BID configuration was excluded because its inducible EGFP expression prevents independent quantification of TTN-mEGFP. N = 3 independently generated clones. One-sample t-tests against the ETN_2_MR_C-UTR_ reference; *** p < 0.001; comparisons with the ETM reference were not significant. **d**, Bulk RNA-seq analysis of undifferentiated hiPSCs and the indicated day-7 forward-programmed cells. Left, PCA. Right, transgenic *MYOD1* expression. N = 3 independent experiments for undifferentiated hiPSCs and ETM and N = 2 for each ETN_2_MR_N_ architecture. **e**, Volcano plot of differential gene expression between ETM and pooled ETN_2_MR_N_ samples from panel d. Selected genes are labelled. Differentially expressed genes were defined by Benjamini–Hochberg-adjusted p < 0.05 and absolute log_2_ fold change > 1. **f**, Gene-set enrichment analysis of Gene Ontology Biological Process terms for the comparison in panel e. Curves show running enrichment scores across genes ranked from ETM-enriched to ETN_2_MR_N_-enriched; adjusted p values are shown.

As an isogenic OPTi-OX reference^16^, we generated ETM cells, which carry inducible *MYOD1* but lack constitutive NANOG and the corresponding miR-F. After 7 days of dox-induced forward programming, ETM cells formed elongated myocytes, whereas constitutive NANOG impaired this transition when *MYOD1* was paired with the control anti-CD151 miR-F. By contrast, all three anti-NANOG ETN_2_MR configurations produced extensive myogenic morphology (Fig. 3a), a result reproduced across additional independently generated clones (Extended Data Fig. 4g). Consistent with this phenotype, flow cytometry showed 95.3% TTN-mEGFP^+^ cells in ETM, 67.2% in ETN_2_MR_C-UTR_, and 98.4% in ETN_2_MR_N-UTR_ (Fig. 3b). Independently generated ETN_2_MR_N-UTR_ and ETN_2_MR_N-INT_ clones consistently approached the ETM reference level of TTN-mEGFP induction (Fig. 3c). Immunofluorescence further showed persistent NANOG-mCherry in the control-miRNA condition despite MYOD1 induction, whereas the anti-NANOG UTR circuit combined MYOD1 expression with loss of the NANOG reporter (Extended Data Fig. 4h). RT-qPCR across the alternative configurations confirmed strong repression of transgenic *NANOG* by each anti-NANOG design, with concomitant induction of *TTN* and *MYOG* (Extended Data Fig. 4i). Thus, efficient forward programming in the NANOG-reinforced background required active withdrawal of the engineered pluripotency input rather than lineage-factor induction alone.

**Figure 4.**
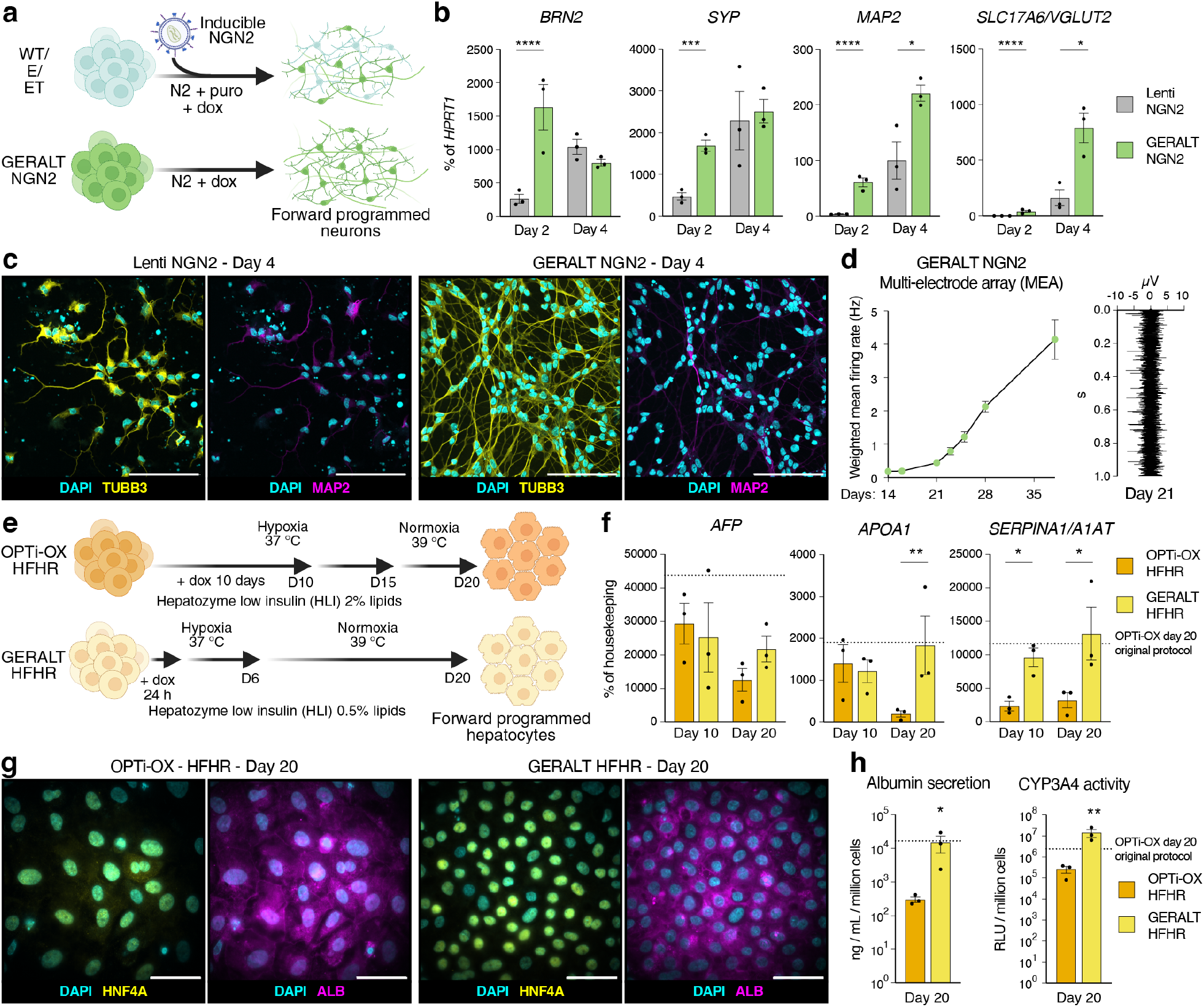
GERALT enables portable forward programming across germ layers. **a**, Benchmarking strategy for GERALT NGN2 neuronal forward programming against lentiviral inducible *NEUROG2/NGN2* delivery and selection in parental WTC11-derived lines (WT, E and ET; Fig. 1a). **b**, RT-qPCR analysis of neuronal markers during forward programming. N = 3 lentiviral-transduced lines and N = 3 GERALT NGN2 clones; each point represents the mean of two independent differentiations. Data shown here and matched homemade Essential 6 (hE6) differentiations in Extended Data Fig. 6g were analysed jointly by three-way repeated-measures ANOVA with platform, medium and time as factors and cell line as the matched experimental unit, followed by Šidák’s multiple-comparison test between platforms within each medium and time point. Here and in panels f and h, * p < 0.05; ** p < 0.01; *** p < 0.001; **** p < 0.0001; p > 0.05 not reported. **c**, Representative immunofluorescence from the experiment in panel b for lentiviral *NEUROG2/NGN2*-transduced ET hiPSCs and the selected GERALT NGN2 clone (Extended Data Fig. 6c). Matched hE6 images and quantification are shown in Extended Data Figs. 6h–6i. Scale bars, 100 µm. **d**, Multi-electrode array analysis of neurons from the selected GERALT NGN2 clone during longer-term culture. Left, weighted mean firing rate from day 14 to day 35 in one representative experiment (n = 14 wells); an independent repeat showed a similar temporal trend. Right, representative 1-s extracellular voltage recording at day 21. **e**, Hepatic forward programming through inducible *HNF1A, FOXA3, ONECUT1/HNF6* and *RORC* (HFHR). The reference OPTi-OX HFHR protocol^31^ is compared with the GERALT-optimized HFHR protocol (Supplementary Fig. 1). In panels f–h, FSPS13B-derived OPTi-OX HFHR and WTC11-derived GERALT HFHR hiPSCs (Extended Data Fig. 7d) were compared under the GERALT-optimized conditions shown here; dotted lines in quantitative panels indicate OPTi-OX HFHR reference values obtained using the reference protocol. **f**, RT-qPCR analysis during forward programming under the conditions in panel e. Expression is normalized to the geometric mean of *PBGD* and *RPLP0*. N = 3 independent differentiations. Two-way repeated-measures ANOVA followed by Šidák’s multiple-comparison test between platforms at each time point; statistical notation as in panel b. **g**, Representative immunofluorescence after 20 days of forward programming under the conditions in panel e. Scale bars, 50 µm. **h**, Albumin secretion and CYP3A4 activity at day 20 under the conditions in panel e. N = 3 independent differentiations. Two-tailed paired t-tests; statistical notation as in panel b.

To compare the resulting myogenic states beyond reporter expression, we performed bulk RNA sequencing after 7 days of forward programming. Undifferentiated hiPSCs separated from all differentiated samples along PC1, which accounted for 79% of the variance, while ETM and the three anti-NANOG configurations occupied distinct regions within the differentiated transcriptional space, with substantial overlap among the latter (Fig. 3d). Replicates clustered closely by condition (Extended Data Fig. 5a). Notably, circuit architecture affected expression of the lineage driver itself: the bidirectional design showed comparatively low transgenic *MYOD1*, whereas the intronic configuration showed the highest expression among the tested anti-NANOG configurations (Fig. 3d).

Pooling the three anti-NANOG configurations revealed extensive transcriptional differences relative to ETM (Fig. 3e; Extended Data Fig. 5b). Gene-set analyses resolved these differences into coherent programs. The pooled anti-NANOG configurations were associated with slow-fiber and neuromuscular junction signatures, whereas ETM retained stronger glycolytic/fast-fiber, cell cycle, satellite cell and myoblast proliferation programs (Fig. 3f; Extended Data Fig. 5c). Consistently, ETM-enriched genes included glycolytic enzymes such as *PGAM2, ENO1, PGK1, LDHA* and *PDK1*, whereas the anti-NANOG configurations were enriched for genes linked to calcium handling, sarcomeric function and oxidative metabolism, including *ATP2A2, MYBPC1, MYL2* and *COX7A1* (Extended Data Fig. 5d).

We finally used the same dataset to distinguish the two most structurally distinct anti-NANOG configurations. Relative to the bidirectional circuit, the intronic design showed a transcriptional bias toward myofiber, sarcomeric, excitation–contraction and neuromuscular-junction programs (Extended Data Figs. 5e–5f). Genes preferentially expressed by the intronic configuration included *MYH8, MYH7, MYH2, CD36* and transgenic *MYOD1*, whereas the bidirectional configuration showed higher expression of genes including *PTPRZ1, NRP2, GRIN3A, DPP6, APCDD1, NOTUM* and *KRT19*, consistent with greater alternative-lineage transcriptional activity (Extended Data Fig. 5e). On the basis of efficient NANOG silencing and TTN-mEGFP induction together with sustained *MYOD1* expression and the more coherent myogenic transcriptional program, we selected ETN_2_MR_N-INT_ as the lead GERALT MYOD1 configuration for downstream studies.

### GERALT enables portable forward programming across germ layers

The GERALT architecture was designed so that the renewal module could remain fixed while the inducible lineage driver was exchanged. We therefore asked whether the same push–pull logic could support cell identities beyond skeletal muscle. As ectodermal and endodermal test cases, we replaced *MYOD1* with either the neuronal determinant *NEUROG2/NGN2*^16^ or a polycistronic hepatic forward-programming cassette encoding *HNF1A, FOXA3, ONECUT1/HNF6* and *RORC* (HFHR)^18^, while retaining intronic anti-NANOG miR-F-mediated silencing (Extended Data Fig. 6a; Extended Data Fig. 7a).

For neuronal programming, we generated three GERALT NGN2 clones and compared them with three WTC11-derived lines (WT, E and ET), each independently transduced with a lentiviral doxycycline-inducible *NEUROG2/NGN2* cassette^14^ (Fig. 4a). The GERALT NGN2 cassette architecture is shown in Extended Data Fig. 6b, and targeted integration was confirmed by genotyping of three clones (Extended Data Fig. 6c). SNP-array profiling showed only alterations also present in parental WT cells in two clones, whereas the third carried additional large-scale copy-number abnormalities on chromosome 7 (Extended Data Fig. 6d). To test whether neuronal programming depended on a dedicated neural medium, we performed matched differentiations in the reference medium (N2)^14^ and homemade Essential 6 (hE6)^30^. Upon induction, GERALT NGN2 showed robust *NEUROG2/NGN2* expression while reducing *NANOG*_exo_ relative to undifferentiated GERALT NGN2 hiPSCs in both media (Extended Data Fig. 6f).

GERALT NGN2 cells acquired neuronal morphology within 2–4 days (Extended Data Fig. 6e) and induced neuronal genes more rapidly than the lentiviral comparator. At day 2 in N2, *BRN2, SYP, MAP2* and *SLC17A6*/*VGLUT2* were all higher in GERALT NGN2 cells; by day 4, expression was broadly comparable between platforms, with *MAP2* and *SLC17A6*/*VGLUT2* remaining higher in GERALT (Fig. 4b). Both approaches generated TUBB3^+^ and MAP2^+^ neurons by day 4 (Fig. 4c). Programming remained effective in hE6, with robust neuronal-marker induction (Extended Data Figs. 6g–6h) and lower fractions of residual SOX2^+^ and NANOG^+^ nuclei than in the lentiviral comparator (Extended Data Fig. 6i). In a separate, longer-term experiment, multi-electrode-array recordings detected spontaneous electrical activity from day 14 and a progressive increase in weighted mean firing rate through day 35; representative extracellular activity was already evident at day 21 (Fig. 4d).

We next applied the same architecture to hepatic programming. GERALT HFHR cells were generated by integrating a TRE-driven intronic anti-NANOG miR-F followed by the *HNF1A*–*FOXA3*–*ONECUT1/HNF6*–*RORC* cassette (Extended Data Fig. 7b). Genotyping confirmed targeted integration in three clones (Extended Data Fig. 7c). SNP-array profiling showed that the selected clone 1 contained only alterations also detected in parental WT cells, whereas clones 2 and 3 carried additional large copy-number alterations on chromosomes 7 and 6, respectively (Extended Data Fig. 7d). Doxycycline robustly activated the HFHR cassette and concomitantly suppressed *NANOG*_exo_, while HNF1A protein was homogeneously induced in the selected clone after 24 h (Extended Data Figs. 7e–7f).

We then empirically adapted the reference OPTi-OX HFHR protocol^31^ to the GERALT architecture (Supplementary Fig. 1). Optimization reduced dox exposure from 10 days to 24 h and HLI lipid supplementation from 2% to 0.5%, together with an earlier transition to the 39°C normoxic maturation phase (Fig. 4e; Supplementary Fig. 1a). GERALT cultures underwent a progressive morphological transition between days 6 and 20, acquiring a cobblestone-like appearance (Extended Data Fig. 7g). For comparison under matched differentiation conditions, OPTi-OX HFHR and GERALT HFHR cells were differentiated under the same GERALT-optimized conditions.

At days 10 and 20, GERALT HFHR cells expressed higher levels of *SERPINA1*, with higher *APOA1* at day 20; no significant difference in *AFP* was detected (Fig. 4f). At day 20, immunofluorescence showed ALB and HNF4A expression in both cultures, alongside apparent differences in nuclear size (Fig. 4g). At this time point, albumin secretion and CYP3A4 activity were higher in GERALT HFHR cells under the matched GERALT-optimized conditions (Fig. 4h). Importantly, GERALT HFHR cells also achieved *SERPINA1* and *APOA1* expression, albumin secretion and CYP3A4 activity in line with OPTi-OX HFHR reference values obtained using the reference protocol (Figs. 4f, 4h).

Two other independently generated GERALT HFHR clones also expressed *ALB* and *CYP3A4* and showed measurable albumin secretion and CYP3A4 activity despite their clone-specific copy-number alterations (Extended Data Fig. 7h; Extended Data Fig. 7j). BODIPY staining further demonstrated intracellular lipid accumulation in both OPTi-OX HFHR and GERALT HFHR cultures after 10 days of forward programming (Extended Data Fig. 7i).

Together with the MYOD1 data, these experiments extend the inducible activation-plus-pluripotency-silencing logic to mesodermal, ectodermal and endodermal outputs. Thus, GERALT is not restricted to a single master regulator or lineage, and the renewal chassis can be coupled to distinct forward-programming modules without redesigning its constitutive pluripotency-support layer.

### Minimal media support rapid GERALT myogenic programming

Having established cross-lineage portability, we returned to the myogenic system to determine whether GERALT MYOD1 could simplify the differentiation environment and alter the kinetics or fidelity of forward programming. We compared GERALT MYOD1 and OPTi-OX in homemade Essential 7 (hE7) and hE6^30^ *versus* the standard OPTi-OX myogenic medium^16^ previously used in Fig. 3. Relative to the myogenic medium, hE7 omits the retinoicacid and CHIR99021 patterning inputs, while hE6 additionally omits FGF2 (Fig. 5a; Extended Data Fig. 8a). This provided a stringent test of whether a genetically encoded push–pull circuit could reduce reliance on exogenous differentiation cues.

**Figure 5.**
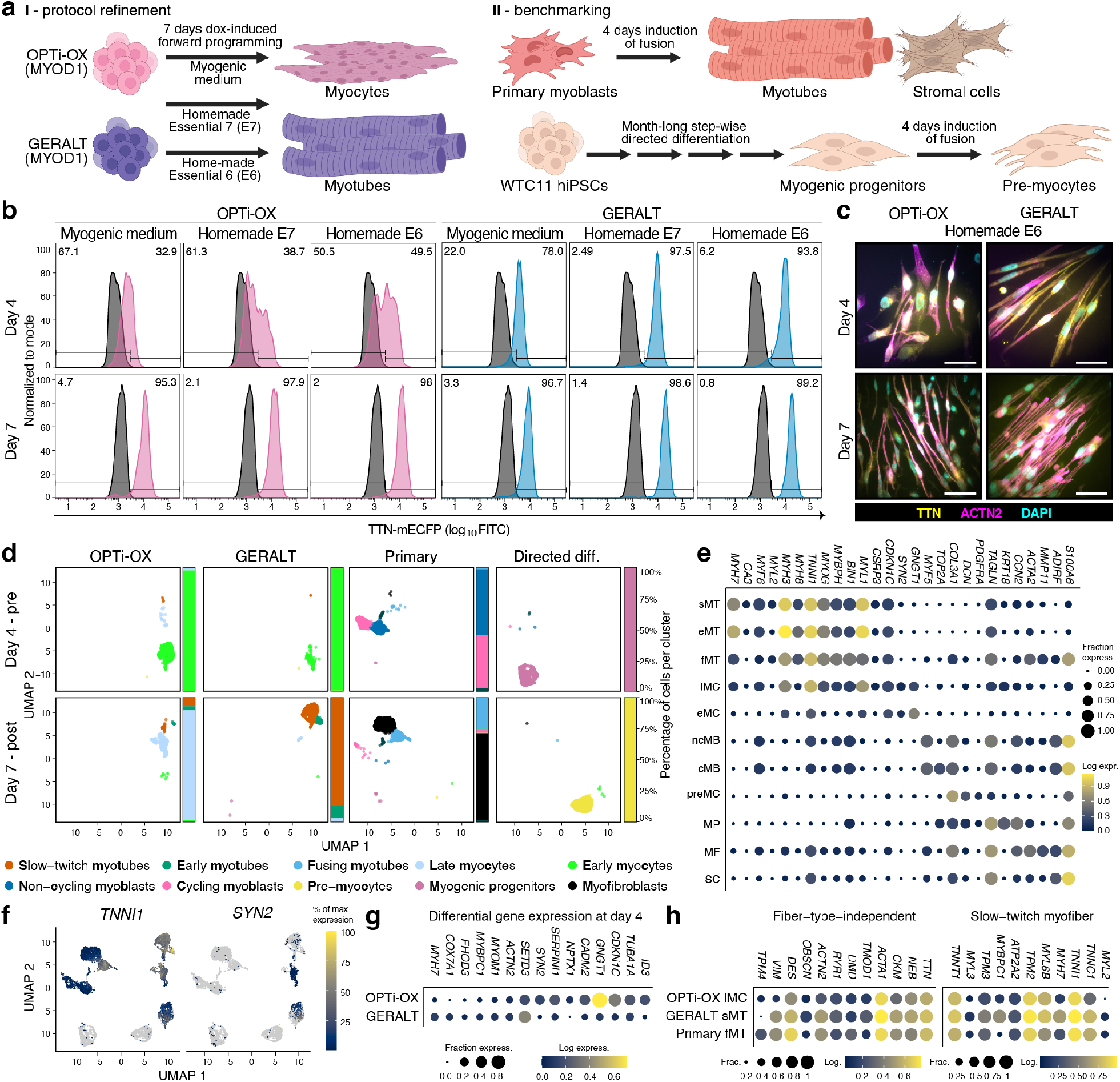
Minimal media support rapid GERALT myogenic programming. **a**, Experimental strategy for refinement and benchmarking of myogenic differentiation. OPTi-OX and GERALT MYOD1 hiPSCs (ETM and ETN_2_MR_N-INT_, respectively, in Fig. 3a) were forward programmed in myogenic medium^16^, homemade Essential 7 (hE7), or hE6 and analysed at days 4 and 7. For benchmarking, primary human myoblasts were analysed before and after 4 days of fusion, and WTC11 hiPSCs after step-wise myogenic directed differentiation and 4 days of fusion. **b**, Representative TTN-mEGFP flow-cytometry histograms for the forward-programming conditions in panel a. Uninduced hiPSCs are shown in dark grey; percentages denote gated fractions. Additional time points and quantification are shown in Extended Data Fig. 8b. **c**, Representative immunofluorescence after 4 and 7 days of forward programming in hE6. Scale bars, 50 µm. **d**, UMAP of single-cell transcriptomes from the conditions in panel a, split by sample type and time/state (pre- and post-fusion) and colored by transcriptional cluster. Bars show cluster composition within each sample. **e**, Dot plot of selected marker genes supporting the cluster annotations in panel d. Dot size indicates the fraction of cells expressing each gene and color indicates log expression. Cluster abbreviations correspond to the names in panel d. **f**, Feature plots of the myogenic gene *TNNI1* and neuronal-associated gene *SYN2*. Expression is scaled independently for each gene. **g**, Dot plot of selected myogenic and alternative-lineage genes differentially expressed between OPTi-OX and GERALT at day 4. Plotting as in panel e. Differentially expressed genes were defined by Bonferroni-adjusted *p* < 0.01 and absolute log_2_ fold change ≥ 1. **h**, Dot plots comparing fiber-type-independent and slow-twitch myofiber gene programs in the indicated sample-associated clusters. Plotting as in panel e. Complementary fast-twitch and mature-muscle-absent gene programs are shown in Extended Data Fig. 9g.

GERALT initiated the TTN-mEGFP program markedly earlier than OPTi-OX across all three media. In representative day-4 cultures, TTN-mEGFP was detected in 78.0% of GERALT cells in myogenic medium, 97.5% in hE7 and 93.8% in hE6, compared with 32.9%, 38.7% and 49.5%, respectively, in OPTi-OX. By day 7, both systems converged on predominantly TTN-mEGFP^+^ populations in all conditions (Fig. 5b). Early TTN-mEGFP induction in GERALT was reproduced at days 3 and 5 (Extended Data Fig. 8b) and was accompanied by extensive cell elongation in simplified media (Extended Data Fig. 8c). Consistently, GERALT cells in hE6 already displayed prominent TTN-mEGFP and ACTN2-positive myofibers at day 4, whereas OPTi-OX cultures showed less developed myofibers; by day 7 both systems formed abundant ACTN2^+^TTN-mEGFP^+^ myotubes (Fig. 5c; Extended Data Fig. 8e). RT-qPCR analysis of *MYOG* and *TTN* across media and time points supported the same accelerated myogenic transition (Extended Data Fig. 8d).

We next asked whether this kinetic difference reflected distinct cell-state trajectories rather than reporter induction alone. We profiled OPTi-OX and GERALT cells by single-cell RNA sequencing at days 4 and 7 in hE6 and analysed them together with primary human myoblast and stepwise directed-differentiation benchmarks^32^ before and after the corresponding fusion/maturation interval (Fig. 5a; Fig. 5d). Integrated analysis resolved myogenic progenitors, cycling and non-cycling myoblasts, early and late myocytes, pre-myocytes, fusing and slow-twitch myotubes, as well as stromal and myofibroblast states (Figs. 5d–5e). The major transcriptional partitions were not obviously associated with the examined quality-control metrics or cell-cycle scores (Extended Data Figs. 9b–9c), and cluster identities were independently supported by markers including *MYH7, MYF5, MYOG, MYBPH, MYL1, COL3A1, PDGFRA, TAGLN* and *ADIRF* (Extended Data Fig. 9d).

At day 4, OPTi-OX and GERALT occupied overlapping but transcriptionally distinct early myogenic states. Feature plots localized the contractile gene *TNNI1* and the neuronal-associated gene *SYN2* to different regions of the integrated UMAP (Fig. 5f), and direct differential-expression analysis revealed higher expression of genes associated with non-myogenic lineages in OPTi-OX alongside greater expression of myogenic genes in GERALT (Fig. 5g; Extended Data Fig. 9e). Among the alternative-lineage genes, *GNGT1* was one of the most strongly OPTi-OX-enriched, whereas GERALT showed higher expression of genes including *MYH7* and *COX7A1* (Extended Data Fig. 9e). These data indicate that the faster GERALT transition was accompanied by earlier consolidation of the myogenic program rather than simply earlier activation of the TTN reporter.

By day 7, both forward-programming systems had progressed toward differentiated muscle states, but their transcriptional identities remained distinct. Differential-expression analysis highlighted greater expression of slow-twitch and fiber-type-independent muscle genes in GERALT, including *MYH7, MYBPC1, TPM3* and *MYL2*, while OPTi-OX retained a broader developmental signature, with higher expression of *GNGT1, KRT19* and *ID3*, as well as the fast-twitch myosin *MYH2* (Extended Data Fig. 9f). At the cluster level, the GERALT slow-twitch-myotube population showed a strong fiber-type-independent and slow-twitch expression program relative to the OPTi-OX late-myocyte population and the primary-myoblast benchmark (Fig. 5h). Complementary analysis showed differences across the same populations in gene programs associated with fast-twitch-myofibers and in those comprising genes typically absent from mature skeletal muscle (Extended Data Fig. 9g). We therefore conclude that GERALT not only supports efficient endpoint myogenesis in media lacking the major exogenous patterning inputs, but also accelerates lineage commitment and shifts the resulting transcriptional state toward a slow-twitch-like program.

### GERALT supports suspension bioprocessing and functional engineered muscle tissues

We next tested two downstream properties relevant to production and model implementation: compatibility with suspension culture and the ability to generate functional three-dimensional engineered muscle tissues (EMTs). These experiments were performed as parallel demonstrations rather than as a continuous suspension-to-EMT process.

For suspension expansion, the lead GERALT myogenic line was transferred to agitated hE7 culture and maintained for 10 passages before dox-induced differentiation was initiated directly in suspension (Fig. 6a). GERALT formed compact aggregates that retained homogeneous mCherry expression from the constitutive NANOG cassette (Fig. 6b) and expanded over successive passages (Extended Data Figs. 10a–10b). After 10 passages, approximately 95% of cells remained NANOG^+^OCT4^+^ across three independent suspension adaptations (Fig. 6c), supporting maintenance of the TGF*β*-independent renewal phenotype outside adherent culture.

**Figure 6.**
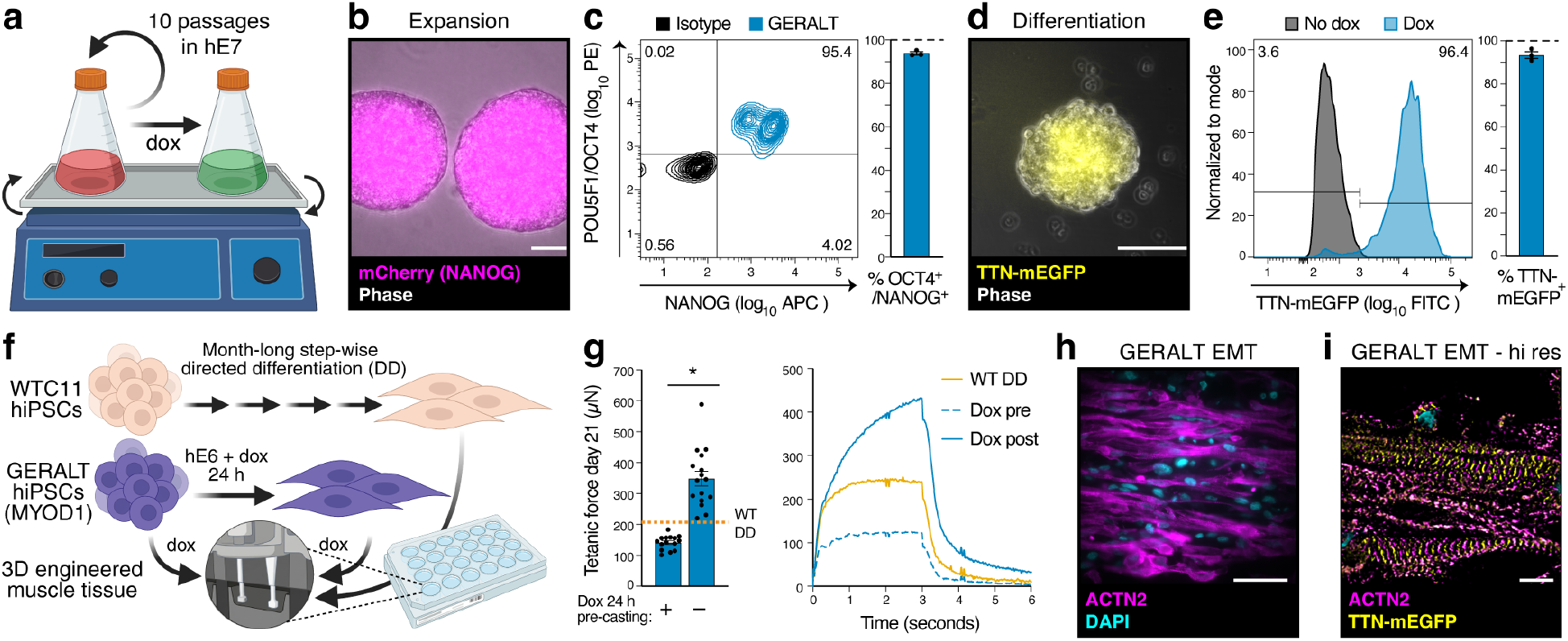
GERALT supports suspension bioprocessing and functional engineered muscle tissues. **a**, GERALT hiPSC expansion for 10 passages in suspension in hE7 followed by myogenic forward programming in suspension. **b**, Representative images of passage 10 hiPSC aggregates. mCherry reports constitutive *NANOG*_exo_. Scale bars, 100 µm. **c**, Representative NANOG and OCT4 flow cytometry after 10 passages in suspension and quantification of NANOG^+^OCT4^+^ cells from N = 3 independent suspension adaptations. Isotype control is shown in black. **d**, Representative images after 7 days of myogenic forward programming in suspension. Scale bars, 100 µm. **e**, Representative TTN-mEGFP flow cytometry and quantification after suspension differentiation, comparing uninduced and doxtreated cells. N = 3 independent differentiation experiments. **f**, Schematic of engineered muscle tissue (EMT) generation. WTC11 directed-differentiation myoblasts (WT DD; Fig. 5a) were compared with GERALT hiPSCs induced for 24 h before casting (dox pre) or after casting (dox post). **g**, Day 21 tetanic force in GERALT dox-pre and dox-post EMTs. Individual EMTs are shown; the dotted line indicates the WT DD reference value. Right, representative tetanic force traces. N = 3 independent casting batches and n = 15–16 EMTs per GERALT condition. Two-tailed nested t-test; * p < 0.05. **h**, Representative immunofluorescence in longitudinal section of a day 21 GERALT dox-post EMT. Scale bars, 50 µm. **i**, Structured illumination microscopy (SIM) of a GERALT dox-post EMT showing sarcomeric organization. Scale bars, 5 µm.

Doxycycline addition to suspension cultures triggered myogenic programming without an intervening adherent phase. TTN-mEGFP-positive aggregates were already evident by day 4 (Extended Data Fig. 10c), and by the final endpoint approximately 95% of cells were TTN-mEGFP^+^ across independent differentiations (Figs. 6d–6e). Thus, the same GERALT line could undergo prolonged growth in aggregate culture and subsequently execute the programmed myogenic switch in suspension.

We then asked whether GERALT could simplify the generation of functional EMTs by initiating programming within the three-dimensional tissue itself. As a reference, we used WTC11 hiPSCs subjected to stepwise directed differentiation before casting, a workflow requiring over a month of culture, selection and expansion. GERALT cells were instead either exposed to hE6 + dox for 24 h before casting (dox pre) or cast directly as hiPSCs and induced after casting (dox post) (Fig. 6f). All conditions formed macroscopically stable tissues between days 7 and 21 (Extended Data Fig. 10d). At day 21, GERALT dox-post EMTs generated greater absolute tetanic force than dox-pre EMTs prepared with matched input cell numbers (Fig. 6g), with the WT directed-differentiation reference value shown for comparison.

GERALT EMTs also displayed frequency-dependent force production and measurable fatigue during repeated stimulation (Extended Data Fig. 10e). Across the analysed tissues, both GERALT conditions retained a larger fraction of initial force over repeated contractions than the WT directed-differentiation reference. Histological analysis showed extensive ACTN2-positive muscle structures throughout GERALT EMTs (Fig. 6h; Extended Data Fig. 10f), and high-resolution imaging resolved aligned ACTN2 and TTN-mEGFP sarcomeric structures (Fig. 6i). RT-qPCR further distinguished the tissue conditions, with dox-post GERALT EMTs showing relatively high expression of contractile genes including *ACTA1, MYH3, MYBPC2* and *TNNI1* (Extended Data Fig. 10g).

Finally, we benchmarked GERALT dox-post EMTs against EMTs generated from a second WT directed-differentiation batch and from primary human myoblasts. GERALT dox-post EMTs generated greater absolute tetanic force than the directed-differentiation controls but remained below primary-myoblast EMTs (Extended Data Fig. 10h). Cross-sections confirmed muscle-marker expression in all three tissue types, with GERALT EMTs appearing larger but showing reduced muscle-marker staining in their central regions (Extended Data Fig. 10i). Together, these data establish that GERALT is compatible with prolonged suspension expansion and suspension myogenic programming, while also supporting generation of electrically responsive, sarcomere-containing engineered muscle tissues directly from hiPSCs.

## Discussion

GERALT extends cell-fate engineering into a phase that forward programming systems have left under extracellular control. By reinforcing NANOG during expansion and withdrawing that reinforcement as lineage programming begins, the platform links renewal and differentiation as opposing states of the same engineered system. The pluripotent state generated by this intervention is not equivalent to conventionally cultured hPSCs: transgenic NANOG altered transcription even in complete medium, and prolonged TGF*β* withdrawal redistributed cells among pluripotency-associated, TGF*β*-responsive and *ZIC1*-expressing states. Despite these changes, cultures continued to expand, retained core pluripotency markers, largely avoided the broad neuroectodermal differentiation seen in control cells, and remained amenable to forward programming. This suggests that, for a manufacturing chassis, the relevant criterion may be a stable and programmable starting state rather than strict molecular equivalence to conventionally cultured hPSCs.

The MYOD1 experiments show why reinforcement of renewal must be coupled to an efficient exit mechanism. Continued NANOG expression impaired myogenic conversion despite induction of the lineage driver, whereas anti-NANOG silencing restored efficient programming. The push–pull logic is therefore functionally consequential rather than merely a feature of circuit design. Its implementation also mattered: placing the amiRNA in the 3’ UTR, under a bidirectional promoter, or within an intron produced different relationships between NANOG repression, MYOD1 expression and transcriptional identity. In the myogenic setting, the intronic configuration achieved the most favorable balance and was therefore used as the architecture for the NGN2 and HFHR circuits.

Portability across myogenic, neuronal and hepatic programs shows that the core GERALT logic is not restricted to one germ layer, although lineage-specific optimization of induction and culture conditions may still be required. For neuronal programming, we deliberately used a widely adopted lentiviral NGN2 system as an orthogonal benchmark^14^. The hepatic experiments addressed a different question: whether an established OPTi-OX program could be transferred to GERALT and still yield functional hepatic-like cells^18,31^. Direct transfer was suboptimal, but empirical adaptation of lineage-driver induction, lipid supplementation and maturation timing supported robust hepatic programming. The final protocol shortened HFHR induction to 24 h, reduced lipid supplementation to 0.5%, and advanced the transition to the 39°C normoxic maturation phase. Because these changes were selected empirically rather than through isolated mechanistic tests, why GERALT performed better under a different operating regime remains unresolved. Possibilities include altered effective lineage-driver kinetics, coupling of HFHR induction to NANOG withdrawal, or the distinct starting state established by NANOG reinforcement.

The closely matched MYOD1 comparison provides the clearest view of how the renewal module influences subsequent programming. GERALT and OPTi-OX share the WTC11 ET background and the same inducible MYOD1 programming framework, with NANOG-supported renewal and coupled anti-NANOG silencing constituting the only intended distinction. GERALT cells entered the myogenic program earlier, displayed less alternative-lineage transcription at early time points, and later acquired a stronger slow-twitch and fiber-type-independent signature. These differences cannot be reduced to TTN-reporter induction alone and suggest that coupling withdrawal of reinforced pluripotency to MYOD1 activation can both accelerate and alter the trajectory of myogenic programming. This is consistent with a broader view of forward programming in which lineage conversion depends on both the strength of the lineage-driving program and the resistance imposed by the starting state. Consistent with this concept, recent combinatorial CRISPRa/i screening identified ID3 repression together with SMARCD3/BAF60C activation as one route to more canalized MYOD1-induced myogenesis^33^.

The suspension and engineered tissue experiments put this architecture to the test in manufacturing-relevant formats. GERALT maintained prolonged TGF*β*-independent expansion in agitated suspension and could subsequently undergo myogenic programming without returning to adherent culture, demonstrating that both renewal and programming can operate in aggregate culture. In parallel, hPSCs could be cast directly into engineered muscle tissues and programmed after casting, during tissue formation, yielding organized contractile structures with stimulus-responsive function. Moving differentiation closer to the assay endpoint may simplify production of tissue-based NAMs by reducing the period for which differentiated cellular intermediates must be handled separately.

The present study also defines several boundaries of the platform. All GERALT lines derive from WTC11, so the extent to which the phenotypes generalize across hPSC backgrounds remains unresolved despite the use of multiple independent clones. The current architecture further requires repeated genome editing and clonal selection, making both engineering efficiency and clone-level genomic quality control relevant to future deployment^34,35^, as underscored by the additional copy-number alterations detected in subsets of the NGN2 and HFHR clones. Likewise, agitated suspension demonstrates format compatibility rather than performance under the controlled, high-density bioreactor conditions required for industrial-scale bioprocessing^36^. The renewal phase is specifically TGF*β*-independent rather than generally growth-factor-independent, as FGF2 and insulin/IGF signaling remain present. The current MYOD1 EMT implementation also maintained doxycycline throughout tissue maturation, so future work should determine whether shorter induction windows or self-limiting control architectures can preserve function. Finally, the engineered-muscle-tissue data demonstrate functional feasibility, but regulatory use as a NAM would require further optimization of tissue geometry and internal muscle organization, reproducible manufacturing and pharmacological performance within a defined context of use, together with appropriate bench-marking and transfer across laboratories^7,37^.

Although developed here around NAM-oriented hPSC manufacture, the underlying GERALT principle is relevant to other settings in which cell state must be controlled reproducibly at scale. Cell therapy presents the clearest biomedical example, but application of the current architecture would require considerably more stringent control of genomic safety, residual pluripotency, transgene behavior and clinical manufacturing^38,39^. Cellular agriculture presents a different set of constraints, but the biological rationale for translation is already substantial: scalable cell production is central to the field^40^, MYOD1-driven forward programming has already been demonstrated in porcine iPSCs^41^, and Activin-dependent pluripotent-state maintenance extends across agriculturally relevant species including pigs and cattle^42,43^. Practical implementation would nevertheless have to address species-specific engineering and the product- and jurisdiction-dependent acceptability of genetically modified cells and doxycycline-responsive circuits^44,45^. These are application-specific engineering challenges rather than limitations of the underlying logic. GERALT therefore provides a NAM-first demonstration of a broader strategy: shifting a greater share of mammalian cell manufacture from repeatedly imposed extracellular conditions to programs encoded within the cell itself.

## Methods

### hiPSC adherent culture and TGF*β* modulation

All hiPSC lines used in this study are detailed in Supplementary Table 1. Briefly, all GERALT lines and the OPTi-OX MYOD1 reference line were derived from the WTC11 hiPSC line, originally established from a healthy male donor^46^ and subsequently engineered by the Allen Institute for Cell Science to carry an endogenous *TTN*-mEGFP reporter (AICS-0048, clone 39)^47^. The Tet-On-3G-expressing derivative used as the parental line for subsequent engineering (ET) was described previously^48^. The OPTi-OX HFHR reference line was the FSPS13B-derived line previously described in Tomaz et al.^18^. Engineered clonal hiPSC lines were used for experiments between passages 3 and 20 after clonal isolation. Cell identity and genomic stability were assessed by molecular karyotyping or SNP-array analysis, as indicated in Supplementary Table 1 and described below. Cultures were routinely tested for mycoplasma contamination by PCR and were negative.

Unless otherwise indicated, adherent hiPSCs were maintained on tissue-culture plates coated overnight at 4°C with 50 µg mL^−1^ hESC-qualified Geltrex (Thermo Fisher Scientific, A1413202). Cells were cultured at 37°C in a humidified atmosphere containing 5% CO_2_. Initial cultures were maintained in commercial Essential 8 (E8; Thermo Fisher Scientific, A1517001), with daily medium replacement and passaging every 4–5 d using 0.5 mM EDTA (Sigma-Aldrich, 4055) in Ca^2+^/Mg^2+^-free PBS. Thiazovivin (Cayman Chemical, 14245) was added at 2 µM for 24 h after reseeding.

Subsequent cultures used homemade media as previously described^30^. Parental and control lines were maintained in hE8, whereas NANOG-overexpressing and GERALT lines were maintained in hE7, which lacked TGF*β*1. Cultures were passaged on Monday mornings and Thursday afternoons, with additional medium changes on Tuesdays and Fridays. EDTA passaging and 24 h thiazovivin supplementation were otherwise unchanged. Complete formulations, including the FGF2 variant used in each experiment, are provided in Supplementary Table 2.

For TGF*β*-withdrawal and pathway-inhibition assays, cells were cultured in commercial Essential 6 (Thermo Fisher Scientific, A1516401) supplemented with 100 ng mL^−1^ FGF2-145aa (Qkine, Qk025). This formulation is referred to as E7 throughout the corresponding Results and figures. Addition of 2 ng mL^−1^ TGF*β*1 (Qkine, Qk010) generated the TGF*β*-replete condition, referred to as E8, whereas addition of 10 µM SB431542 (Selleck Chemicals, S1067) generated the pathway-inhibited condition, referred to as E7+SB. Thus, in these experiments, “E8” denotes the E6-based FGF2/TGF*β*1 formulation rather than the ready-to-use commercial Essential 8 medium used for routine hiPSC maintenance.

Acute pathway-inhibition experiments compared ET, ETN_1_ and ETN_2_ at baseline and after 2, 24 and 48 h of treatment. Cells were seeded at 1 × 10^5^ cells per well in Geltrex-coated 12-well plates, and treatments were staggered to permit simultaneous collection of all time points. Longer-term assays compared the indicated lines after 4 or 8 d in E8, E7 or E7+SB; cells were seeded at 7 × 10^4^ cells per well and medium was replaced daily. For serial culture without exogenous TGF*β*, ET cells were analysed at passage 3 and ETN_2_ cells at passages 3, 5 and 10.

### Circuit design and genome editing

The renewal cassette comprised CAG-driven *NANOG* coexpressed with an mCherry reporter *via* a 2A peptide at *CLYBL*. Tet-On-3G was expressed from *hROSA26*, and dox-responsive lineage-driver cassettes were targeted to *AAVS1*, as previously described^16^. Anti-*NANOG* miR-F hairpins (based on a modified miR-E backbone carrying an A-to-G substitution that eliminates a basal-stem bulge^29,49^) were positioned in the driver 3’ UTR, in a separate bidirectional transcript, or within an intron; anti-CD151 and anti-RLuc hairpins served as controls. Construct descriptions and Addgene accession numbers are provided in Supplementary Table 3, with assembly details in the Supplementary Note 2. All plasmids were verified by whole-plasmid sequencing (Plasmidsaurus or Eurofins Genomics).

*hROSA26* targeting used paired Cas9 nickases, *CLYBL* targeting used paired TALENs, and *AAVS1* targeting used paired zinc-finger nucleases^50^ (Supplementary Table 3). In all cases, donor and nuclease plasmids were delivered at a 2:1:1 mass ratio. For *CLYBL* targeting and *AAVS1* targeting of the GERALT HFHR cassette, 1 × 10^6^ Accutase-dissociated cells were nucleofected with 10 µg total DNA using the P3 Primary Cell 4D-Nucleofector X Kit (Lonza, V4XP-3032; program CA137). Cells were recovered in CEPT-supplemented E8 for *CLYBL* targeting or hE8 for GERALT HFHR^51^; the CEPT formulation is provided in Supplementary Table 2. All other *hROSA26* and *AAVS1* targeting was performed by lipofection. Cells were seeded at 2 × 10^5^ per well of a six-well plate and transfected 24 h later for 4 h in Opti-MEM (Thermo Fisher Scientific, 31985062) with 2 µL Lipofectamine Stem (Thermo Fisher Scientific, STEM00015) and 2 µg total DNA per well.

Edited cells were selected for 5 d with 4 µg mL^−1^ blasticidin S (Sigma-Aldrich, 15205) for *hROSA26*, 50 µg mL^−1^ Geneticin (Thermo Fisher Scientific, 10131035) for *CLYBL*, or 0.5 µg mL^−1^ puromycin (Thermo Fisher Scientific, A1113803) for *AAVS1*. Thiazovivin (2 µM) was included for 48 h during the period of greatest selection-associated cell loss. Individual antibiotic-resistant colonies were manually isolated and expanded as pseudoclones. Targeted integration was assessed by junctional, locus and donor-backbone PCRs according to the strategy shown in Extended Data Fig. 1b; the resulting genotypes are summarized in Supplementary Table 1. Genomic DNA was extracted using the Monarch Genomic DNA Purification Kit (NEB, T3010L) and eluted in nuclease-free water. Genotyping reactions contained 1× LongAmp buffer, 2 U LongAmp Taq (New England Biolabs, M0323), 250 nM each primer, 2% DMSO, 300 µM of each dNTP and 100 ng genomic DNA in a 10 µL reaction and were amplified for 35 cycles. Primer sequences, annealing and extension conditions, and expected amplicon sizes are provided in Supplementary Table 4.

### Genomic copy-number analysis

Genomic copy-number analyses were performed between passages 5 and 10 after clonal isolation, as summarized in Supplementary Table 1. ETN_2_ was analysed by chromosomal microarray using the KaryoStat+ Assay service (Life Technologies Corporation/Thermo Fisher Scientific; sample ID KS-14237) on the CytoScan HT-CMA 96F platform (SKU 906025; approximately 1.1 million probes) using 100 ng genomic DNA. Copy-number data were mapped to GRCh38 and normalized to the platform reference set. Reported detection thresholds were >2 Mb for chromosomal gains and >1 Mb for losses.

SNP-array profiling of WT and the indicated engineered clones was performed by Life & Brain using the Illumina Infinium Global Screening Array v3.0 (GSAMD-24v3-0; >700,000 markers), following the analysis workflow described previously^30^. Genomic positions were based on GRCh37/hg19. Data were processed using GenomeStudio 2.0.5 and cnvPartition 3.2.1 with a minimum of six probes per call. Downstream filtering retained calls with confidence >75 and size >350 kb, or >100 kb within previously defined hPSC genomic-instability hotspots^35^. Analysis scripts, including hotspot definitions and downstream filtering criteria, are available as described under Code availability.

### Inducible miR-F validation

TRE-driven EGFP cassettes bearing anti-CD151, anti-CD46 or anti-RLuc miR-F hairpins in the 3’ UTR were targeted to *AAVS1* in ET cells. Antibiotic-selected pools were analyzed without clonal isolation. Cells were maintained in E8 with or without 1 µg mL^−1^ dox (doxycycline hyclate; Sigma-Aldrich, D9891), with daily medium replacement. Surface-protein expression was evaluated after 5 d of dox exposure. Anti-*NANOG* miR-F was evaluated analogously in ETN_2_ pools. Sequences of miR-F hairpins are provided in Supplementary Table 5.

### Myogenic forward programming

Accutase-dissociated engineered hiPSCs were seeded on Geltrex at 2 × 10^5^ cells per well of a 12-well plate with 2 µM thiazovivin for 24 h. Dox was then added at 1 µg mL^−1^, defining differentiation day 0.

Forward programming was performed in myogenic medium, hE7 or hE6, as defined in Supplementary Table 2. The myogenic medium followed the ioSkeletal Myocytes culture protocol (bit.bio) and was similar in composition to that described by Pawlowski et al.^16^. Relative to myogenic medium, hE7 lacked retinoic acid and CHIR99021, whereas hE6 additionally lacked FGF2. Media were replaced daily.

### Directed myogenic differentiation

WTC11 hiPSCs (Coriell Institute, GM25256) were seeded at 2 × 10^5^ cells per well of a 6-well plate on 167 µg mL^−1^ Matrigel (Corning, 356231) in mTeSR (STEMCELL Technologies, 85850) supplemented with 10 µM Y-27632. Differentiation was performed for 28 d using a protocol adapted from Smith et al.^32^. Complete medium compositions, final supplement concentrations and reagent identifiers are provided in Supplementary Table 2. CHIR99021 and LDN193189 were present from day 0, and FGF2 was added on day 2. On day 5, CHIR99021 and ITS were withdrawn, while KnockOut Serum Replacement, IGF1 and HGF were added. FGF2, HGF and LDN193189 were withdrawn on day 7, and HGF was restored on day 11. Thus, from day 11 through day 28, cultures were maintained in DMEM/F12 supplemented only with NEAA, KnockOut Serum Replacement, IGF1 and HGF, with daily medium replacement.

On day 28, cells were dissociated with TrypLE (Gibco 12563011) for approximately 15 min at 37 °C, filtered through a 40-µm strainer and centrifuged at 300 *g* for 5 min. Cells were replated in skeletal muscle growth medium (SKGM; Supplementary Table 2) containing 10 µM Y-27632 onto Matrigel-coated 10-cm dishes (100 µg mL^−1^), at a ratio of two original 6-well plate wells per dish. Cultures were maintained until day 35, after which NGFR-positive cells were isolated by FACS and expanded in SKGM. For terminal differentiation, derived myoblasts were seeded on Geltrex at 2 × 10^5^ cells per well of a 6-well plate and switched after 24 h to myoblast fusion medium (Supplementary Table 2) for 4 d, with daily medium replacement.

### Primary human myoblasts

Primary human skeletal muscle myoblasts used for the single-cell benchmark (HSMM; Lonza, CC-2580; lot 23TL023429) were derived from a 32-year-old male donor, supplied cryopreserved at passage 2 and used experimentally at passage 4. Cells were expanded in SkGM-2 (Supplementary Table 2) at 37 °C and 5% CO_2_, maintained below 70% confluence and passaged using TrypLE Express (Gibco, 12604013). At 50–60% confluence, cultures were switched to HSMM fusion medium (Supplementary Table 2) for 4 d, with medium replacement every other day.

Primary human skeletal muscle-derived cells used for the independent EMT benchmark were obtained from Cook MyoSite (SK-1111; lot P01318-35M.3) and were derived from the vastus lateralis of a 35-year-old male Caucasian donor. Cells were designated passage 0 upon receipt (supplier passage unknown) and used for EMT generation at passage 4. Before EMT generation, cells were expanded in SKGM at 37 °C and 5% CO_2_, maintained below approximately 80% confluence and passaged using TrypLE.

### Neuronal forward programming

Three GERALT NGN2 clones were compared with WT, E and ET lines independently transduced with a dox-inducible *NEUROG2/NGN2*/puromycin-resistance lentiviral system^14^. Lentivirus was produced in HEK293T cells using the third-generation packaging plasmids pRSV-Rev (Addgene, 12253), pMDLg/pRRE (Addgene, 12251) and pMD2.G (Addgene, 12259), together with the transfer vectors pTet-O-Ngn2-puro (Addgene, 52047) and FUW-M2rtTA (Addgene, 20342). For each lentiviral preparation, the volume of viral supernatant used for transduction was determined empirically to achieve efficient transduction.

GERALT NGN2 cells were seeded on Geltrex at 7.5 × 10^4^ cells per well of a 12-well plate and induced 24 h later with 1 µg mL^−1^ dox. Lentiviral controls were seeded on Geltrex at 1.75 × 10^5^ cells per well of a 12-well plate in StemFlex (Gibco, A3349401) supplemented with 10 µM Y-27632 (Sigma-Aldrich, SCM075), and simultaneously co-transduced with pTet-O-Ngn2-puro and FUW-M2rtTA lentivirus. Dox was added 16–18 h later at 2 µg mL^−1^; this defined differentiation day 0. For immunofluorescence, both platforms were seeded at 8.7 × 10^4^ cells per well on 12-mm coverslips in 24-well plates.

Both platforms were differentiated in hE6 or N2 medium (Supplementary Table 2). For lentiviral controls, puromycin selection was initiated on day 1, after 24 h of dox exposure, and maintained at 2 µg mL^−1^ (Sigma-Aldrich, P8833). Media were replaced daily.

### Hepatic forward programming

Induction of the polycistronic HFHR cassette was assessed in undifferentiated GERALT HFHR cells after 24 h of treatment with 1 µg mL^−1^ dox in hE7.

Development of the GERALT-optimized protocol is described in Supplementary Note 1. GERALT HFHR and FSPS13B OPTi-OX HFHR cells were differentiated using an empirically adapted version of the reference HFHR forward-programming protocol^31^. For the final GERALT-optimized protocol, cells were seeded on Geltrex on day −1 at 9 × 10^4^ cells per well of a 12-well plate in hE7 supplemented with 2 µM thiazovivin. Thiazovivin was withdrawn on day 0. On day 1, HFHR expression was induced for 24 h with 1 µg mL^−1^ dox in hE6 at 37 °C and 5% O_2_. On day 2, cells were washed with PBS and transferred to HLI containing 0.5% lipid concentrate (Supplementary Table 2). Medium was replaced daily through day 4 and every other day thereafter. On day 6, cultures were transferred to 39 °C under normoxia (20% O_2_, 5% CO_2_) and maintained to day 20.

For benchmarking against the reference OPTi-OX protocol^31^, FSPS13B OPTi-OX HFHR cells were seeded on Geltrex on day −1 at 7.5 × 10^4^ cells per well of a 12-well plate in hE8 supplemented with 2 µM thiazovivin. Thiazovivin was withdrawn on day 0, and HFHR induction was initiated on day 1 and maintained through day 10, with cultures transferred to HLI containing 2% lipid concentrate from day 2. Cells were maintained at 37 °C under 5% O_2_ until transfer to 39 °C under normoxia on day 15.

### Suspension expansion and differentiation

GERALT MYOD1 hiPSCs were dissociated with Accutase and seeded at 3 × 10^5^ cells mL^−1^ in 125-mL vented Erlenmeyer flasks (Corning, 734-1885) containing 20 mL hE7 (Supplementary Table 2). Cultures were maintained at 70 rpm on a 19-mm-orbit CO_2_-resistant orbital shaker (Thermo Fisher Scientific, 88881104) at 37 °C and 5% CO_2_. Cells were treated with 2 µM thiazovivin for 24 h before transfer from adherent culture and during the first 24 h after suspension seeding. Medium was then completely replaced and subsequently exchanged every 2 d.

Cultures were passaged every 4–5 d. Aggregates were incubated in 0.5 mM EDTA/PBS for 10 min at 37 °C with gentle pipetting at 5-min intervals, diluted with DMEM/F-12 and centrifuged at 100*g* for 5 min. The resulting small aggregates were reseeded at 1.5–2 × 10^5^ cells mL^−1^ in hE7 containing 2 µM thiazovivin. For quantitative cell-yield measurements, aggregates were dissociated to single cells with Accutase and counted manually using a hemo-cytometer. Pluripotency was assessed after 10 passages.

For suspension forward programming, aggregates were cultured for 3 d after reseeding and then transferred to myogenic medium (Supplementary Table 2) containing 1 µg mL^−1^ dox. Cultures remained under continuous 70-rpm agitation and medium was replaced daily for 7 days of differentiation.

### Engineered muscle tissue generation

Engineered muscle tissues (EMTs) were generated using the Mantarray platform (Curi Bio) from GERALT MYOD1 hiPSCs induced for 24 h before casting (dox pre), GERALT MYOD1 hiPSCs induced after casting (dox post), or WTC11-derived myoblasts generated by directed differentiation as described above^32^. A second independent WTC11 differentiation was used for the additional benchmark. Primary human skeletal muscle-derived cells were expanded according to the manufacturer’s instructions.

Mantarray post lattices were treated sequentially with 0.1% polyethylenimine (Sigma-Aldrich, P3143) and 0.01% glutaraldehyde (Thermo Fisher Scientific, BP25481). HS-27A human bone-marrow stromal cells (ATCC, CRL-2496) were used as supporting cells and expanded in RPMI-1640 supplemented with 10% FBS. Each EMT contained 1 × 10^6^ hiPSCs or myogenic cells and 1 × 10^5^ HS-27A cells in a 150-µL fibrin hydrogel containing 3.33 mg mL^−1^ fibrinogen (Sigma-Aldrich, F8630), 2 U mL^−1^ thrombin (Sigma-Aldrich, T4648) and 20% (v/v) Matrigel in high-glucose DMEM. Tissues were polymerized for 80 min at 37 °C.

Immediately after casting, dox-post GERALT tissues were cultured for 24 h in EMT post-casting medium for direct hiPSCs, whereas dox-pre GERALT and WTC11-derived myoblast tissues received the corresponding pre-induced post-casting medium (Supplementary Table 2). Thiazovivin was then withdrawn and tissues were maintained in EMT differentiation medium (Supplementary Table 2), with dox maintained throughout GERALT tissue culture. Medium was replaced every other day.

### RNA extraction and RT-qPCR

RNA from monolayer cultures was extracted using Quick-RNA Miniprep (Zymo Research, R1055). Individual EMTs were processed using RNeasy Plus Micro (QIAGEN, 74034): tissues were digested for 10 min at 56 °C with agitation in 100 µL DPBS containing 2 mg mL^−1^ proteinase K (NEB, P8107S), lysed with 350 µL RLT Plus containing 1% 2-mercaptoethanol (Sigma-Aldrich, M6250), passed through a gDNA Eliminator column, mixed with 250 µL ethanol and processed on MinElute columns.

cDNA was generated with the High-Capacity cDNA Reverse Transcription Kit (Thermo Fisher Scientific, 4374967), using 400 ng total RNA per reverse-transcription reaction. Quantitative PCR used PowerUp SYBR Green Master Mix (Thermo Fisher Scientific, A25742), cDNA corresponding to 10 ng input RNA and 0.3 µM each primer in a final reaction volume of 10 µL on a QuantStudio 6 Flex for 40 cycles. Each sample was analysed in technical duplicate; technical-duplicate *C*_*t*_ values were averaged. No-reverse-transcription controls prepared from pooled RNA and no-template controls containing nuclease-free water were included. Amplicon specificity and primer efficiency were assessed by dissociation and standard curves, respectively, and undetected reactions were set to *C*_*t*_ = 40. Primer sequences and target specificity are provided in Supplementary Table 6.

Expression was normalized to the indicated reference gene or genes, with Δ*C*_*t*_ = *C*_*t*,target_ − *C*_*t*,reference_. Where multiple housekeeping genes were used together, the reference *C*_*t*_ was the geometric mean of their *C*_*t*_ values. Reference-normalized expression was calculated as 100 × 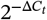 and plotted as a percentage of reference-gene expression. Heatmaps display gene-wise z-scores calculated from reference-normalized expression after replicate averaging, without prior log transformation.

Because relative-expression values derived from 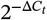 can span several orders of magnitude and show unequal variance, statistical analyses of RT-qPCR data were performed on the corresponding Δ*C*_*t*_ values, which better approximate the assumptions of parametric tests^52^. Where multiple differentiations were summarized into a single experimental unit, Δ*C*_*t*_ values were averaged at that level before statistical analysis.

### Flow cytometry

For live EGFP, mCherry and TTN-mEGFP measurements, adherent cells and, where applicable, suspension aggregates were dissociated to single cells with 0.5% trypsin (Euroclone, ECB3051D), viability-stained with eFluor 450 (Thermo Fisher Scientific, 65-0863-14; 1 µL mL^−1^) for 10 min at room temperature and washed in FACS buffer (PBS containing 5% FBS).

For surface CD151/CD46 measurements, trypsin-dissociated cells were viability-stained and incubated with the corresponding APC-conjugated antibody for 1 h at room temperature. Target fluorescence was evaluated separately within induced EGFP-positive and EGFP-negative populations, with matching isotype and uninduced controls.

For intracellular NANOG/OCT4 measurements, cells were dissociated to single cells with TrypLE and viability-stained before fixation in 4% paraformaldehyde for 10 min at room temperature. Cells were permeabilized with 0.1% Triton X-100 for 30 min and stained for 1 h at room temperature with Alexa Fluor 647-NANOG and PE-OCT4 in FACS buffer containing 0.75% saponin (Sigma-Aldrich, S7900). The same intracellular-staining procedure was used for anti-*NANOG* miR-F pool assays. Antibodies, isotype controls and working concentrations are listed in Supplementary Table 7.

Samples were acquired on a BD FACSVerse (BD Biosciences), except for the mCherry expression assay, which was acquired on a Sony SH800S cell sorter. At least 10,000 events were acquired per sample. Compensation used single-stained controls.

Data were analyzed in FlowJo 10.10.0. Debris and doublets were excluded by FSC/SSC and singlet gating, respectively, followed by exclusion of eFluor 450-positive non-viable cells. Reporter- or antibody-positive populations were defined relative to the corresponding uninduced, unstained or isotype controls, as appropriate.

### Western blotting

Cells were lysed on ice for 10 min in PBS containing 1% Triton X-100 and protease inhibitors (Roche, 11697498001), and lysates were clarified at 13,000*g* for 10 min at 4 °C. Protein concentration was determined by BCA assay (Thermo Fisher Scientific, 23227). Twenty micrograms of protein per sample were heated in 1× LDS sample buffer containing *β*-mercaptoethanol for 10 min at 95 °C and separated on 4–20% SDS–PAGE gels (Bio-Rad, 4561094).

Proteins were wet-transferred to 0.45-µm nitrocellulose membranes (Amersham Protran; Cytiva, 10600002) at 100 V for 1 h at 4 °C. Membranes were blocked in TBST containing 5% BSA for 1 h at room temperature, incubated with NANOG and VCL primary antibodies overnight at 4 °C and with HRP-conjugated secondary antibodies for 1 h at room temperature. Signals were developed using Clarity ECL (Bio-Rad, 170-5061) and acquired on a Chemi-Doc MP using the Auto Optimal exposure setting. Antibody details are provided in Supplementary Table 7.

Band intensities were quantified in ImageJ after background subtraction. NANOG signal was normalized to VCL within each lane and expressed relative to TGF*β*-containing baseline medium.

### Microscopy

Myogenic coverslip cultures were fixed in 4% paraformaldehyde for 10 min, permeabilized in 0.1% Triton X-100 for 20 min and blocked in 5% BSA for 1 h at room temperature. Primary antibodies diluted in 1% BSA were applied overnight at 4 °C, followed by fluorescent secondary antibodies for 1 h at room temperature. Nuclei were counterstained with DAPI (Thermo Fisher Scientific, 62248), and coverslips were mounted using ProLong Glass Antifade Mountant (Thermo Fisher Scientific, P36984). Direct mCherry and TTN-mEGFP fluorescence and antibody-based MYOD1 and ACTN2 detection were used as appropriate. Antibody identities and working concentrations or dilutions are provided in Supplementary Table 7.

Neuronal cultures were fixed in 4% paraformaldehyde/4% sucrose in PBS, pH 7.4, for 15 min at room temperature and washed in PBS containing 1% BSA and 0.1% Tween-20 (PBST). Cells were permeabilized for 10 min in PBS containing 3% BSA, 0.1% Triton X-100 and 1% goat or donkey serum, followed by blocking in 5% BSA for 30 min. Primary antibodies diluted in PBS containing 1% BSA were applied for 1 h, followed by three washes in PBST. Cells were then incubated with Alexa Fluor-conjugated secondary antibodies diluted in PBS containing 1% BSA for 1 h and stained with DAPI in PBS containing 1% BSA for 10 min. Coverslips were mounted using ProLong Gold Antifade Mountant (Thermo Fisher Scientific, P36934).

For hepatic immunofluorescence, cultures were fixed in 4% paraformaldehyde for 10 min, permeabilized with 0.1% Triton X-100 for 10 min and blocked in 3% BSA containing 1% goat serum for 1 h. Primary antibodies were applied overnight at 4 °C, followed by fluorescent secondary antibodies for 1 h at room temperature and DAPI staining for 2 min. Cultures were grown on coverslips or *µ*-Slide 8 Well imaging chambers (ibidi, 80806). For lipid staining, fixed hepatic cultures were incubated with 5 µM BODIPY 493/503 (MedChemExpress, HY-W090090) for 30 min at room temperature. Samples were washed three times and counterstained with DAPI for 5 min and washed again.

Engineered muscle tissues were fixed in 4% paraformaldehyde for 20 min, cryoprotected overnight at 4 °C in 30% sucrose (Sigma-Aldrich, S0389), embedded in OCT compound (Bio-Optica, 05-9801), cryosectioned at 12 µm using a Leica CM1950 and stored at −20 °C. Longitudinal sections were obtained along the long axis of the tissue. Sections were permeabilized in 0.1% Triton X-100 for 20 min and blocked in 5% BSA for 1 h; primary- and secondary-antibody staining and mounting followed the myogenic procedure.

Live phase-contrast and reporter images were acquired in culture medium on a ZEISS Observer.Z1 equipped with ApoTome using AxioVision. Myogenic cultures were imaged using a ZEISS Axiovert 5 with Labscope or a ZEISS ELYRA 7; hepatic cultures were imaged using a ZEISS ELYRA 7, and neuronal cultures using a Leica TCS SP8 with LAS X. Whole-EMT images were acquired on a Leica THUNDER using a 10× objective and reconstructed by tile stitching with computational clearing in LAS X. High-resolution EMT imaging was performed on a ZEISS ELYRA 7 using either a 63× objective in structured illumination microscopy (SIM) mode or a 40× objective for conventional fluorescence imaging. SIM images were reconstructed in ZEN Black (AIMApplication version 16.0.19.306) using SIM2 reconstruction.

### Image quantification

Quantitative image analysis was performed in Fiji (ImageJ 1.54p)^53^. Nuclear segmentation was performed on the DAPI channel after Gaussian smoothing and automated intensity thresholding. Binary nuclear regions of interest (ROIs) were filtered by area and circularity to exclude debris, non-nuclear objects and improperly segmented nuclear clusters. For SOX2 and NANOG quantification, local background subtraction was applied to the fluorescence channel and mean fluorescence intensity was measured within each DAPI-defined nuclear ROI. Cells were classified as SOX2- or NANOG-positive using an empirically defined intensity threshold separating specific signal from background, and the fraction of SOX2- or NANOG-positive cells was calculated relative to the total number of segmented nuclei in each field. Single-cell fluorescence values were additionally exported for analysis of population-level intensity distributions.

For TUBB3 and MAP2 area measurements, signal-positive regions were segmented by automated intensity thresholding and converted to binary masks. The area occupied by each segmented signal was measured within each field of view. The fluorescence-positive area was then normalized to DAPI-defined nuclear number.

For all neuronal image analyses, three random fields were acquired per sample using consistent acquisition settings across compared conditions. Field-level measurements were averaged per sample.

### Multielectrode-array recordings

On day 3 of forward programming, GERALT NGN2 cells were dissociated with TrypLE and seeded at 1 × 10^5^ cells per well in poly-D-lysine- (Gibco, A3890401) and laminin-coated (Gibco, 23017015) CytoView 48-well MEA plates (Axion Biosystems). Cells were initially plated in a 6 µL droplet of hE6 supplemented with 1 µg mL^−1^ dox and 10 µM Y-27632 to promote attachment over the recording electrodes. After approximately 1 h, 400 µL of culture medium was added. Subsequently, 300 µL was replaced every 3 d, retaining approximately 100 µL per well during medium exchange. Dox was maintained throughout the experiment.

One-minute recordings were acquired from day 14 at 37 °C and 5% CO_2_ using a Maestro MEA system (Axion Biosystems) and AxIS 2.5 software, following temperature and CO_2_ equilibration. Signals were sampled at 12.5 kHz per channel with median referencing, a 200-Hz digital high-pass IIR filter and a 3-kHz digital low-pass Kaiser-window filter. Electrodes detecting at least five spikes per minute were considered active, and weighted mean firing rate was calculated across active electrodes. Bursts were defined as at least five spikes with a maximum inter-spike interval of 100 ms. Recordings were performed in two independent differentiation experiments.

### Albumin secretion and CYP3A4 activity

Conditioned medium from day-20 hepatic cultures was collected 48 h after the last medium change and albumin concentration was measured using the RayBio Human Albumin ELISA Kit (RayBiotech, ELH-Albumin) according to the manufacturer’s instructions, with samples diluted to fall within the calibration range. Concentrations were determined after blank subtraction and normalization for sample dilution and total cell number.

CYP3A4 activity was measured at day 20 using the P450-Glo CYP3A4 Assay with Luciferin-IPA (Promega, V9001). Cultures were incubated with 3 µM Luciferin-IPA for 1 h at 39 °C. Aliquots of conditioned medium were combined 1:1 with Luciferin Detection Reagent, incubated for 20 min at room temperature and luminescence measured using a GloMax luminometer. Signals were background-subtracted using the corresponding cell-free HLI condition containing Luciferin-IPA and normalized to sampled volume and total cell number. Cell numbers for both assays were determined using a Countess 3 Automated Cell Counter (Thermo Fisher Scientific, AMQAX2000), and results were expressed per 10^6^ cells.

### Engineered muscle tissue contractility

EMT contractility was assessed on day 21 using the Mantarray platform at 37 °C. Magnetic post displacement was acquired at 100 Hz and converted to force using the manufacturer’s calibration. Tissues were stimulated with biphasic 5-ms, 100-mA pulses. Force–frequency responses were measured at 1, 2, 3, 5, 10, 20, 30, 40 and 50 Hz, with 8-s rest intervals; stimulation was applied for 5 s at 1 Hz and for 3 s at all higher frequencies. Tetanic force was additionally assessed using a 3-s, 50-Hz stimulation train.

Fatigue was assessed using 10 consecutive 3-s, 50-Hz stimulation trains separated by 3-s rest intervals. Force for each train was normalized to that of the first train within the same tissue.

Raw force–time traces were analyzed using EMCAP v1.0.1, an open-source Python-based peak-detection and contractility-analysis pipeline, with protocol-timing definitions adapted to the stimulation sequences used here. The analysis code is archived as described in the Code availability section. As all EMTs were generated using matched input cell numbers, force was reported in absolute units without normalization to tissue cross-sectional area, allowing comparison of total contractile output from equivalent starting cell inputs.

### Bulk RNA sequencing

Sample identities, experimental conditions, sequencing metrics and quality-control information are detailed in Supplementary Table 8. RNA was extracted as above and assessed with RNA ScreenTape (Agilent, 5067-5576), with RNA integrity > 9. Poly(A)-selected stranded libraries were prepared from 500 ng RNA using Illumina Stranded mRNA Prep, Ligation (Illumina, 20040534; protocol 1000000124518 v02) with Illumina-compatible unique dual indexes (IDT for Illumina RNA UD Indexes Set A, Ligation, 20040553). Libraries were checked with D1000/D5000 ScreenTape (Agilent, 5067-5582/5067-5588), quantified by Qubit (Life Technologies, Q33230) and pooled according to their molar concentrations to target 35–40 million read pairs per sample.

NEGEDIA sequenced the pluripotency and myogenic experiment libraries on DNBSEQ-T7 and DNBSEQ-G400 instruments, respectively, using paired-end 100-bp reads. Sequencing libraries were converted to DNBSEQ-compatible libraries using the MGIEasy Universal Library Conversion Kit (MGI Tech, 1000004155), following the manufacturer’s instructions.

Sequencing reads were aligned using the alignment component of our dockerized custom Bulk RNA-seq pipeline, with STAR v 2.7.11b^54^ against a custom GRCh38/Ensembl release 112 reference including *NANOG*_exo_, Tet-On 3G, transgenic *MYOD1* and other construct-associated sequences. Additional custom reference features and annotations are archived on Zenodo (doi:10.5281/zenodo.21471691). Reads underwent trimming before alignment with Trim Galore v2.3.0 embedded in the pipeline. Gene-level counts were obtained from the stranded-count column of ReadsPerGene.out.tab.

Analyses used R 4.4.2; Ensembl identifiers were annotated with symbols and biotypes through BioMart. Genes were retained if they had at least three counts in at least as many samples as the smallest biological group included in the relevant comparison. Analyses were primarily restricted to protein-coding genes, with relevant custom transgene features retained.

DESeq2 1.46.0^55^ provided median-of-ratios normalization and variance-stabilized expression for exploratory analyses and visualization. Dataset quality was assessed from library sizes, gene-detection rates, mitochondrial-transcript abundance and transcript-biotype composition. Sample relationships were evaluated by Pearson-correlation clustering and PCA of the 1,000 most variable genes, without additional batch correction. TPM values, where used for display, were calculated using non-overlapping exon lengths and were not used for differential expression.

Myogenic pairwise comparisons used DESeq2 Wald tests with design `~group`, including pooled anti-*NANOG* samples versus ETM and BID versus INT. The pluripotency model used design `~cell_type * treatment * time`, referenced to ET, baseline medium and 24 h; within-genotype models used `~treatment * time`. Wald tests evaluated the stated main effects and interactions. Likelihood-ratio tests compared the full factorial model with `~cell_type * treatment` to test all time-related terms jointly, and with a model containing all main effects and two-way interactions to test the three-way interaction. Within each genotype, treatment-by-time interactions were tested by comparing `~treatment * time with ~treatment + time`. No replicate or block covariate was included in these bulk differential-expression models. Interaction effect sizes were obtained from the corresponding Wald fits. Differentially expressed genes required Benjamini–Hochberg-adjusted *P* < 0.05 and absolute log_2_ fold change > 1.

GO Biological Process over-representation analyses and ranked enrichment used clusterProfiler 4.14.6^56^ and `org.Hs.eg.db`. Over-representation analyses were performed separately for up- and downregulated genes using expressed genes. Ranked analyses used DESeq2 log_2_ fold changes with `gseGO` using the `fgsea` algorithm, 10,000 simple permutations and a minimum set size of five genes. Terms with Benjamini–Hochberg-adjusted *P* < 0.05 were considered significantly enriched. Genes with the top 1% most negative PC1 loadings were tested for functional enrichment. Curated signatures covering pluripotency, lineage, myogenesis, cell cycle, metabolism and muscle fiber type are listed in Supplementary Table 10.

Heatmaps were generated from variance-stabilized or standardized expression values. Figures were generated using pheatmap v1.0.13, ggplot2 v4.0.2, EnhancedVolcano v1.24.0 and enrichplot v1.26.6. Custom downstream R workflows are listed in the Code availability section.

### Single-cell RNA sequencing

#### 10x Genomics dataset

Twelve samples represented ET and ETN_2_ in E8 and after E7 adaptation (passages 3 and 10, respectively), with three independent cultures per group. Cells were dissociated using TrypLE Express, pelleted at 300*g* for 5 min, washed in PBS containing 1% BSA (Miltenyi Biotec, 130-091-376) and counted by trypan-blue exclusion (Sigma-Aldrich, T8154). Approximately 1.5 × 10^6^ cells per sample were labelled with CellPlex oligos (10x Genomics, 1000261/1000262; CG000391 Rev. A), washed three times and pooled equally. Approximately 26,400 pooled cells were loaded onto a Chromium Next GEM Chip G.

Gene-expression and CellPlex libraries were prepared using the Chromium 3^′^ v3.1 multiplexing workflow (10x Genomics, 1000269; CG000388 Rev. B), assessed using D5000 ScreenTape and pooled at a 6:1 gene-expression:CellPlex molar ratio. Sequencing used NextSeq 1000/P2 XLEAP-SBS (Illumina, 20100987), with read/index cycles 28/10/10/90. Cell Ranger 9 jointly processed and demultiplexed gene-expression and CellPlex reads using multi, followed by aggr, within Docker image hedgelab/rstudio-hedgelab:iPS2seq_new_cellranger9. mkref generated the GRCh38/Ensembl 112 reference supplemented with *NANOG*_exo_, Tet-On 3G, and mCherry. Sample-level sequencing and post-filter quality-control metrics are reported in Supplementary Table 9; custom reference files, processing configurations and analysis code are listed in the Data and Code availability sections.

The filtered gene-by-cell count matrix was imported into Seurat 5.3.0^57^ in R 4.4.2. Cell filters were > 300 and < 7,000 detected genes, > 100 RNA counts, 1–10% mitochondrial transcripts and > 4% ribosomal transcripts. Analyses were restricted to protein-coding genes while retaining the relevant custom transgene features. Filtered counts and metadata were converted to a Monocle 3 v1.3.4^58^ `cell_data_set`. Preprocessing used 30 dimensions; UMAP used minimum distance 0.2, and `cluster_cells` used *k* = 15 and resolution 2×10^−4^. Markers detected in at least 10% of cells were ranked by pseudo-*R*^2^ using `top_markers`, and cluster labels and embeddings were transferred to Seurat for downstream analyses.

Cell-level differential expression was assessed on log normalized counts using Seurat `FindAllMarkers` and `FindMarkers` Wilcoxon rank-sum tests, with pairwise and one-versus-rest comparisons based on the Monocle cluster assignments. Differentially expressed genes required FDR-adjusted *P* ≤ 0.01 and absolute log_2_ fold change ≥ 1.

Pseudobulk profiles were generated from raw counts using `AggregateExpression`, either by sample or by sample and cluster. Genes with fewer than 10 total counts in the fitted dataset were excluded. Condition-level models included cell line, medium and their interaction, referenced to ET/E7. For cluster-level analyses, observed sample–cluster combinations were retained without an additional minimum-cell threshold and absent combinations were omitted. Within each genotype, DESeq2 Wald tests used design `~monocle_clusters`, without sample, replicate or medium covariates. Contrasts compared Undiff versus Neuro clusters in ET and TGF*β*-high versus ZIC1-positive clusters in ETN_2_. Pseudobulk significance required adjusted *P* < 0.05 and absolute log_2_ fold change ≥ 1. Sample-level pseudobulk PCA used counts summed across clusters, DESeq2 median-of-ratios normalization, variance stabilization (`blind = TRUE`) and the 500 most variable transformed genes.

GO Biological Process over-representation analysis used `enrichGO` on Neuro-upregulated genes identified by cell-level Wilcoxon tests, with unique Entrez IDs mapped from all retained genes in the comparison table as the background universe. Separately, pseudobulk GSEA used `gseGO` with the `fgsea` backend and 10,000 simple permutations, testing gene sets containing 5–500 genes. Genes with non-missing adjusted *P* values and effect estimates were ranked by decreasing DESeq2 log_2_ fold change. For ranked analyses, duplicate Entrez IDs were resolved by retaining the entry with the largest absolute fold change. Both analyses used Benjamini–Hochberg correction.

Cell-cycle scores were calculated using Seurat `CellCycleScoring` with the S-phase and G2/M gene sets in `cc.genes` and 20 expression bins. Cells were assigned to G1, S or G2/M using the default scoring-based classification. No cell-cycle regression was applied.

Reference-label transfer used the pluripotent-state dataset of Balmas et al.^27^ (E-MTAB-14065), excluding the Neuro query cluster. Variable features were selected using the `vst` method with 8,000 features. Reference labels were transferred using `FindTransferAnchors (normalization.method = “LogNormalize”`) and `TransferData`, using dimensions 1–30 and the `originalexp` assays. Transferred labels comprised iPSC-N, iPSC-P, iPSC-1, iPSC-2 and iPSC-M. Predicted labels were retained without a confidence threshold or explicit unassigned category. Label fractions were calculated as the number of cells assigned to each reference label divided by all cells in the corresponding query cluster, pooled across samples.

Differential abundance was assessed using MiloR 2.2.0^28^ and SingleCellExperiment 1.28.1. Neighborhoods were defined using 30 dimensions, *k* = 30, a sampling proportion of 0.1 and refinement enabled, and were annotated according to their predominant Monocle cluster. Counts per biological sample were analyzed using `~0 + condition` for ET E7 versus ET E8, ETN_2_ E8 versus ET E8, and ETN_2_ E7 versus ET E8; the ETN_2_-only E7-versus-E8 analysis used `~medium`. Positive log_2_ fold changes indicated enrichment in the first condition. No replicate-blocking term was included. Spatial FDR used the default `k-distance` weighting, with spatial FDR < 0.05 defining significant neighborhoods.

For cluster-level summaries, the fraction of significant neighborhoods was calculated relative to all neighborhoods assigned to that cluster. Descriptive enrichment flags required a median log_2_ fold change among significant neighborhoods > 0.5 together with > 50% significant neighborhoods; depletion flags required a median log_2_ fold change < −0.5 together with > 40% significant neighborhoods. No separate directional-coherence criterion was applied.

### PIPseq dataset

Eight conditions comprised day 4 and day 7 OPTi-OX and GERALT MYOD1 cells in hE6, and primary and directed-differentiation myoblasts before and after 4 d of fusion. Sample identities, experimental conditions, replicate structure and quality-control metrics are reported in Supplementary Table 9. Cells were enzymatically detached using TrypLE Express, collected by centrifugation at 300*g* for 5 min and washed in the cell-resuspension buffer supplied with the library-preparation kit. Cell concentration and viability were determined by trypan-blue exclusion. Viable suspensions were adjusted to 1,250 cells µL^−1^, and approximately 5,000 cells were loaded per reaction.

Single-cell 3^′^ RNA-sequencing libraries were generated using the Illumina Single Cell 3^′^ RNA Prep T2 workflow (Illumina, 20135689; document 200063178 v01). cDNA and final libraries were assessed using D5000 Screen-Tape, pooled and sequenced on a NextSeq 1000 using a P2 XLEAP-SBS flow cell (Illumina, 20100986), with read/index cycles 45/10/10/150.

BCL Convert 2.8.0 generated FASTQs using an experiment-specific dual-index sample sheet. DRAGEN Single Cell RNA 4.4.6000 and Reference Builder 4.4.6002 used GRCh38/Ensembl release 112 supplemented with the same custom features as the 10x Genomics reference. Libraries were processed separately, and automated cell calls were reviewed using library-specific quality-control profiles and knee plots before downstream analysis. Primary and processed data are available as described in the Data availability section.

Gene-by-cell count matrices were imported into R and analyzed using Seurat and Monocle 3. Protein-coding genes and the relevant custom transgene features were retained. For the PIPseq dataset, cells were retained if they contained > 100 and < 7,000 detected genes, > 100 RNA counts, and < 15% mitochondrial transcripts. Down-stream analysis followed the Seurat/Monocle 3 framework described for the 10x Genomics dataset, with dataset-specific quality-control thresholds. Clusters were annotated using canonical mesodermal, myogenic-progenitor, myoblast, fusion, myotube and myofiber markers together with the developmental and skeletal-muscle signatures in Supplementary Table 10. Cluster fractions were calculated using each sample’s retained cells as the denominator.

Cell-level differential expression was assessed using Seurat `FindMarkers` and `FindAllMarkers` with Wilcoxon rank-sum tests and Bonferroni-adjusted *P* values. Day-4 and day-7 GERALT-versus-OPTi-OX comparisons used the `riginalexp` assay’s data layer, with adjusted *p* < 0.01 and absolute `avg_log2FC` ≥ 1.

Gene-expression signatures were displayed as cluster-level dot plots using `monocle3::plot_genes_by_group`. Dot color represents mean log-normalized expression scaled to the range [−3, 3], whereas dot size indicates the fraction of cells with non-zero expression. Feature plots were generated with `plot_cells` using the default `scale_to_range = TRUE`, such that each gene was independently rescaled to its own expression range (`min_expr = 0.1`) and absolute intensities are not comparable across genes. No additional per-plot cell subsampling was applied.

Differences in cluster abundance were tested using pair-wise Fisher’s exact tests comparing OPTi-OX and GERALT MYOD1 at each time point and day 4 versus day 7 within each condition, with Benjamini–Hochberg correction within each comparison family (*P*_adj_ ≤ 0.01). Because only one scRNA-seq library was available per condition and individual cells were used as the statistical unit for cell-level differential-expression and cluster-abundance tests, these analyses quantify differences between the profiled libraries rather than reproducibility across biological replicates.

### Statistics and reproducibility

Unless indicated otherwise, data are presented as mean ± SEM. *N* denotes the number of independent biological experiments, cell lines, clones, differentiations or casting batches, as specified in the corresponding figure legend; *n* denotes observations nested within an independent experiment or batch. Technical replicates were averaged before statistical analysis.

Except for the transcriptomic and other bioinformatic analyses described in the preceding sections, statistical analyses were performed using GraphPad Prism version The statistical test, multiple-comparison procedure, sample size and, where applicable, matching or nesting structure used for each analysis are specified in the corresponding figure legend.

No statistical method was used to predetermine sample size; sample sizes were selected on the basis of previous experimental experience. Experiments were not randomized, and investigators were not blinded. No experimental data points were excluded beyond the sequencing quality-control procedures described above. Undetected RT-qPCR reactions were assigned a Ct value of 40, as specified above; no other missing values were imputed.

## Use of generative artificial intelligence

During preparation of this manuscript, the authors used ChatGPT (OpenAI) for language refinement and editorial assistance. After using this tool, the authors critically reviewed and edited the content as needed and take full responsibility for the final content of the manuscript.

## Data availability

Raw and processed sequencing data and associated sample metadata generated in this study have been deposited in BioStudies/ArrayExpress under accession numbers E-MTAB-17408 (bulk RNA-seq, myogenic circuit comparison), E-MTAB-17410 (bulk RNA-seq, TGF*β* modulation), E-MTAB-17411 (10x Genomics single-cell RNA-seq) and E-MTAB-17414 (PIPseq myogenic single-cell RNA-seq).

Custom genome/reference files and intermediate analysis objects required to reproduce the transcriptomic analyses are archived on Zenodo under doi:10.5281/zenodo.21471691. SNP-array GenomeStudio outputs and filtered CNV/LoH calls are also available in this archive.

## Code availability

Study-specific scripts and processing configurations used for bulk and single-cell RNA-seq analyses are available at https://github.com/alessandrobertero/Bottini-et-al-2026. Single-cell processing and analysis were performed within the Docker environment hedgelab/rstudio-hedgelab:iPS2seq_new_cellranger9.

The reusable bulk RNA-seq preprocessing and alignment pipeline is available at https://github.com/sara-bianchi/Bulk_RNA_seq_pipeline. CNV analysis and filtering scripts, including the genomic-instability hotspot definitions used in this study, are available at https://github.com/ebalmas/CNV-analysis-Bertero-lab/tree/main.

The EMCAP code used for engineered-muscle-tissue contractility analysis is archived on Zenodo under doi:10.5281/zenodo.18500644.

## Acknowledgments

We thank Matteo Sorge for assistance with Mantarray experiments, Francesca Anselmi for next-generation sequencing support through the NGS UniTo Platform, Sara Bianchi for assistance with RNA-seq processing and quality control, and Marta Gai and the Open Lab of Advanced Microscopy (OLMA@MBC) for imaging support. We are grateful to Samuele Marro for sharing lentiviral NGN2 reagents and protocols, Johannes Zuber for providing miR-F constructs and hairpin designs, and Emilia Turco for valuable scientific input. Selected schematics were created using BioRender.com.

This project has received funding from the European Union’s Horizon Europe Framework Programme for Research and Innovation through the European Innovation Council (EIC) Advanced Innovation Challenges pilot (SCALE-MYO, Grant Agreement No. 101327207, to A.B.). Views and opinions expressed are however those of the author(s) only and do not necessarily reflect those of the European Union or the European Innovation Council and SMEs Executive Agency (EISMEA). Neither the European Union nor the granting authority can be held responsible for them. Additional support was provided by the Italian Ministry of University and Research (MUR) under the National Recovery and Resilience Plan (NRRP), Mission 4, Component 2, Investment 1.1, funded by the European Union – NextGenerationEU (CUP D53D2302196000; Grant Assignment Decree No. 1377, to A.B.); the National Center for Gene Therapy and Drugs based on RNA Technology (PNRR Mission 4, Component 2, Investment 1.4, Spoke 2, project CN00000041, to P.D.); the PoC Instrument 2024 – PoC Launchpad programme of Fondazione Compagnia di San Paolo (to A.B.); and the Giovanni Armenise-Harvard Foundation (Career Development Award 2021, to A.B.). S.B. was supported by a PON R&I Azione IV.5 & IV.6 – FSE REACT-EU PhD fellowship. D.A. was supported through the University of Washington Global Innovation Fund.

## Author contributions

A.B. conceived the study and was responsible for project administration and funding acquisition. S.B. led the experimental investigation and development of the core experimental methodology, with additional investigation performed by F.A., D.A., M.D.G., M.A., and E.B.; D.A. additionally performed validation experiments. E.B. curated the data and, together with A.B., performed formal analyses and data visualization. P.B. developed software used in the study. F.B., D.L.M., P.E.P., P.D. and L.V. provided resources, including cell lines and experimental protocols. P.D., A.S.T.S., E.B. and A.B. provided supervision. A.B. wrote the original manuscript. S.B., F.A., E.B. and A.B. reviewed and edited the manuscript. All authors read and approved the final manuscript.

## References

1. Liu, W., Pang, P. D., Wu, C. A., Tagle, D., and Wu, J. C. (2, 2026). New approach methodologies for drug discovery. Cell 189, 1877–1903. 10.1016/j.cell.2026.02.012.

2. Lee, J., Herron, T., Studer, L., Mummery, C. L., and Wu, J. C. (27, 2026). Induced pluripotent stem cells from discovery to translation. Nat Med. 10.1038/s41591-026-04584-3.

3. Jara, T. C., Park, K., Vahmani, P., Van Eenennaam, A. L., Smith, L. R., and Denicol, A. C. (2023). Stem cell-based strategies and challenges for production of cultivated meat.eng. Nature Food 4, 841–853. 10.1038/s43016-023-00857-z.

4. Nelson, C. P., Brown, P., Fitzpatrick, S., Ford, K. A., Howard, P. C., MacGill, T., Margerrison, E. E. C., O’Shaughnessy, J., Patterson, T. A., Raghuwanshi, R., et al. (15, 2024). Advancing alternative methods to reduce animal testing. Science 386, 724–726. 10.1126/science.adg6228.

5. Edwards, M., Blanquie, O., Moriarty, O., Beken, S., and Ehmann, F. (2025). New approach methodologies: EU regulatory horizons. Nat Rev Drug Discov 24, 571–572. 10.1038/d41573-025-00053-7.

6. Ludwig, T. E., Andrews, P. W., Barbaric, I., Benvenisty, N., Bhattacharyya, A., Crook, J. M., Daheron, L. M., Draper, J. S., Healy, L. E., Huch, M., et al. (2023). ISSCR standards for the use of human stem cells in basic research. Stem Cell Reports 18, 1744–1752. 10.1016/j.stemcr.2023.08.003.

7. Selfa Aspiroz, L., Mennecozzi, M., Batlle, L., Corneo, B., Healy, L., Kotter, M., Kurtz, A., Ludwig, T. E., Mummery, C., Pera, M., et al. (8, 2025). Promoting the adoption of best practices and standards to enhance quality and reproducibility of stem cell research. Stem Cell Reports 20, 102531. 10.1016/j.stemcr.2025.102531.

8. Yamanaka, S. (2026). Two decades of induced pluripotent stem cell research: From discovery to diverse applications. Cell Stem Cell 33, 372–381. 10.1016/j.stem.2026.02.003.

9. Murry, C. E. and Keller, G. (2008). Differentiation of Embryonic Stem Cells to Clinically Relevant Populations: Lessons from Embryonic Development. Cell 132, 661–680. 10.1016/j.cell.2008.02.008.

10. Jenkins, M. J. and Farid, S. S. (2015). Human pluripotent stem cell-derived products: advances towards robust, scalable and cost-effective manufacturing strategies. Biotechnol J 10, 83–95. 10.1002/biot.201400348.

11. Francis, N., Aho, J., Ben-Nun, I. F., Bharti, K., Dianat, N., Makovoz, B., Nouri, P., Rothberg, J., Song, H., Zamilpa, R., et al. (2025). Scaling up pluripotent stem cell-based therapies-considerations, current challenges and emerging technologies: perspectives from the ISCT Emerging Regenerative Medicine Working Group. Cytotherapy 27, 1031–1042. 10.1016/j.jcyt.2025.04.058.

12. Sutcliffe, M. A., Wong, E., Wingett, S. W., Morris, C. A. J., Stachelscheid, H., Schoenfelder, S., and Lancaster, M. A. (7, 2026). Reversible epiblast regionalization determines differentiation potential of human pluripotent stem cells. Nat Biotechnol. 10.1038/s41587-026-03254-6.

13. Volpato, V., Smith, J., Sandor, C., Ried, J. S., Baud, A., Handel, A., Newey, S. E., Wessely, F., Attar, M., Whiteley, E., et al. (9, 2018). Reproducibility of Molecular Phenotypes after Long-Term Differentiation toăHuman iPSC-Derived Neurons: A Multi-Site Omics Study. Stem Cell Reports 11, 897–911. 10.1016/j.stemcr.2018.08.013.

14. Zhang, Y., Pak, C., Han, Y., Ahlenius, H., Zhang, Z., Chanda, S., Marro, S., Patzke, C., Acuna, C., Covy, J., et al. (5, 2013). Rapid single-step induction of functional neurons from human pluripotent stem cells. Neuron 78, 785–798. 10.1016/j.neuron.2013.05.029.

15. Ng, A. H. M., Khoshakhlagh, P., Rojo Arias, J. E., Pasquini, G., Wang, K., Swiersy, A., Shipman, S. L., Appleton, E., Kiaee, K., Kohman, R. E., et al. (2021). A comprehensive library of human transcription factors for cell fate engineering. en. Nature Biotechnology 39, 510–519. 10.1038/s41587-020-0742-6.

16. Pawlowski, M., Ortmann, D., Bertero, A., Tavares, J. M., Pedersen, R. A., Vallier, L., and Kotter, M. R. (2017). Inducible and Deterministic Forward Programming of Human Pluripotent Stem Cells into Neurons, Skeletal Myocytes, and Oligodendrocytes. Stem Cell Reports 8, 803–812. 10.1016/j.stemcr.2017.02.016.

17. Speicher, A. M., Korn, L., Csatári, J., Gonzalez-Cano, L., Heming, M., Thomas, C., Schroeter, C. B., Schafflick, D., Li, X., Gola, L., et al. (2022). Deterministic programming of human pluripotent stem cells into microglia facilitates studying their role in health and disease. eng. Proceedings of the National Academy of Sciences of the United States of America 119, e2123476119. 10.1073/pnas.2123476119.

18. Tomaz, R. A., Zacharis, E. D., Bachinger, F., Wurmser, A., Yamamoto, D., Petrus-Reurer, S., Morell, C. M., Dziedzicka, D., Wesley, B. T., Geti, I., et al. (2022). Generation of Functional Hepatocytes by Forward Programming with Nuclear Receptors. eLife 11, e71591. 10.7554/eLife.71591.

19. Vallier, L., Reynolds, D., and Pedersen, R. A. (2004). Nodal Inhibits Differentiation of Human Embryonic Stem Cells along the Neuroectodermal Default Pathway. Developmental Biology 275, 403–421. 10.1016/j.ydbio.2004.08.031.

20. James, D., Levine, A. J., Besser, D., and Hemmati-Brivanlou,A.(2005). TGFbeta/activin/nodal signaling is necessary for the maintenance of pluripotency in human embryonic stem cells. Development 132, 1273–82. 10.1242/dev.01706.

21. Xu, C., Rosler, E., Jiang, J., Lebkowski, J. S., Gold, J. D., O’Sullivan, C., Delavan-Boorsma, K., Mok, M., Bronstein, A., and Carpenter, M. K. (2005). Basic fibroblast growth factor supports undifferentiated human embryonic stem cell growth without conditioned medium. Stem cells 23, 315–23. 10.1634/stemcells.2004-0211.

22. Vallier, L., Alexander, M., and Pedersen, R. A. (2005). Activin/Nodal and FGF pathways cooperate to maintain pluripotency of human embryonic stem cells. Journal of cell science 118, 4495–509. 10.1242/jcs.02553.

23. Vallier, L., Mendjan, S., Brown, S., Chng, Z., Teo, A., Smithers, L. E., Trotter, M. W. B., Cho, C. H.-H., Martinez, A., Rugg-Gunn, P., et al. (2009). Activin/Nodal Signalling Maintains Pluripotency by Controlling Nanog Expression. Development 136, 1339–1349. 10.1242/dev.033951.

24. Bertero, A., Madrigal, P., Galli, A., Hubner, N. C., Moreno, I., Burks, D., Brown, S., Pedersen, R. A., Gaffney, D., Mendjan, S., et al. (2015). Activin/Nodal signaling and NANOG orchestrate human embryonic stem cell fate decisions by controlling the H3K4me3 chromatin mark. Genes & Development 29, 702–717. 10.1101/gad.255984.114.

25. Navarro, P., Festuccia, N., Colby, D., Gagliardi, A., Mullin, N. P., Zhang, W., Karwacki-Neisius, V., Osorno, R., Kelly, D., Robertson, M., et al. (12, 2012). OCT4/SOX2-independent Nanog autorepression modulates heterogeneous Nanog gene expression in mouse ES cells. EMBO J 31, 4547–4562. 10.1038/emboj.2012.321.

26. Maskalenka, K., Alagöz, G., Krueger, F., Wright, J., Rostovskaya, M., Nakhuda, A., Bendall, A., Krueger, C., Walker, S., Scally, A., et al. (18, 2023). NANOGP1, a tandem duplicate of NANOG, exhibits partial functional conservation in human naïve pluripotent stem cells. Development 150, dev201155. 10.1242/dev.201155.

27. Balmas, E., Ratto, M. L., Snijders, K. E., Becca, S., Liaci, C., Ricca, I., Merlo, G. R., Calogero, R. A., Alessandrí, L., Mendjan, S., et al. (2025). Single Cell Transcriptional Perturbome in Pluripotent Stem Cell Models. Molecular Systems Biology 22, 179–227. 10.1038/s44320-025-00172-8.

28. Dann, E., Henderson, N. C., Teichmann, S. A., Morgan,M. D., and Marioni, J. C. (2022). Differential abundance testing on single-cell data using k-nearest neighbor graphs.en. Nature Biotechnology 40, 245–253. 10.1038/s41587-021-01033-z.

29. Blahetek, G., Mayer, C., Zuber, J., Lenter, M., and Strobel, B. (10, 2024). Suppression of toxic transgene expression by optimized artificial miRNAs increases AAV vector yields in HEK-293 cells. Mol Ther Methods Clin Dev 32, 101280. 10.1016/j.omtm.2024.101280.

30. Truszkowski, L., Bottini, S., Bianchi, S., Scrivano, M. G., Ramondo, G. F., Belluci, L., Bell, H., Becca, S., Snijders, K. E., Savorè, G., et al. (2025). Refined and benchmarked homemade media for cost-effective, weekend-free human pluripotent stem cell culture. Open Research Europe 4, 192. 10.12688/openreseurope.18245.2.

31. Tan, J., Virtue, S., Norris, D. M., Conway, O. J., Yang, M., Bidault, G., Gribben, C., Lugtu, F., Kamzolas, I., Krycer, J. R., et al. (2024). Limited oxygen in standard cell culture alters metabolism and function of differentiated cells. EMBO J 43, 2127–2165. 10.1038/s44318-024-00084-7.

32. Smith, A. S., Luttrell, S. M., Dupont, J.-B., Gray, K., Lih, D., Fleming, J. W., Cunningham, N. J., Jepson, S., Hesson, J., Mathieu, J., et al. (2022). High-throughput, real-time monitoring of engineered skeletal muscle function using magnetic sensing.en. Journal of Tissue Engineering 13, 20417314221122127. 10.1177/20417314221122127.

33. Sozza, F., Romano, A., DElia, N., Terenzi, M., Ratto, M. L., Cliff, E. R., Nattenberg, G., Bianchi, S., Becca, S., Klug, H., et al. (2026). Combinatorial and Inducible CRISPRa/i Enables Canalized hiPSC Forward Programming and Iterative Refinement via Single-Cell Genomics.en. 10.64898/2026.05.31.729073.

34. Ihry, R. J., Worringer, K. A., Salick, M. R., Frias, E., Ho, D., Theriault, K., Kommineni, S., Chen, J., Sondey, M., Ye, C., et al. (2018). P53 Inhibits CRISPRCas9 Engineering in Human Pluripotent Stem Cells. Nature Medicine 24, 939–946. 10.1038/s41591-018-0050-6.

35. Assou, S., Girault, N., Plinet, M., Bouckenheimer, J., Sansac, C., Combe, M., Mianné, J., Bourguignon, C., Fieldes, M., Ahmed, E., et al. (14, 2020). Recurrent Genetic Abnormalities in Human Pluripotent Stem Cells: Definition and Routine Detection in Culture Supernatant by Targeted Droplet Digital PCR. Stem Cell Reports 14, 1–8. 10.1016/j.stemcr.2019.12.004.

36. Cyrys, K., Manstein, F., Triebert, W., Kriedemann, N., Hernandez-Bautista, C. A., and Zweigerdt, R. (22, 2026). Direct inoculation of bioreactor-controlled stirred suspension culture with cryopreserved human pluripotent stem cells. Nat Protoc. 10.1038/s41596-026-01398-y.

37. Nahon, D. M., Moerkens, R., Aydogmus, H., Lendemeijer, B., Martínez-Silgado, A., Stein, J. M., Dostani, M., Frimat, J.-P., Gontan, C., Graaf M. N. S. de, et al. (2024). Standardizing designed and emergent quantitative features in microphysiological systems. Nat. Biomed. Eng 8, 941–962. 10.1038/s41551-024-01236-0.

38. Barry, J., Abranches, E., Baptista, R. P., Barry, C., Brandsten, C., Bruce, K., Carpenter, M. K., Cavagnaro, J., Collins, L., Csaszar, E., et al. (10, 2026). Charting the translational pathway: ISSCR best practices for the development of PSC-derived therapies. Stem Cell Reports 21, 102784. 10.1016/j.stemcr.2025.102784.

39. Generali, M., Fujita, Y., Kehl, D., Hirosawa, M., Emmert, M. Y., Takahashi, J., Hoerstrup, S. P., and Saito, H. (2024). Purification technologies for induced pluripotent stem cell therapies. Nat Rev Bioeng 2, 930–943. 10.1038/s44222-024-00220-2.

40. Post, M. J., Levenberg, S., Kaplan, D. L., Genovese, N., Fu, J., Bryant, C. J., Negowetti, N., Verzijden, K., and Moutsatsou, P. (2020). Scientific, sustainability and regulatory challenges of cultured meat. Nature Food 1, 403–415. 10.1038/s43016-020-0112-z.

41. Genovese, N. J., Domeier, T. L., Telugu, B. P. V. L., and Roberts, R. M. (6, 2017). Enhanced Development of Skeletal Myotubes from Porcine Induced Pluripotent Stem Cells. Sci Rep 7, 41833. 10.1038/srep41833.

42. Kinoshita, M., Kobayashi, T., Planells, B., Klisch, D., Spindlow, D., Masaki, H., Bornelöv, S., Stirparo, G. G., Matsunari, H., Uchikura, A., et al. (2021). Pluripotent stem cells related to embryonic disc exhibit common self-renewal requirements in diverse livestock species. Development 148, dev199901. 10.1242/dev.199901.

43. Zhi, M., Gao, D., Yao, Y., Zhao, Z., Wang, Y., He, P., Feng, Z., Zhang, J., Huang, Z., Gu, W., et al. (8, 2024). Elucidation of the pluripotent potential of bovine embryonic lineages facilitates the establishment of formative stem cell lines. Cell Mol Life Sci 81, 427. 10.1007/s00018-024-05457-z.

44. Riquelme-Guzmán, C., Stout, A. J., Kaplan, D. L., and Flack, J. E. (2024). Unlocking the potential of cultivated meat through cell line engineering. iScience 27, 110877.10.1016/j.isci.2024.110877.

45. Johnson, H. and Monaco, A. (2025). Global developments in the regulation of cultivated meat: A comparative study of the EU, Singapore, US and Australia and New Zealand. Review of European, Comparative & International Environmental Law 34, 496–511. 10.1111/reel.70007.

46. Kreitzer, F. R., Salomonis, N., Sheehan, A., Huang, M., Park, J. S., Spindler, M. J., Lizarraga, P., Weiss, W. A., So, P.-L., and Conklin, B. R. (2013). A robust method to derive functional neural crest cells from human pluripotent stem cells. eng. American Journal of Stem Cells 2, 119–131.

47. Roberts, B., Hendershott, M. C., Arakaki, J., Gerbin, K. A., Malik, H., Nelson, A., Gehring, J., Hookway, C., Ludmann, S. A., Yang, R., et al. (2019). Fluorescent Gene Tagging of Transcriptionally Silent Genes in hiPSCs. Stem Cell Reports 12, 1145–1158. 10.1016/j.stemcr.2019.03.001.

48. Guichardaz, M., Bottini, S., Balmas, E., and Bertero, A. (2024). Overcoming the Silencing of Doxycycline-Inducible Promoters in hiPSC-derived Cardiomyocytes. Open Research Europe 4, 266. 10.12688/openreseurope.19024.1.

49. Fellmann, C., Hoffmann, T., Sridhar, V., Hopfgartner, B., Muhar, M., Roth, M., Lai, D. Y., Barbosa, I. A., Kwon, J. S., Guan, Y., et al. (2013). An optimized microRNA backbone for effective single-copy RNAi. Cell Reports 5, 1704–1713. 10.1016/j.celrep.2013.11.020.

50. Bertero, A., Pawlowski, M., Ortmann, D., Snijders, K., Yiangou, L., Cardoso de Brito, M., Brown, S., Bernard, W. G., Cooper, J. D., Giacomelli, E., et al. (2016). Optimized Inducible shRNA and CRISPR/Cas9 Platforms for in Vitro Studies of Human Development Using hPSCs. Development 143, 4405–18. 10.1242/dev.138081.

51. Chen, Y., Tristan, C. A., Chen, L., Jovanovic, V. M., Malley, C., Chu, P. H., Ryu, S., Deng, T., Ormanoglu, P., Tao, D., et al. (2021). A Versatile Polypharmacology Platform Promotes Cytoprotection and Viability of Human Pluripotent and Differentiated Cells. Nature Methods 18, 528–541. 10.1038/s41592-021-01126-2.

52. Yuan, J. S., Reed, A., Chen, F., and Stewart, C. N. (2006). Statistical analysis of real-time PCR data. BMC Bioinformatics 7, 85. 10.1186/1471-2105-7-85.

53. Schindelin, J., Arganda-Carreras, I., Frise, E., Kaynig, V., Longair, M., Pietzsch, T., Preibisch, S., Rueden, C., Saalfeld, S., Schmid, B., et al. (28, 2012). Fiji: an open-source platform for biological-image analysis. Nat Methods 9, 676–682. 10.1038/nmeth.2019.

54. Dobin, A., Davis, C. A., Schlesinger, F., Drenkow, J., Zaleski, C., Jha, S., Batut, P., Chaisson, M., and Gingeras, T. R. (2013). STAR: ultrafast universal RNA-seq aligner. Bioinformatics (Oxford, England) 29, 15–21. 10.1093/bioinformatics/bts635.

55. Love, M. I., Huber, W., and Anders, S. (2014). Moderated estimation of fold change and dispersion for RNA-seq data with DESeq2. Genome Biol 15, 550. 10.1186/s13059-014-0550-8.

56. Wu, T., Hu, E., Xu, S., Chen, M., Guo, P., Dai, Z., Feng, T., Zhou, L., Tang, W., Zhan, L., et al. (28, 2021). clusterProfiler 4.0: A universal enrichment tool for interpreting omics data. Innovation (Camb) 2, 100141. 10.1016/j.xinn.2021.100141.

57. Hao, Y., Stuart, T., Kowalski, M. H., Choudhary, S., Hoffman, P., Hartman, A., Srivastava, A., Molla, G., Madad, S., Fernandez-Granda, C., et al. (2024). Dictionary learning for integrative, multimodal and scalable single-cell analysis. en. Nature Biotechnology 42, 293–304. 10.1038/s41587-023-01767-y.

58. Cao, J., Spielmann, M., Qiu, X., Huang, X., Ibrahim,D. M., Hill, A. J., Zhang, F., Mundlos, S., Christiansen, L., Steemers, F. J., et al. (2019). The single-cell transcriptional landscape of mammalian organogenesis. Nature 566, 496–502. 10.1038/s41586-019-0969-x.

