## Extended Data Figures for "Genetic control couples TGF*β*-independent pluripotent stem cell renewal to deterministic forward programming across germ layers"

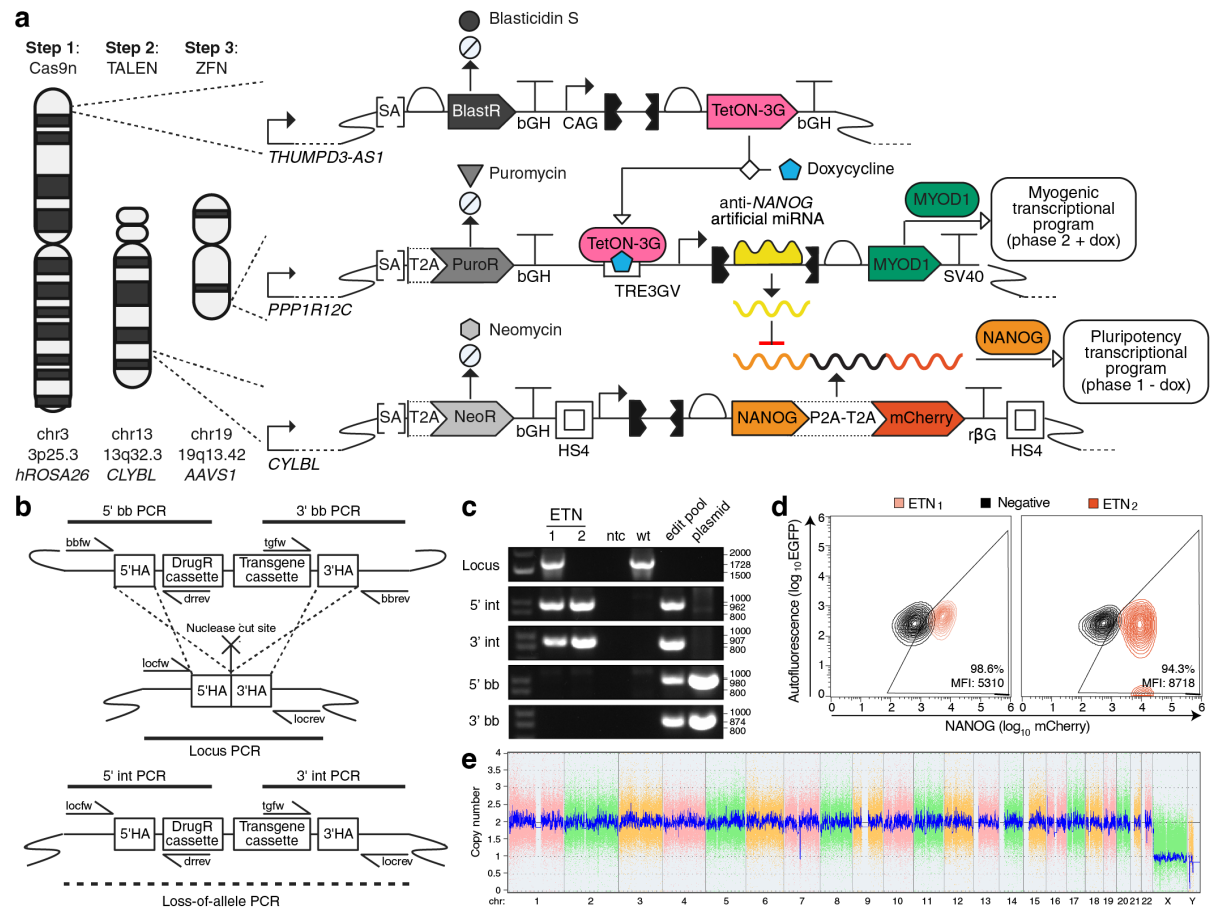

**Extended Data Fig. 1 | Genome engineering and validation of NANOG-overexpressing hiPSCs.**

**a**, Genome-engineering strategy represented using Synthetic Biology Open Language (SBOL) notation. WTC11 *TTN*-mEGFP hiPSCs were sequentially targeted with *S. pyogenes* D10A Cas9 nickases (Cas9n), transcription activator-like effector nucleases (TALENs), and zinc-finger nucleases (ZFNs) at the *hROSA26*, *CLYBL*, and *AAVS1* genomic safe-harbor loci, respectively. Tet-On-3G is constitutively expressed from *hROSA26*; *NANOG*<sub>exo</sub> and mCherry are constitutively co-expressed from *CLYBL*; and *AAVS1* carries a Tet-On-3G-responsive cassette encoding MYOD1 and an embedded anti-NANOG artificial microRNA. Chromosomal positions and genes encompassing the targeted loci are shown. SA, splice acceptor; BlastR, blasticidin resistance; PuroR, puromycin resistance; NeoR, neomycin resistance; bGH, bovine growth hormone polyadenylation signal; CAG, cytomegalovirus immediate-early enhancer/chicken beta-actin/rabbit beta-globin hybrid promoter; TRE3GV, third-generation tetracycline-responsive promoter; SV40, simian virus 40 polyadenylation signal; rβG, rabbit beta-globin polyadenylation signal; P2A and T2A, ribosome-skipping peptides; HS4, chicken beta-globin hypersensitive site 4 insulator. Constructs not shown to scale.

**b**, General PCR genotyping strategy for targeted integration. Site-specific integration was assessed by 5'INT and 3'INT junctional PCRs combining cassette primers with locus primers outside the corresponding homology arms. Retention of an unedited allele was assessed by locus or loss-of-allele PCR. For large, GC-rich insertions, including the CAG-containing *NANOG* cassette, these conditions amplify the shorter unedited allele but not the targeted allele; absence of the locus product together with positive 5'INT and 3'INT products is therefore consistent with biallelic integration. Donor-backbone integration was assessed by 5'BB and 3'BB PCRs. HA, homology arm; DrugR, drug-resistance cassette; INT, integration junction; BB, plasmid backbone; fw, forward primer; rev, reverse primer.

**c**, Representative PCR genotyping of the *NANOG-2A-mCherry* cassette at the *CLYBL* locus in ET hiPSCs, which carry *hROSA26*-targeted Tet-On-3G<sup>48</sup>. ETN<sub>1</sub> retained an unedited *CLYBL* allele and carried monoallelic integration, whereas ETN<sub>2</sub> lacked the locus product and carried biallelic integration. Both clones generated 5'INT and 3'INT products without detectable 5'BB or 3'BB products. ntc, no-template control; wt, parental WTC11 hiPSCs; edit pool, antibiotic-selected genome-edited pool used as a positive control for integration PCRs; plasmid, targeting plasmid used as a positive control for backbone PCRs.

**d**, Representative flow cytometry analysis of mCherry expression in ETN<sub>1</sub> and ETN<sub>2</sub> hiPSCs, overlaid with the parental ET negative control. mCherry is co-expressed with *NANOG*<sub>exo</sub> from the same polycistronic transcript. EGFP-channel fluorescence was used to visualize autofluorescence and facilitate discrimination of mCherry-positive cells from background. Percentages indicate mCherry<sup>+</sup> cells. MFI, median fluorescence intensity.

**e**, Genome-wide copy-number profile of ETN<sub>2</sub> hiPSCs by chromosomal microarray analysis (CMA). Colored probe signals represent individual chromosomes, and the blue trace shows the normalized smoothed copy-number estimate. The segmental chrY loss is a known alteration in the WTC11 donor background, whereas the small segmental loss on chromosome 7 was previously detected in the parental WTC11 *TTN*-mEGFP hiPSCs<sup>30</sup>. No additional newly acquired large-scale copy-number alterations were detected.

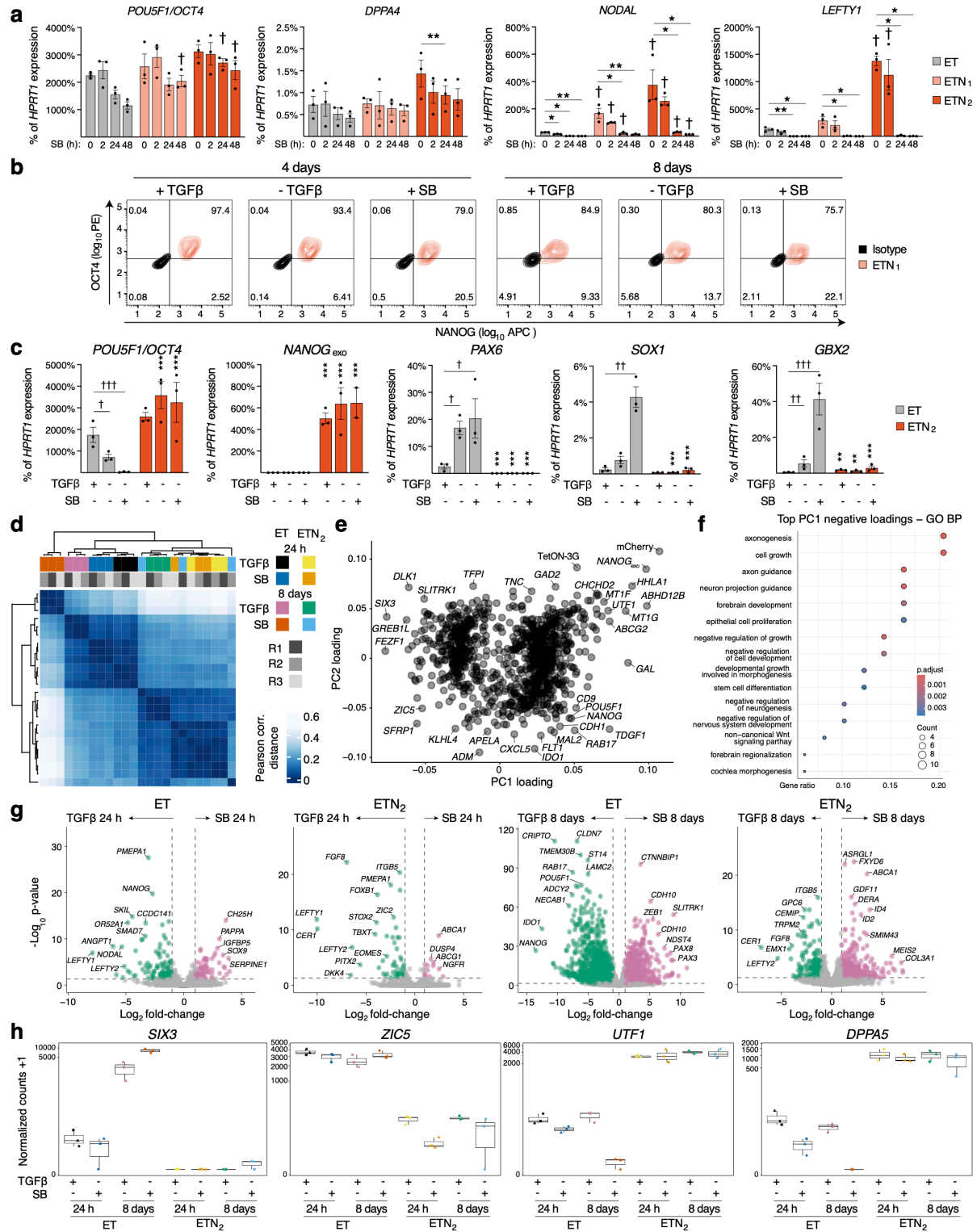

**Extended Data Fig. 2 | Extended characterization of TGFβ-independent pluripotency.**
Legend on the next page

**Extended Data Fig. 2 | Extended characterization of TGF $\beta$ -independent pluripotency (continued).**

- **a**, RT-qPCR analysis of additional pluripotency-associated (*POU5F1/OCT4* and *DPPA4*) and TGFβ-responsive (*NODAL* and *LEFTY1*) genes in the experiment described in Fig. 1c. N = 3 independent experiments; each point represents the mean of two parallel cultures. Statistical analysis and notation as in Fig. 1c.
- **b**, Representative flow cytometry analysis of NANOG and OCT4 expression in ETN<sub>1</sub> hiPSCs cultured for 4 or 8 days in E8, E7 or E7 supplemented with SB, as described for ET and ETN<sub>2</sub> hiPSCs in Fig. 1e. Isotype controls are shown in black.
- **c**, RT-qPCR analysis of pluripotency-associated genes and early neuroectodermal markers in ET and ETN<sub>2</sub> hiPSCs after 8 days under the conditions described in panel b. N = 3 independent experiments. Statistical analysis and notation as in Fig. 1e.
- **d**, Hierarchical clustering of bulk RNA-seq samples from Fig. 1f based on pairwise Pearson correlation distance. Annotation bars indicate genotype, culture duration, treatment condition and biological replicate.
- **e**, Gene loadings on PC1 and PC2 from the principal component analysis in Fig. 1f. Selected endogenous genes and transgenes are labeled.
- **f**, Gene Ontology Biological Process (GO BP) enrichment analysis of genes with the top 1% most negative PC1 loadings from panel e. Dot size indicates the number of genes associated with each term and color indicates the adjusted p value.
- **g**, Volcano plots of differential gene expression between SB-treated and time-matched TGFβ-containing baseline cultures in ET and ETN<sub>2</sub> hiPSCs after 24 h or 8 days. Genes enriched under baseline and SB-treated conditions are shown on the left and right, respectively, and colored green and pink. Differential-expression criteria as in Fig. 1g.
- **h**, Normalized RNA-seq expression of *SIX3*, *ZIC5*, *UTF1* and *DPPA5* from the experiment in Fig. 1f. Values are shown as normalized counts + 1; points represent N = 3 independent experiments.

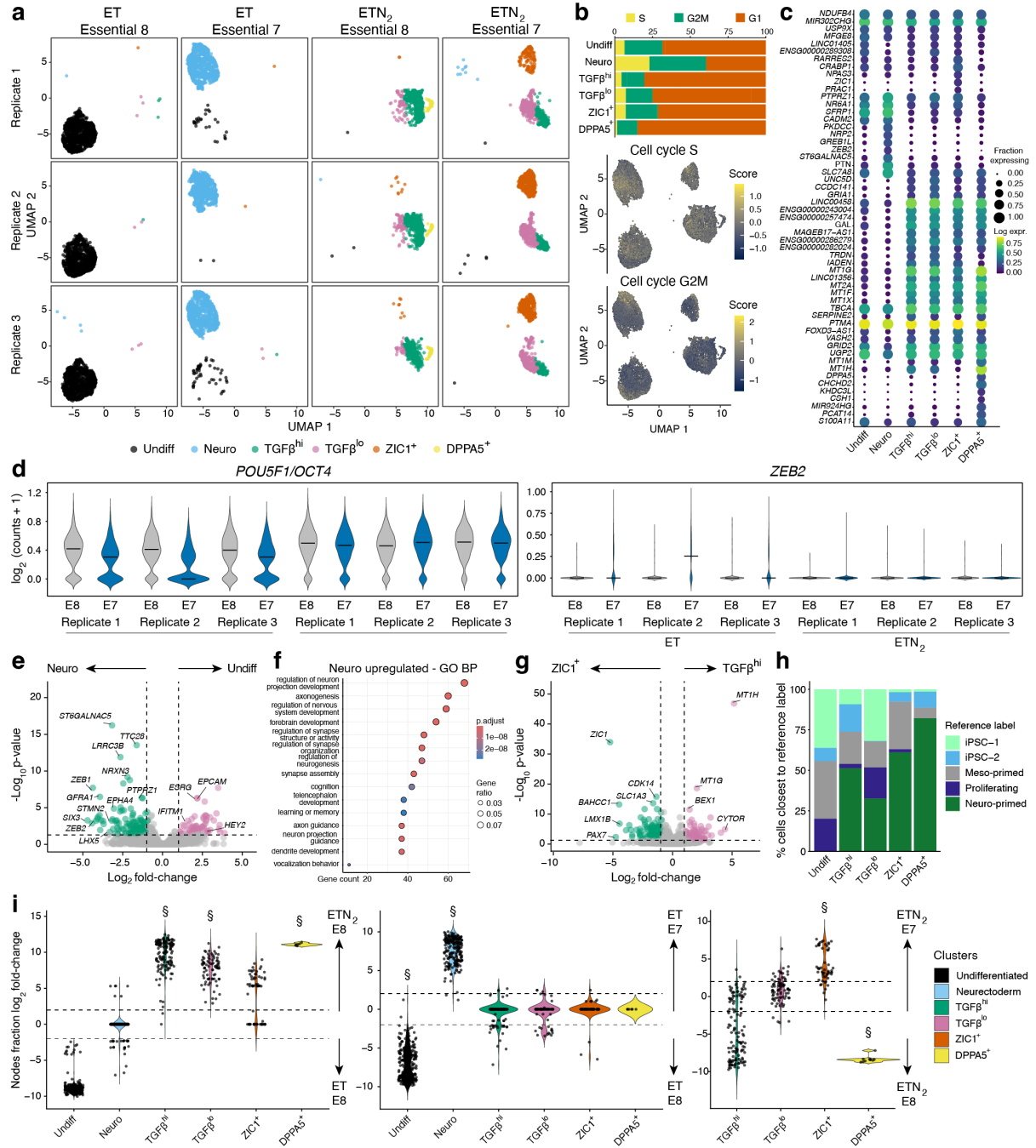

**Extended Data Fig. 3 | Single-cell characterization of transcriptional states during prolonged TGFβ-free culture.**
Legend on the next page

**Extended Data Fig. 3 | Single-cell characterization of transcriptional states during prolonged TGF $\beta$ -free culture (continued).**

- **a**, UMAP projections from Fig. 2e, split by biological replicate and colored by transcriptional cluster. Each row represents one of three independent long-term cultures.
- **b**, Cell-cycle analysis of the transcriptional clusters identified in Fig. 2e. The stacked bar chart shows the proportions of cells assigned to G1, S and G2/M phases within each cluster. UMAP feature plots show S-phase and G2/M scores.
- **c**, Top ten marker genes for each transcriptional cluster, identified using Monocle. Dot size indicates the fraction of cells expressing each gene and color indicates scaled log expression.
- **d**, Violin plots of *POU5F1/OCT4* and *ZEB2* expression in ET and ETN<sub>2</sub> hiPSCs cultured in E8 or E7, shown separately for each independent long-term culture. Values are shown as log<sub>2</sub>(counts + 1).
- **e**, Volcano plot of pseudobulk differential gene expression between the Neuro and Undiff clusters. Genes enriched in Neuro and Undiff cells are shown on the left and right, respectively, and colored green and pink. Differential-expression criteria as in Fig. 2d.
- **f**, Gene Ontology Biological Process (GO BP) enrichment analysis of genes upregulated in the Neuro cluster. The x axis indicates the number of genes associated with each term, dot size indicates the gene ratio, and color indicates the adjusted p value.
- **g**, Volcano plot of pseudobulk differential gene expression between the *ZIC1*<sup>+</sup> and TGFβ<sup>hi</sup> clusters. Genes enriched in the *ZIC1*<sup>+</sup> and TGFβ<sup>hi</sup> clusters are shown on the left and right, respectively, and colored green and pink. Differential-expression criteria as in Fig. 2d.
- **h**, Label-transfer analysis against a reference atlas of pluripotent states<sup>27</sup>. Bars show the percentage of cells in each transcriptional cluster assigned to the closest reference label. The Neuro cluster was excluded to restrict the analysis to pluripotent-appearing states.
- **i**, Milo differential-abundance analysis accounting for local transcriptional neighborhoods and biological replicate<sup>28</sup>. Distributions show neighborhood log<sub>2</sub> fold changes for ETN<sub>2</sub> E8 versus ET E8 (left), ET E7 versus ET E8 (middle), and ETN<sub>2</sub> E7 versus ETN<sub>2</sub> E8 (right), grouped by predominant transcriptional cluster. §, cluster descriptively classified as enriched if > 50% of its neighborhoods were significant (spatial FDR < 0.05) with a median log<sub>2</sub> fold change > 0.5, or as depleted if > 40% were significant with a median log<sub>2</sub> fold change < -0.5, calculated across significant neighborhoods only.

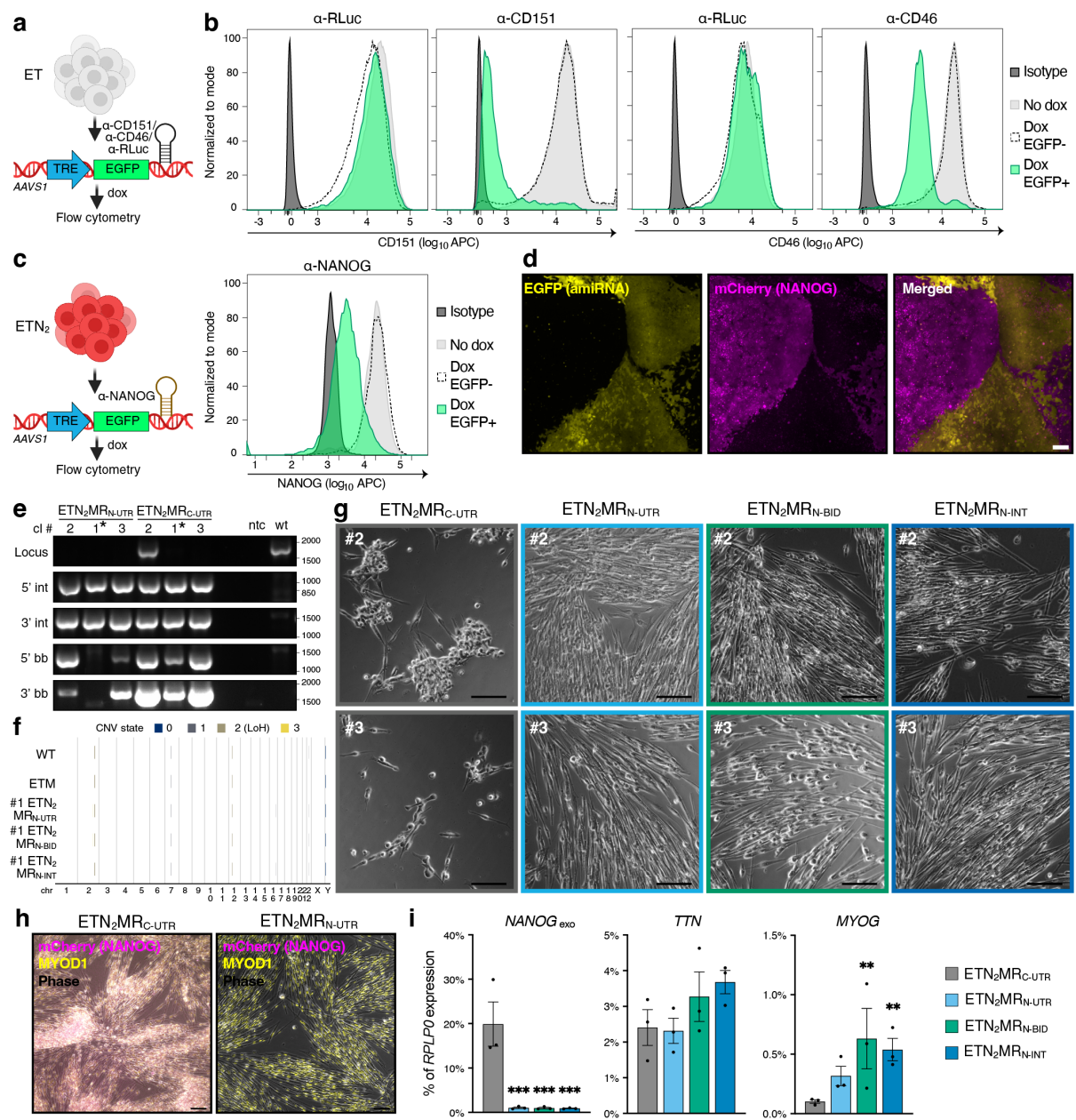

**Extended Data Fig. 4 | Establishment and validation of inducible miR-F silencing in hiPSCs.**
Legend on the next page

**Extended Data Fig. 4 | Establishment and validation of inducible miR-F silencing in hiPSCs (continued).**

- **a**, Schematic of inducible miR-F validation in ET hiPSCs. TRE-driven EGFP was coupled in its 3' UTR to amiRNAs targeting CD151, CD46, or Renilla luciferase (RLuc) as a control. The constructs were targeted to the *AAVS1* locus and analysed as antibiotic-selected pools without clonal isolation. Following 5 days of dox induction, EGFP-positive and negative populations were analysed by flow cytometry.
- **b**, Representative flow-cytometry histograms of CD151 and CD46 expression in the experiment described in panel a. Isotype controls, uninduced cells, and dox-treated EGFP-negative and EGFP-positive fractions are shown as indicated.
- **c**, Schematic and representative flow-cytometry analysis of inducible anti-NANOG miR-F activity in ETN<sub>2</sub> hiPSCs using the same experimental strategy as in panel a. NANOG FACS quantification presented as described in panel b.
- **d**, Representative fluorescence images from the experiment in panel c. EGFP reports induction of the amiRNA cassette and mCherry reports expression of *NANOG*<sub>exo</sub>. Scale bars, 100 μm.
- **e**, PCR genotyping of ETN<sub>2</sub> hiPSCs re-targeted at the *AAVS1* locus with dox-inducible *MYOD1* carrying either anti-NANOG or anti-CD151 control miR-F in its 3' UTR, corresponding to ETN<sub>2</sub>MR<sub>N-UTR</sub> and ETN<sub>2</sub>MR<sub>C-UTR</sub> in Fig. 3a. Three independently generated clones of each UTR design are shown. Genotyping logic and assay nomenclature are as in Extended Data Fig. 1b. Equivalent genotyping was performed for three ETN<sub>2</sub>MR<sub>N-BID</sub> clones, three ETN<sub>2</sub>MR<sub>N-INT</sub> clones, and one ETM clone. For the anti-NANOG architectures, only the biallelically targeted clone without detectable donor-backbone integration was taken forward for the experiments in Figs. 3a–3b.
- **f**, Genome-wide SNP-array copy-number variation (CNV) profiles of parental WT cells, ETM, and the selected ETN<sub>2</sub>MR<sub>N-UTR</sub>, ETN<sub>2</sub>MR<sub>N-BID</sub>, and ETN<sub>2</sub>MR<sub>N-INT</sub> clones. Colored segments indicate inferred copy-number states and copy-number-neutral loss of heterozygosity (LoH) as defined in the key. The larger chromosome 7 and Y alterations correspond to parental WTC11-background events also observed in Extended Data Fig. 1e. Copy-number-neutral LoH was additionally detected on chromosomes 2 and 12. Small, lower-confidence CNV calls on chromosomes 16 and 22 were detected in subsets of clones; the chromosome 22 event was also detected in WT cells, suggesting that it may be present in the parental background near the sensitivity limit of the assay.
- **g**, Representative phase-contrast images of two additional independently generated clones for each indicated amiRNA architecture after 7 days of myogenic forward programming under the conditions in Fig. 3a. Scale bars, 500 μm.
- **h**, Representative merged phase-contrast and fluorescence images of selected ETN<sub>2</sub>MR<sub>C-UTR</sub> and ETN<sub>2</sub>MR<sub>N-UTR</sub> clones after 7 days of forward programming. mCherry reports expression of *NANOG*<sub>exo</sub>. Scale bars, 50 μm.
- **i**, RT-qPCR analysis of transgenic *NANOG*, *TTN*, and *MYOG* after 7 days of forward programming in the indicated architectures. N = 3 clones per architecture. One-way ANOVA followed by Dunnett's multiple-comparison test against ETN<sub>2</sub>MR<sub>C-UTR</sub>; \*\* p < 0.01; \*\*\* p < 0.001; p > 0.05 not reported.

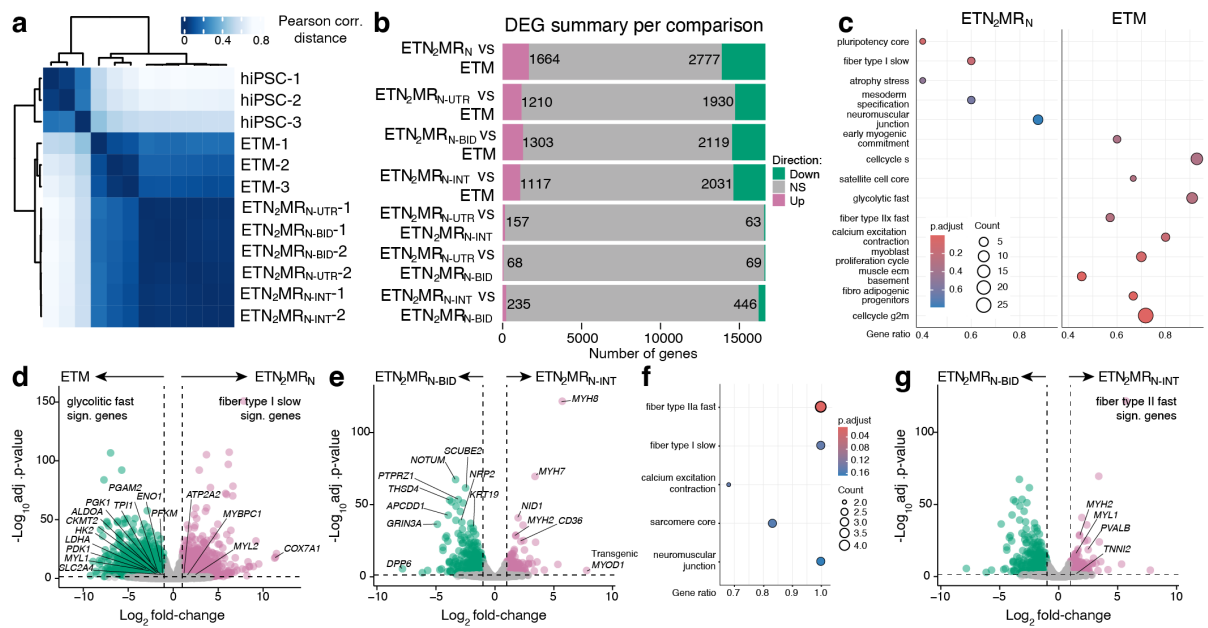

**Extended Data Fig. 5 | Transcriptomic benchmarking of ETN<sub>2</sub>MR<sub>N</sub> circuit architectures.**

- **a**, Hierarchical clustering of the bulk RNA-seq samples in Fig. 3d based on pairwise Pearson-correlation distance.
- **b**, Summary of differentially expressed genes for the indicated pooled ETN<sub>2</sub>MR<sub>N</sub> versus ETM and pairwise comparisons. Bars show genes up- or down-regulated in the first versus second condition. NS, not significant. Differential-expression criteria as in Fig. 3e.
- **c**, Gene-set enrichment analysis of pooled ETN<sub>2</sub>MR<sub>N</sub> versus ETM using manually curated pluripotency, myogenic, fiber-type, cell-cycle and lineage-associated signatures. Dot size indicates gene count, the x axis indicates the adjusted p value, and color indicates the adjusted p value.
- **d**, Volcano plot of differential expression between ETM and pooled ETN<sub>2</sub>MR<sub>N</sub> samples. Genes enriched in ETM and pooled ETN<sub>2</sub>MR<sub>N</sub> are shown on the left and right, respectively, and colored green and pink. Selected genes from the glycolytic-fast and fiber-type-I-slow signatures are labelled. Differential-expression criteria as in Fig. 3e.
- **e**, Volcano plot of differential expression between ETN<sub>2</sub>MR<sub>N</sub>-BID and ETN<sub>2</sub>MR<sub>N</sub>-INT. Genes enriched in BID and INT are shown on the left and right, respectively, and colored green and pink. Selected lineage- and muscle-associated genes are labelled, including transgenic MYOD1. Differential-expression criteria as in Fig. 3e.
- **f**, Gene-set enrichment analysis of ETN<sub>2</sub>MR<sub>N</sub>-BID versus ETN<sub>2</sub>MR<sub>N</sub>-INT using the indicated manually curated muscle-related signatures. Plotting as in panel c.
- **g**, As in panel e, with selected genes contributing to the fiber-type-II-fast signature highlighted.

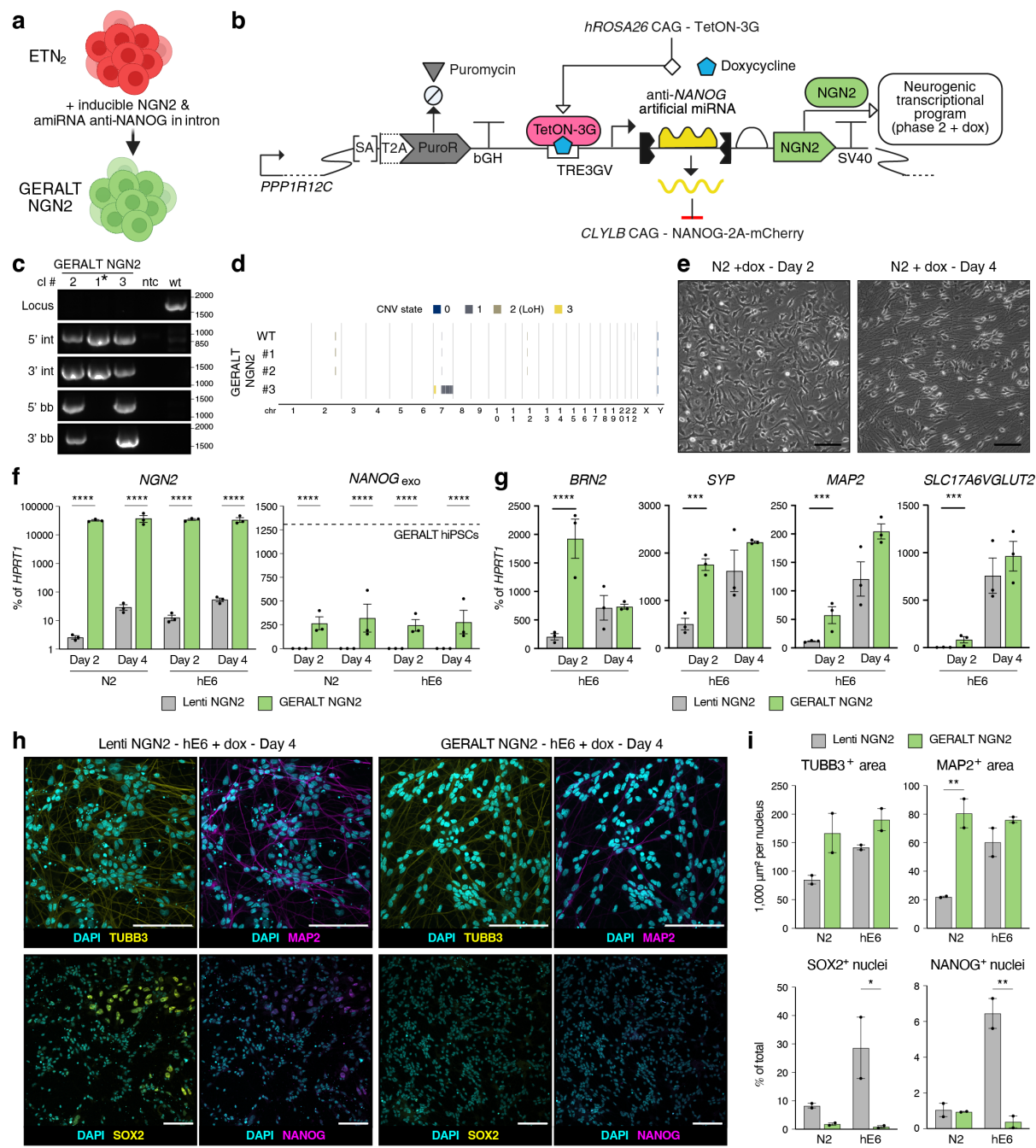

**Extended Data Fig. 6 | Generation and extended characterization of GERALT NGN2 hiPSCs.**
Legend on the next page

**Extended Data Fig. 6 | Generation and extended characterization of GERALT NGN2 hiPSCs (continued).**

- **a**, Schematic of the generation of GERALT NGN2 cells from ETN<sub>2</sub> hiPSCs by addition of a dox-inducible *NEUROG2/NGN2* cassette coupled to an intronic anti-NANOG amiRNA.
- **b**, Architecture of the GERALT NGN2 AAVS1-targeting cassette in SBOL notation. Constitutive Tet-On-3G expression from *hROSA26* and *NANOG-2A-mCherry* expression from *CLYBL* are indicated schematically; symbols and abbreviations are as in Extended Data Fig. 1a. Construct not shown to scale.
- **c**, PCR genotyping at the AAVS1 locus of three GERALT NGN2 clones. Genotyping logic and assay nomenclature are as in Extended Data Fig. 1b. A biallelically targeted clone without detectable donor-backbone integration, marked by an asterisk, was selected for the analyses in panels e, h and in Figs. 4c–4d.
- **d**, Genome-wide SNP-array CNV profiles of parental WT cells and three GERALT NGN2 clones. The parental WT profile is the same dataset shown in Extended Data Fig. 4f; plotting and CNV/LoH notation are as in that panel. Two GERALT NGN2 clones showed only alterations also present in the parental WT profile, whereas the third carried additional large-scale copy-number abnormalities on chromosome 7.
- **e**, Representative phase-contrast images of the selected GERALT NGN2 clone after 2 and 4 days of forward programming in N2 + dox. Scale bars, 100 μm.
- **f**, RT-qPCR analysis of inducible *NEUROG2/NGN2* and transgenic *NANOG* during the benchmarking experiment described in Fig. 4a. The dotted line in the *NANOG*<sub>exo</sub> plot indicates the level in undifferentiated GERALT NGN2 hiPSCs. N = 3 lentiviral-transduced lines and N = 3 GERALT NGN2 clones; each point represents the mean of two independent differentiations. Statistical analysis and notation as in Fig. 4b.
- **g**, RT-qPCR analysis of the neuronal markers in Fig. 4b during forward programming in hE6 + dox. N, replicate structure, statistical analysis and notation as in Fig. 4b.
- **h**, Representative immunofluorescence of lentiviral *NEUROG2/NGN2*-transduced ET hiPSCs and the selected GERALT NGN2 clone after 4 days of forward programming in hE6 + dox. Scale bars, 100 μm.
- **i**, Quantification of the neuronal and residual pluripotency markers in panel h and the corresponding N2 experiment in Fig. 4c. TUBB3- and MAP2-positive areas and the fractions of SOX2- and NANOG-positive nuclei are shown. N = 2 Lenti NGN2 lines and N = 2 GERALT NGN2 clones; each point represents the mean of two technical replicates. Two-way ANOVA followed by Šidák's multiple-comparison test between platforms within each medium; statistical notation as in Fig. 4b.

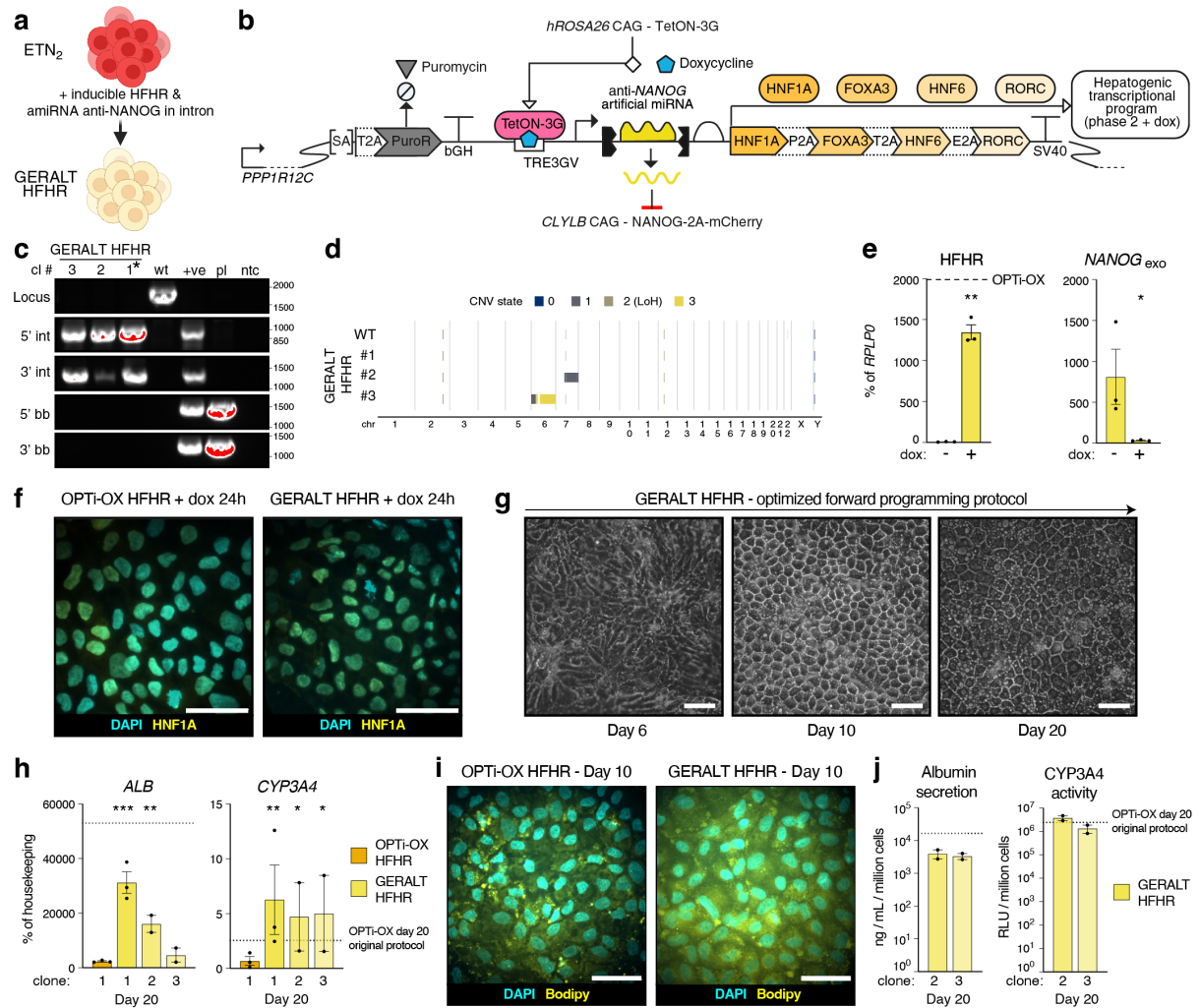

**Extended Data Fig. 7 | Generation and extended characterization of GERALT HFHR hiPSCs.**

**a**, Schematic of the generation of GERALT HFHR cells from ETN<sub>2</sub> hiPSCs by addition of a dox-inducible HFHR hepatic-programming cassette coupled to an intronic anti-NANOG amiRNA.

**b**, Architecture of the GERALT HFHR AAVS1-targeting cassette in SBOL notation. The TRE3GV-driven transcription unit contains an intronic anti-NANOG amiRNA followed by *HNF1A*, *FOXA3*, *ONECUT1/HNF6*, and *RORC* separated by 2A peptides. Constitutive Tet-On-3G expression from *hROSA26* and *NANOG-2A-mCherry* expression from *CLYBL* are indicated schematically; symbols and abbreviations are as in Extended Data Fig. 1a. Construct not shown to scale.

**c**, PCR genotyping at the AAVS1 locus of three GERALT HFHR clones. Genotyping logic and assay nomenclature are as in Extended Data Fig. 1b. A biallelically targeted clone without detectable donor-backbone integration, marked by an asterisk, was selected as the lead clone for the main phenotypic analyses in Figs. 4f–4h and panels f, g and i.

**d**, Genome-wide SNP-array CNV profiles of parental WT cells and three GERALT HFHR clones. The parental WT profile and CNV/LoH notation are as in Extended Data Fig. 4f. The selected clone 1 contained only alterations also detected in parental WT cells, whereas clones 2 and 3 carried additional large copy-number alterations on chromosomes 7 and 6, respectively.

**e**, RT-qPCR analysis of the inducible HFHR transgene cassette and transgenic *NANOG* in three independently generated GERALT HFHR clones without or with dox after 24 h of induction. The dotted line in the HFHR plot indicates the corresponding OPTi-ox HFHR reference value. N = 3 clones. Two-tailed paired t-tests. Here and in panel h, \* p < 0.05; \*\* p < 0.01; \*\*\* p < 0.001; p > 0.05 not reported.

**f**, Representative immunofluorescence of OPTi-ox HFHR and the selected GERALT HFHR clone after 24 h of dox induction. Scale bars, 50 μm.

**g**, Representative phase-contrast images of the selected GERALT HFHR clone at days 6, 10 and 20 of the GERALT-optimized protocol described in Fig. 4e. Scale bars, 50 μm.

**h**, RT-qPCR analysis of *ALB* and *CYP3A4* at day 20 in cultures of OPTi-ox HFHR and GERALT HFHR clones 1–3 under the GERALT-optimized conditions in Fig. 4e. Expression is normalized to the geometric mean of *PBGD* and *RPLP0*. Dotted lines indicate OPTi-ox HFHR reference values obtained using the reference protocol. N = 2–3 independent differentiations. Repeated-measures mixed-effects model followed by Dunnett's multiple-comparison test against OPTi-ox HFHR; statistical notation as in panel e.

**i**, Representative BODIPY 493/503 staining of lipid droplets in OPTi-ox HFHR and GERALT HFHR cells after 10 days of forward programming, with DAPI nuclear counterstaining. Scale bars, 50 μm.

**j**, Albumin secretion and CYP3A4 activity at day 20 in the two additional GERALT HFHR clones (clones 2 and 3), extending the analysis in Fig. 4h. Dotted lines indicate OPTi-ox HFHR reference values obtained using the reference protocol. N = 2 differentiations.

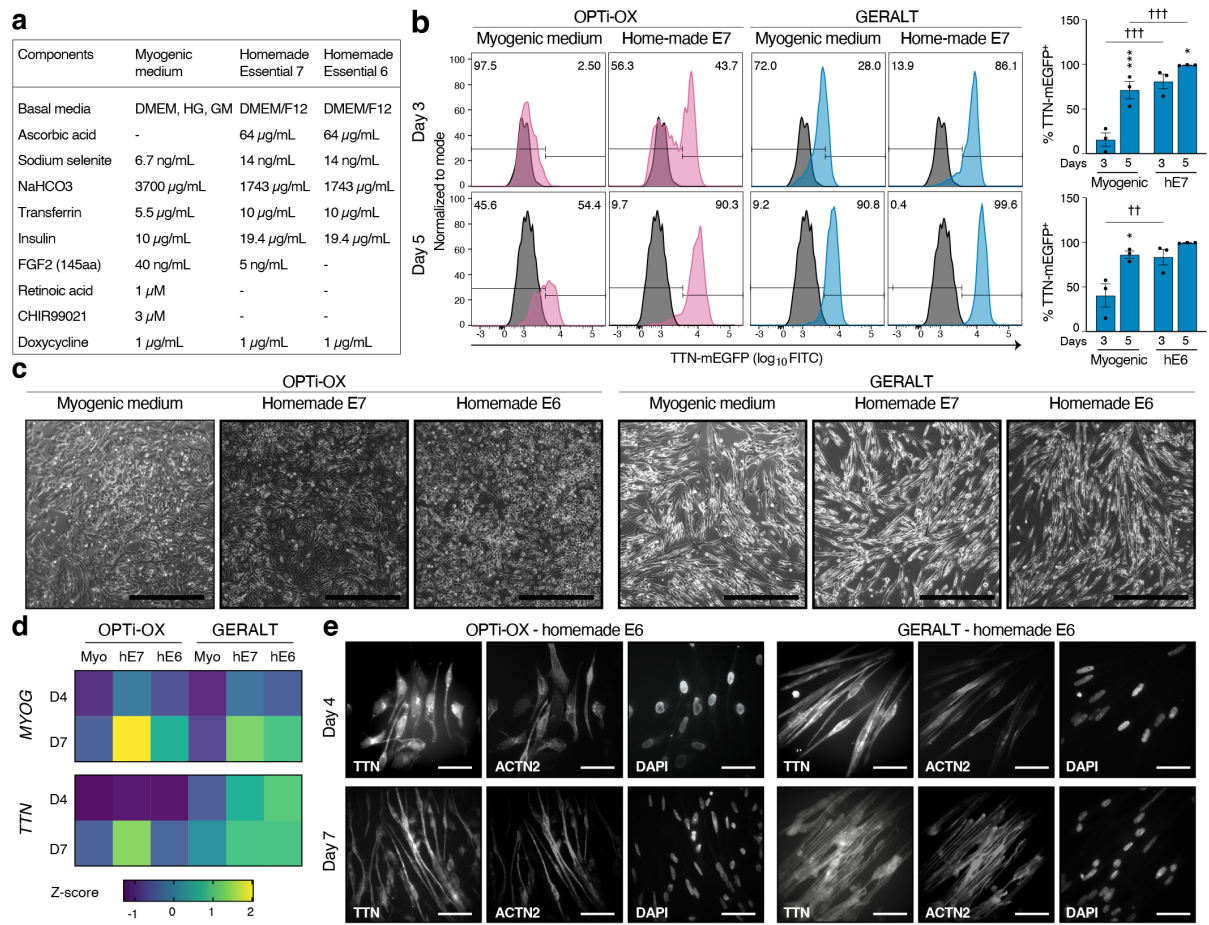

**Extended Data Fig. 8 | Optimization of GERALT myogenic differentiation in minimal media.**

**a**, Composition of the myogenic medium, hE7 and hE6 formulations used for the experiments in Fig. 5.

**b**, Additional TTN-mEGFP analysis complementing Fig. 5b. Representative flow-cytometry histograms show OPTi-OX and GERALT cells at days 3 and 5 in myogenic medium and hE7. Right, quantification of TTN-mEGFP<sup>+</sup> GERALT cells across three independently generated clones, comparing myogenic medium with hE7 (top) or hE6 (bottom). Repeated-measures two-way ANOVA followed by Sidák's multiple-comparison test. Asterisks denote comparisons between time points within each media and daggers comparisons between media at matched time points; \* $p < 0.05$; \*\*\* $p < 0.001$; † $p < 0.01$; †† $p < 0.001$.

**c**, Representative phase-contrast images of OPTi-OX and GERALT cells after 4 days of forward programming in myogenic medium, hE7 or hE6. Scale bars, 500 $\mu\text{m}$.

**d**, RT-qPCR analysis of *MYOG* and *TTN* at days 4 and 7 under the conditions in Fig. 5a. Heatmaps show mean gene-wise z-scores calculated from *HPRT1*-normalized expression across $N = 2$ cultures.

**e**, Individual grayscale channels corresponding to the immunofluorescence images in Fig. 5c. Scale bars, 50 $\mu\text{m}$.

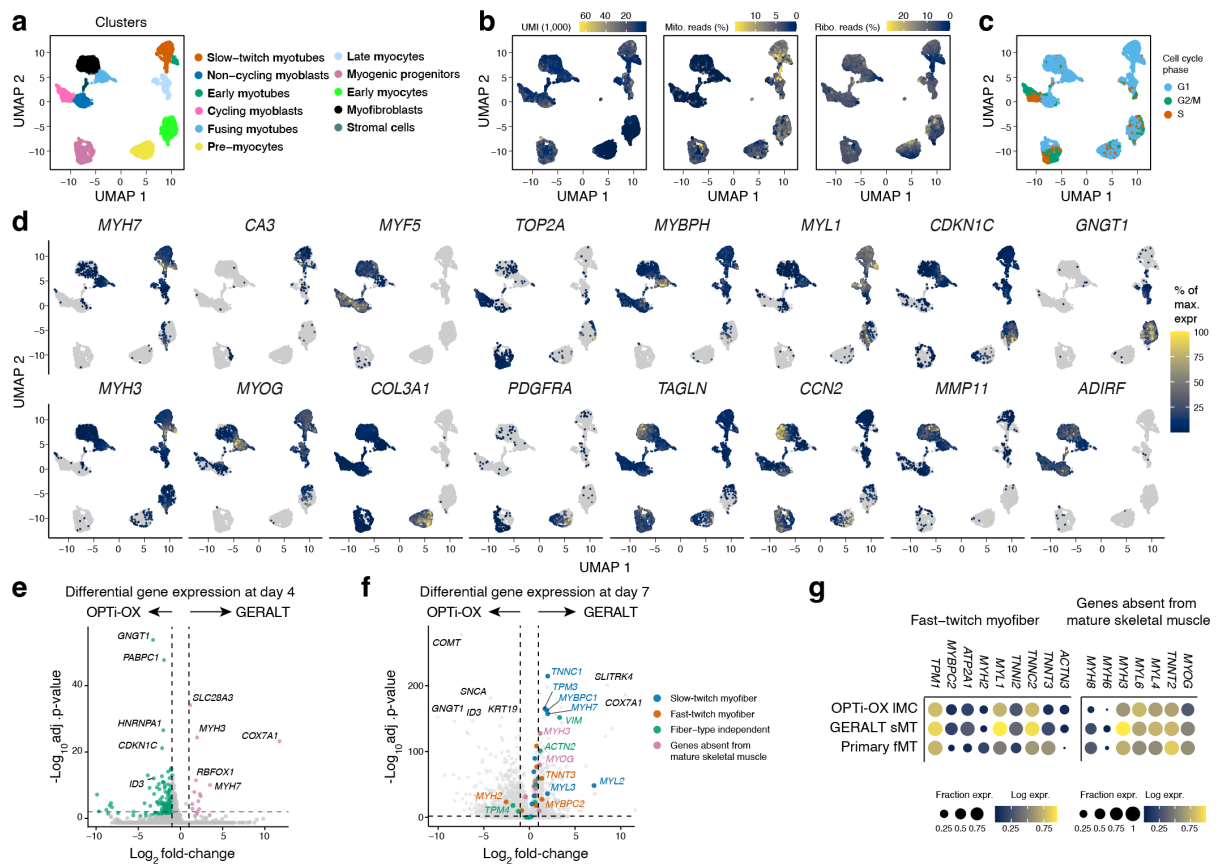

**Extended Data Fig. 9 | Single-cell RNA-seq benchmarking of GERALT myogenic differentiation.**

**a**, UMAP of the complete single-cell RNA-seq dataset in Fig. 5d, colored by transcriptional cluster.

**b**, Quality-control features projected onto the integrated UMAP: UMI counts (thousands), mitochondrial transcript fraction, and ribosomal transcript fraction.

**c**, Cell-cycle phase projected onto the integrated UMAP.

**d**, Feature plots of representative genes supporting the transcriptional-cluster annotations in Fig. 5d. Expression is shown as a percentage of maximum expression for each gene.

**e**, Volcano plot of differential gene expression between OPTi-OX and GERALT cells at day 4, complementing Fig. 5g. Genes enriched in OPTi-OX and GERALT are shown on the left and right, respectively, and colored green and pink. Selected genes are labelled. Differential-expression criteria as in Fig. 5g.

**f**, Volcano plot of differential gene expression between OPTi-OX and GERALT cells at day 7. Genes enriched in OPTi-OX and GERALT are shown on the left and right, respectively. Colors identify the indicated curated muscle signatures; selected genes are labelled. Differential-expression criteria as in Fig. 5g.

**g**, Complementary analysis to Fig. 5h, showing fast-twitch-myofiber and mature-muscle-absent gene signatures in the indicated OPTi-OX late-myocyte (IMC), GERALT slow-twitch-myotube (sMT), and primary fusing-myotube (fMT) populations. Dot size indicates the fraction of cells expressing each gene and color indicates log expression.

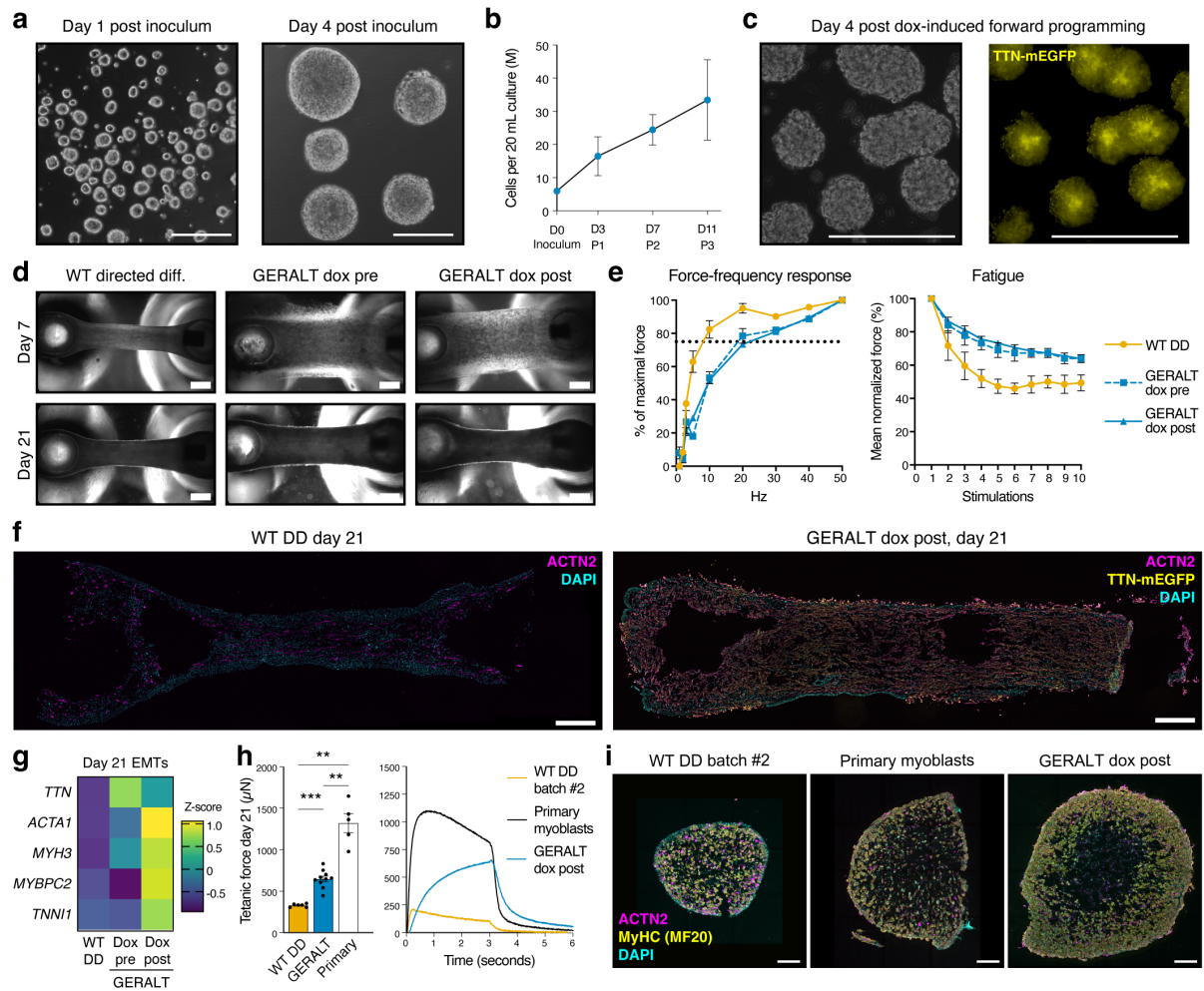

**Extended Data Fig. 10 | Extended characterization and benchmarking of GERALT engineered muscle tissues.**

**a**, Low-magnification phase-contrast images of GERALT suspension cultures at days 1 and 4 after inoculation. Scale bars, 200 μm.

**b**, GERALT suspension-expansion kinetics, showing cell yield at inoculation and after three successive passages. $N = 3$ clones.

**c**, Low-magnification phase-contrast and TTN-mEGFP fluorescence images of GERALT aggregates after 4 days of dox-induced myogenic forward programming in suspension, complementing Fig. 6d. Scale bars, 200 μm.

**d**, Representative phase-contrast images of WT directed-differentiated (WT DD), GERALT dox-pre and GERALT dox-post EMTs at days 7 and 21 after casting under the conditions in Fig. 6f. Scale bars, 200 μm.

**e**, Functional characterization of a subset of the EMTs in Fig. 6g. Left, force-frequency response expressed as a percentage of maximal force across increasing electrical-stimulation frequencies. Right, fatigue response over 10 repeated stimulations, expressed as mean normalized force. $N = 2$ independent casting batches; $n = 4, 14$ and $9$ EMTs for WT DD, GERALT dox-pre and GERALT dox-post, respectively.

**f**, Representative longitudinal sections of day 21 WT DD and GERALT dox-post EMTs. Scale bars, 750 μm.

**g**, RT-qPCR heatmap of the indicated myogenic genes in day 21 WT DD, GERALT dox-pre and GERALT dox-post EMTs. Heatmaps show gene-wise z-scores calculated from *HPRT1*-normalized expression. $n = 2$ EMTs per condition.

**h**, Day 21 tetanic-force benchmark of GERALT dox-post EMTs and EMTs generated from a second WT DD differentiation batch and primary human myoblasts, with representative traces shown at right. $n = 6, 10$ and $5$ EMTs for WT DD, GERALT dox-post and primary-myoblast EMTs, respectively, from one casting batch. One-way ANOVA followed by Tukey's multiple-comparison test. \*\* $p < 0.01$; \*\*\* $p < 0.001$.

**i**, Representative cross-sections of EMTs from panel h. Scale bars, 250 μm.
