## Supplemental Information for "Genetic control couples TGF*β*-independent pluripotent stem cell renewal to deterministic forward programming across germ layers"

### Supplementary information

#### Supplementary Note 1 - Hepatic protocol optimization

##### Optimization of GERALT-HFHR hepatic forward programming

To establish whether the GERALT architecture could support hepatic forward programming, we initially transferred the published OPTi-OX HFHR differentiation protocol<sup>31</sup> directly to GERALT HFHR cells. In this protocol, HFHR expression was induced continuously for 10 days while cells were maintained under hypoxia at 37°C in Hepatozyme low-insulin (HLI) medium containing 2% lipids, followed by maturation with transfer to normoxia at 39°C from day 15 (Supplementary Fig. 1a). Under these conditions, GERALT HFHR cells showed a hepatic-marker expression profile distinct from the OPTi-OX comparator (Supplementary Fig. 1b) and a markedly different day 10 morphology, lacking the more homogeneous polygonal appearance of OPTi-OX HFHR cells and displaying prominent intracellular vacuole- or lipid droplet-like structures (Supplementary Fig. 1c). The condition was also poorly tolerated by GERALT HFHR cultures, with extensive cell death such that two of three differentiations did not reach day 20. Together, these observations indicated that direct transfer of the OPTi-OX protocol was suboptimal for the GERALT HFHR architecture.

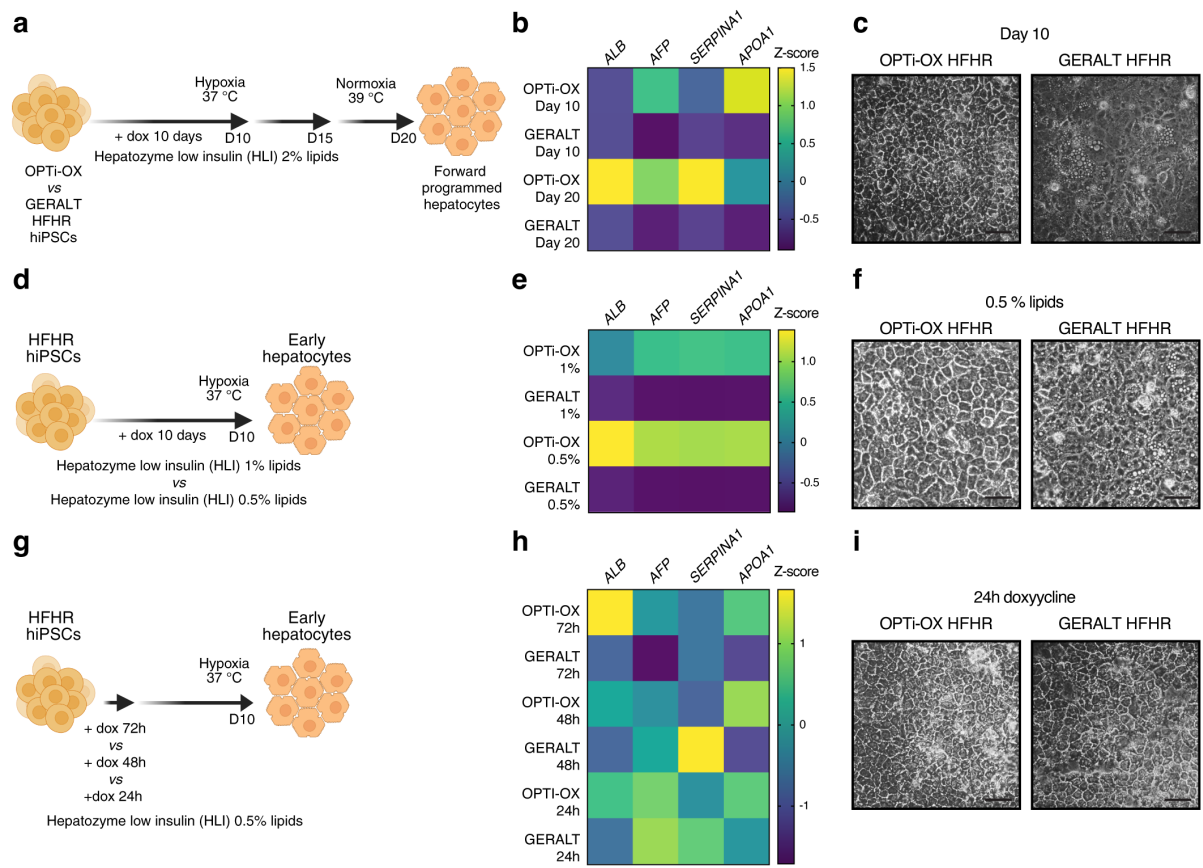

**Supplementary Fig. 1 | Stepwise optimization of GERALT HFHR hepatic forward programming.**

- Initial hepatic forward-programming protocol applied to OPTi-OX HFHR and GERALT HFHR (clone 1; Extended Data Fig. 7d).
- RT-qPCR analysis of hepatic markers under the conditions in panel a. Here and in panels e and h, expression was normalized to the geometric mean of *RPLP0* and *PBGD* before gene-wise z-scoring. At day 10, N = 3 independent differentiations per platform; at day 20, N = 3 OPTi-OX HFHR and N = 1 GERALT HFHR differentiation, as two GERALT cultures were lost to extensive cell death.
- Representative phase-contrast images at day 10 under the conditions in panel a. Here and in panels f and i, scale bars, 50 µm.
- Lipid-supplementation optimization during hepatic forward programming, comparing HLI containing 1% or 0.5% lipids.
- RT-qPCR analysis at day 10 under the conditions in panel d. N = 3 GERALT HFHR and N = 1 OPTi-OX HFHR independent differentiation experiments, each evaluated across both lipid conditions. Plotting and normalization as in panel b.
- Representative phase-contrast images under the 0.5% lipid condition in panel d.
- HFHR induction-time optimization, comparing 72, 48 and 24 h of dox exposure.
- RT-qPCR analysis at day 10 under the conditions in panel g. One independent differentiation experiment was analysed for each platform across induction-duration conditions. Plotting and normalization as in panel b.
- Representative phase-contrast images following 24 h of dox induction in panel g.

The conspicuous lipid-associated morphology and poor culture tolerance prompted us to first reduce lipid supplementation while leaving the duration of HFHR induction unchanged. We therefore compared HLI containing 1% or 0.5% lipids during the first 10 days of differentiation (Supplementary Fig. 1d). Reducing lipid supplementation did not by itself restore an OPTi-OX-like hepatic-marker expression profile (Supplementary Fig. 1e), but 0.5% lipids yielded a more homogeneous epithelial monolayer with reduced accumulation of the prominent intracellular structures observed under the original conditions (Supplementary Fig. 1f). We therefore retained 0.5% lipid supplementation for subsequent optimization.

We next considered whether prolonged lineage-driver expression was necessary in this configuration. Because the GERALT HFHR cassette is stably integrated at the *AAVS1* locus and showed robust doxycycline-inducible expression (Extended Data Figs. 7e–7f), we compared 72, 48, and 24 h periods of dox exposure while maintaining HLI containing 0.5% lipids (Supplementary Fig. 1g). Marker responses varied across induction durations rather than changing uniformly, with no condition maximizing all transcripts examined; however, 24 h of dox exposure provided the most favourable overall combination of hepatic-marker expression and culture morphology (Supplementary Fig. 1h; Supplementary Fig. 1i). We therefore selected 24 h of HFHR induction for subsequent experiments.

Overall, these experiments showed that the published OPTi-OX HFHR protocol could not be transferred unchanged to the GERALT background. Reduced lipid supplementation and shorter HFHR induction were selected empirically, while the earlier morphological progression observed in GERALT HFHR cultures by day 6 (Extended Data Fig. 7g) prompted an earlier transition to the 39°C normoxic maturation phase. The resulting protocol, incorporating 0.5% lipid supplementation, 24 h of dox exposure, and the earlier maturation transition, was subsequently used for the hepatic characterization reported in Figs. 4e–4h and Extended Data Figs. 7g–7j.

#### Supplementary Note 2 - Plasmid construction

##### General procedures

Restriction/ligation cloning used enzymes from New England Biolabs or Thermo Fisher Scientific and T4 DNA ligase (Thermo Fisher Scientific, EL0011). HiFi assembly used NEBuilder HiFi DNA Assembly Master Mix (New England Biolabs, E2621). PCR used Q5 Hot Start High-Fidelity DNA Polymerase (New England Biolabs, M0493). DNA ends were repaired using the Fast DNA End Repair Kit (Thermo Fisher Scientific, K0771); vectors were dephosphorylated with FastAP (Thermo Fisher Scientific, EF0654). Oligonucleotides were obtained from Eurofins Genomics and gBlocks from Integrated DNA Technologies. Gel and PCR products were purified using QIAEX II (QIAGEN, 20021) or NucleoSpin Gel and PCR Clean-up (MACHEREY-NAGEL, 740609), respectively.

Ligations were transformed into homemade chemically competent *Escherichia coli* DH5α; HiFi assemblies used competent cells from New England Biolabs (C2987). Plasmids were prepared using QIAGEN Mini-, Midi- or Maxi-prep kits (27104, 12243 and 12263); genome-editing preparations used endotoxin-removal buffer set 19048. Procedures followed the corresponding manufacturers' instructions. Final plasmids were verified by whole-plasmid sequencing (Plasmidsaurus or Eurofins Genomics). Final study constructs and Addgene accession numbers are provided in Supplementary Table 3.

##### Nucleases

The *hROSA26* Cas9-nickase/guide plasmids pSpCas9n(BB)\_R26-L and pSpCas9n(BB)\_R26-R were described previously<sup>50</sup>. AAVS1 ZFN and *CLYBL* TALEN plasmids were obtained from Addgene (159297/159298 and 62196/62197, respectively).

##### Renewal and transactivator cassettes

pR26-Bst\_CAG-Tet-On-3G was generated from pR26-Bst\_CAG-EGFPd2<sup>50</sup> by replacing EGFPd2 with synthetic Tet-On-3G cDNA using MluI (Thermo Fisher Scientific, FD0564) and BamHI (Thermo Fisher Scientific, FD0054).

pCLYBL-Neo\_CAG-NANOG\_2A\_mCherry was generated by cloning two synthetic fragments into BsrGI/MluI-cut pC13N-iCAG.copGFP (Addgene, 66578): a Bsp1407I/BamHI-cut fragment encoding human NANOG cDNA (RefSeq NM\_024865.4) and a BamHI/MluI-cut fragment encoding P2A-T2A-mCherry (Bsp1407I: Thermo Fisher Scientific, FD0933). The NANOG coding sequence differed from RefSeq NM\_024865.4 at seven positions: c.9G>A, c.165T>C, c.246G>T, c.276G>A, c.363C>T, c.531T>C and c.798C>T. Six substitutions were synonymous, whereas c.246G>T corresponds to the naturally occurring NANOG variant rs2889551 (p.Lys82Asn).

##### EGFP-linked miR-F validation vectors

pAAV-Puro\_TRE-EGFP, pAAV-Puro\_TRE-MYOD1 and pAAV-Puro\_TRE-NGN2 were synthesized based on published sequences<sup>16,50</sup>.

pAAV-Puro\_TRE-EGFP-(miR-F) was generated by cloning the miR-F cassette from LT11GFP\_Ren.713 (a kind gift of Johannes Zuber) into the 3' UTR of EGFP. The miR-F cassette was amplified as two fragments introducing paired AarI sites for subsequent hairpin insertion, and the two PCR products were assembled with EcoRI/EcoRV-cut pAAV-Puro\_TRE-EGFP (New England Biolabs, R3101/R3195) using HiFi assembly. The corresponding assembly oligonucleotides are listed below.

Complementary oligonucleotides encoding miR-F hairpins targeting CD151, CD46, RLuc or NANOG were annealed, phosphorylated and ligated into AarI-cut pAAV-Puro\_TRE-EGFP-(miR-F) (Thermo Fisher Scientific, ER1582) according to the previously described hairpin-insertion procedure<sup>50</sup>. Hairpin oligonucleotide sequences are provided in Supplementary Table 5.

##### MYOD1 UTR, intronic and bidirectional configurations

pAAV-Puro\_TRE-MYOD1-(miR-F-NANOG) was generated by transferring the anti-NANOG miR-F cassette from the corresponding EGFP vector into pAAV-Puro\_TRE-MYOD1 using EcoRI and EcoRV. The matched anti-CD151 control vector was generated analogously by transferring the anti-CD151 miR-F cassette from the corresponding EGFP vector into pAAV-Puro\_TRE-MYOD1 using the same restriction sites.

pAAV-Puro\_TRE-(intron-miR-F-NANOG)-MYOD1 was generated in three steps. First, a chimeric intron from pAAV-Puro\_siKO (Addgene, 86696) was excised with BssHII (New England Biolabs, R0199) and PacI (Thermo Fisher Scientific, FD2204), end-repaired and inserted into SpeI-cut and end-repaired pAAV-Puro\_TRE-MYOD1 (New England Biolabs, R3133). Second, the miR-F cassette was excised from pAAV-Puro\_TRE-EGFP-(miR-F) using EcoRV and NotI (Thermo Fisher Scientific, FD0593), end-repaired and inserted into the MreI-cut and end-repaired intermediate plasmid (Thermo Fisher Scientific, FD2024). Third, the NANOG hairpin was inserted into the AarI-cut miR-F cassette using the procedure described above.

pAAV-Puro\_TREb-EGFP-(miR-F-NANOG)\_MYOD1 was generated by HiFi assembly of two fragments into

SpeI/PspXI-cut pAAV-Puro\_TRE-MYOD1 (PspXI: New England Biolabs, R0656). The first insert comprised PCR-amplified EGFP and anti-NANOG miR-F from pAAV-Puro\_TRE-EGFP-(miR-F-NANOG); the second comprised the PCR-amplified bidirectional TRE3G promoter from the recorded pTRE3G-bi-alpha-chain donor (Addgene, 133730). The corresponding assembly oligonucleotides are listed below.

##### NGN2 and HFHR intronic configurations

pAAV-Puro\_TRE-(intron-miR-F-NANOG)-NGN2 was generated by HiFi assembly of the NEUROG2/NGN2 cDNA cassette PCR-amplified from pAAV-Puro\_TRE-NGN2 into the NcoI/KpnI-cut pAAV-Puro\_TRE-(intron-miR-F-NANOG)-MYOD1 backbone (Thermo Fisher Scientific, FD0574/FD0524). The corresponding assembly oligonucleotides are listed below.

pAAV-Puro\_TRE-(intron-miR-F-NANOG)-HNF1A-FOXA3-HNF6-RORC (Addgene, 260623) was generated by HiFi assembly of the *HNF1A-FOXA3-ONECUT1/HNF6-RORC* cassette from the published HFHR construct<sup>18</sup> into the NcoI/KpnI-cut pAAV-Puro\_TRE-(intron-miR-F-NANOG)-MYOD1 backbone. The four coding sequences were separated, in order, by P2A, T2A and E2A self-cleaving peptide sequences. The corresponding assembly oligonucleotides are listed below.

##### Assembly oligonucleotides

The sequences below are written 5' to 3'. They comprise the oligonucleotides used for the assembly steps described above; miR-F hairpin, genotyping and RT-qPCR oligonucleotides are reported separately in the corresponding Supplementary Tables.

| Assembly | Primer | Sequence (5' to 3') |
| --- | --- | --- |
| miR-F fragment 1 | F | CATGGACGAGCTGTACAAGTAAGCGTACGGCGGCCGCCTC |
|  | R | CAGGTGAGTCAGTCCACCTGCACCACGCTCACTGCCAACAGC |
| miR-F fragment 2 | F | CAGGTGGACTGACTCACCTGCACCTTGCCTACTGCCTCGG |
|  | R | GATTATGATCCTCTAGAGATATCAATTACGCGTCAATTGATGCATTC |
| BID EGFP-miR-F | F | GTGAACCGTCAGATCGCCTAGTGCCACCATGGTGAGCAAGG |
|  | R | GTGGGCTCTATGGTCACTCGAATGCATTCAGCTTTG |
| BID TRE3G | F | CCTTGCTCACCATGGTGGCACTAGGCGATCTGACGGTTCAC |
|  | R | ACAGTAGCTCCATGGTGGCACTAGTTTACGAGGGTAGGAAG |
| NEUROG2/NGN2 | F | TTACTAGTGCCACCA<sup>t</sup>gTTCGTCAAATCCGAGACCTTGGAG |
|  | R | GGATCCCCGGGTACCGAGCTCGAATTCCTAGATACAATCC |
| HFHR | F | TTACTAGTGCCACCATGGTTTCTAAACTGAG |
|  | R | AACAACAACAATTGCATTATTTATGTTTCAGgttc |

#### Supplementary Tables

| Cell ID used in paper | Source / registry ID | Genotype / engineered features | Genomic characterization |
| --- | --- | --- | --- |
| WT | WTC11; hPSCreg UCSFi001-A; RRID:CVCL_Y803; Coriell GM25256 | Unmodified parental WTC11 hiPSC line | SNP array (B,F,H,J,K) |
| E | WTC11 derivative; AICS-0048-039; hPSCreg UCSFi001-A-27; RRID:CVCL_UD16 | Monoallelic C-terminal *TTN*-mEGFP reporter | G-banding from producer (https://www.coriell.org/0/PDF/Allen/ipsc/AICS-0048-039_CofA.pdf) |
| ET | E | E + monoallelic *hROSA26* CAG-Tet-On-3G | Targeted integration PCR |
| ETN<sub>1</sub> | ET | ET + monoallelic *CLYBL*-targeted *NANOG*-2A-mCherry | Targeted integration PCR |
| ETN<sub>2</sub> | ET | ET + biallelic *CLYBL*-targeted *NANOG*-2A-mCherry | Targeted integration PCR; chromosomal microarray analysis (CMA) |
| ETM / OPTi-OX MYOD1 | ET | ET + biallelic *AAVS1*-targeted dox-inducible *MYOD1* | Targeted integration PCR; SNP array (B,F,H,K) |
| ETN<sub>2</sub>MR<sub>C-UTR</sub>, clone 1 | ETN<sub>2</sub> | ETN<sub>2</sub> + biallelic *AAVS1*-targeted dox-inducible *MYOD1* with 3'-UTR anti-CD151 control miR-F | Targeted integration PCR |
| ETN<sub>2</sub>MR<sub>C-UTR</sub>, clone 2 | ETN<sub>2</sub> | As clone 1, except monoallelic *AAVS1* targeting | Targeted integration PCR |
| ETN<sub>2</sub>MR<sub>C-UTR</sub>, clone 3 | ETN<sub>2</sub> | As clone 1 | Targeted integration PCR |
| ETN<sub>2</sub>MR<sub>N-UTR</sub>, clone 1 | ETN<sub>2</sub> | ETN<sub>2</sub> + biallelic *AAVS1*-targeted dox-inducible *MYOD1* with 3'-UTR anti-*NANOG* miR-F | Targeted integration PCR; SNP array (A,B,F,H,I,J,K) |
| ETN<sub>2</sub>MR<sub>N-UTR</sub>, clone 2 | ETN<sub>2</sub> | As clone 1 | Targeted integration PCR |
| ETN<sub>2</sub>MR<sub>N-UTR</sub>, clone 3 | ETN<sub>2</sub> | As clone 1 | Targeted integration PCR |
| ETN<sub>2</sub>MR<sub>N-BID</sub>, clone 1 | ETN<sub>2</sub> | ETN<sub>2</sub> + biallelic *AAVS1*-targeted dox-inducible *MYOD1* with bidirectional EGFP-anti-*NANOG* miR-F cassette | Targeted integration PCR; SNP array (A,B,F,H,J,K) |
| ETN<sub>2</sub>MR<sub>N-BID</sub>, clone 2 | ETN<sub>2</sub> | As clone 1 | Targeted integration PCR |
| ETN<sub>2</sub>MR<sub>N-BID</sub>, clone 3 | ETN<sub>2</sub> | As clone 1 | Targeted integration PCR |
| GERALT MYOD1 / ETN<sub>2</sub>MR<sub>N-INT</sub>, clone 1 | ETN<sub>2</sub> | ETN<sub>2</sub> + biallelic *AAVS1*-targeted dox-inducible *MYOD1* with intronic anti-*NANOG* miR-F | Targeted integration PCR; SNP array (B,F,H,I,J,K) |
| ETN<sub>2</sub>MR<sub>N-INT</sub>, clone 2 | ETN<sub>2</sub> | As clone 1 | Targeted integration PCR |
| ETN<sub>2</sub>MR<sub>N-INT</sub>, clone 3 | ETN<sub>2</sub> | As clone 1 | Targeted integration PCR |
| GERALT NGN2, clone 1 | ETN<sub>2</sub> | ETN<sub>2</sub> + biallelic *AAVS1*-targeted dox-inducible *NEUROG2/NGN2* with intronic anti-*NANOG* miR-F | Targeted integration PCR; SNP array (B,F,H,K) |
| GERALT NGN2, clone 2 | ETN<sub>2</sub> | As clone 1 | Targeted integration PCR; SNP array (B,F,H,K) |
| GERALT NGN2, clone 3 | ETN<sub>2</sub> | As clone 1 | Targeted integration PCR; SNP array (E,G,K) |
| GERALT HFHR, clone 1 | ETN<sub>2</sub> | ETN<sub>2</sub> + biallelic *AAVS1*-targeted dox-inducible *HNF1A-FOXA3-ONECUT1/HNF6-RORC* cassette with intronic anti-*NANOG* miR-F | Targeted integration PCR; SNP array (B,F,H,K) |
| GERALT HFHR, clone 2 | ETN<sub>2</sub> | As clone 1 | Targeted integration PCR; SNP array (B,G,H,K) |
| GERALT HFHR, clone 3 | ETN<sub>2</sub> | As clone 1 | Targeted integration PCR; SNP array (B,C,D,F,H,K) |

Continued on next page

| Cell ID used in paper | Source / registry ID | Genotype / engineered features | Genomic characterization |
| --- | --- | --- | --- |
| OPTi-OX HFHR | FSPS13B; HipSci<br>HPSI0813i-fpdm_2;<br>ECACC 77650622;<br>RRID:CVCL_AH05 | Monoallelic *hROSA26* CAG–Tet-On-3G + biallelic<br>*AAVS1*-targeted dox-inducible<br>*HNF1A-FOXA3-ONECUT1/HNF6-RORC* | Targeted integration PCR;<br>StemGenomics iCS-digital PSC<br>24-probe CNV assay: 20q11 gain<br>(also detected in earliest available<br>parental stock) |

###### Supplementary Table 1 | Cell lines

Parental, reporter and engineered hiPSC lines used in the study, with source or registry identifiers, engineered genomic features and the genomic characterization performed for each line or clone. A dash indicates that no additional genomic characterization was performed or was available for the indicated line. **SNP-array alteration key (GRCh37/hg19; approximate intervals in Mb).** **A**, chr2 gain (179.3–179.9); **B**, chr2 copy-neutral loss of heterozygosity (LoH; 192.7–198.2); **C**, chr6 loss (0.1–33.2); **D**, chr6 gains (33.1–171.0; discontinuous); **E**, chr7 gain (6.3–20.2); **F**, chr7 loss (69.2–70.2); **G**, chr7 losses (61.0–159.2; discontinuous); **H**, chr12 copy-neutral LoH (39.1–41.8); **I**, chr16 loss (73.7–73.9); **J**, chr22 loss (25.6–26.0); **K**, chrY loss (6.0–9.2). Intervals are conservatively rounded outwards and summarize overlapping or concordant calls. For complex abnormalities (D and G), outer coordinates encompass the called segments but do not imply continuous alteration. Short gaps between chrY calls were consolidated. Codes apply only to SNP-array-profiled lines and should not be interpreted as genomic characterization of untested derivatives. The OPTi-OX HFHR 20q11 gain was detected by a separate targeted assay. The complete set of filtered SNP-array calls, including exact coordinates and confidence scores, is available on Zenodo, as detailed in the Data Availability section.

| Medium | Basal medium | Supplements / final concentrations | Use |
| --- | --- | --- | --- |
| Commercial E8 | Essential 8 (Thermo Fisher Scientific, A1517001) | Ready-to-use commercial medium. | Initial hiPSC maintenance; base medium for CLYBL nucleofection recovery. |
| E8 | Essential 6 (Thermo Fisher Scientific, A1516401) | 100 ng mL<sup>-1</sup> FGF2-145aa (Qkine, Qk025); 2 ng mL<sup>-1</sup> TGFβ1 (Qkine, Qk010). | TGFβ-replete comparator in pathway-withdrawal and inhibition assays. |
| E7 | Essential 6 | 100 ng mL<sup>-1</sup> FGF2-145aa. | TGFβ-free condition in pathway-withdrawal assays. |
| E7+SB | Essential 6 | 100 ng mL<sup>-1</sup> FGF2-145aa; 10 μM SB431542 (Selleck Chemicals, S1067). | TGFβ pathway-inhibition condition. |
| hE8 | DMEM/F-12 with L-glutamine and HEPES (Gibco, 31330038) | 64 μg mL<sup>-1</sup> 2-phospho-L-ascorbic acid trisodium salt (Sigma-Aldrich, 49752-10G); 14 ng mL<sup>-1</sup> sodium selenite (Sigma-Aldrich, S5261-25G); 1,743 μg mL<sup>-1</sup> sodium bicarbonate (Sigma-Aldrich, S8875-500G); 10 μg mL<sup>-1</sup> Optiferrin (Invitria, 777TRF029-1G); 19.4 μg mL<sup>-1</sup> insulin (Gibco, A11382II); 5 ng mL<sup>-1</sup> t-FGF2-G3 (145aa; Qkine, Qk052); 2 ng mL<sup>-1</sup> TGFβ1<sup>30</sup>. | Parental/control hiPSC maintenance; base medium for GERALT HFHR nucleofection recovery. |
| hE7 | DMEM/F-12 with L-glutamine and HEPES | hE8 formulation lacking TGFβ1. | NANOG-overexpressing/GERALT maintenance, suspension culture and minimal-medium forward programming. |
| hE6 | DMEM/F-12 with L-glutamine and HEPES | hE8 formulation lacking TGFβ1 and FGF2. | MYOD1, NGN2 and hepatic forward programming. |
| CEPT recovery medium | Commercial E8 or hE8, as indicated | 1:1,000 polyamine supplement (Sigma-Aldrich, P8483); 50 nM Chroman 1 (MedChemExpress, HY-15392); 5 μM emricasan (Sigma-Aldrich, SML2227); 0.7 μM trans-ISRIB (Sigma-Aldrich, SML0843)<sup>51</sup>. | Recovery after CLYBL nucleofection (commercial E8) or GERALT HFHR AAVS1 nucleofection (hE8). |
| Myogenic medium (MYOD1) | High-glucose DMEM with GlutaMAX (Thermo Fisher Scientific, 61965026) | 1× ITS-G (Thermo Fisher Scientific, 41400045); 40 ng mL<sup>-1</sup> FGF2-145aa (Qkine, Qk025); 1 μM retinoic acid (Sigma-Aldrich, R2625); 3 μM CHIR99021 (Cayman Chemical, 13122)<sup>16</sup>. | MYOD1 forward programming. |
| Stepwise myogenic differentiation media | DMEM/F-12 (Gibco, 11320033) | 1× non-essential amino acids (Thermo Fisher Scientific, 11140050); 1× ITS-G; 3 μM CHIR99021; 500 nM LDN193189 (Miltenyi Biotec, 130-106-540); 20 ng mL<sup>-1</sup> FGF-2 (R&D Systems, 233-FB/CF); 15% KnockOut Serum Replacement (Gibco, 10828028); 2 ng mL<sup>-1</sup> IGF1 (R&D Systems, 291-G1-200); 10 ng mL<sup>-1</sup> HGF (R&D Systems, 294-HGN-100/CF). | 28-d stepwise differentiation of WTC11 hiPSCs into myoblasts; timing of supplement addition and withdrawal is described in Methods. |
| SKGM | High-glucose DMEM (Thermo Fisher Scientific, 11965092) | 1 mM sodium pyruvate (Thermo Fisher Scientific, 11360070); 10% heat-inactivated FBS (Thermo Fisher Scientific, 10270106); 1 μM dexamethasone (Sigma-Aldrich, D4902); 40 ng mL<sup>-1</sup> FGF2-145aa; 100 nM human insulin. | Expansion of stepwise hiPSC-derived myoblasts. |
| Myoblast fusion medium | High-glucose DMEM + sodium pyruvate | 2% horse serum (Thermo Fisher Scientific, 16050114); 1× B27 (Thermo Fisher Scientific, 17504044); 1× non-essential amino acids; 10 ng mL<sup>-1</sup> IGF1; 10 ng mL<sup>-1</sup> HGF; 10 μM SB431542. | Terminal differentiation/fusion of stepwise hiPSC-derived myoblasts. |
| SkGM-2 | SkGM-2 BulletKit (Lonza, CC-3245) | Manufacturer formulation. | HSMM expansion. |
| HSMM fusion medium | DMEM/F-12 (Gibco, 11320033) | 2% horse serum. | Terminal differentiation of primary HSMM. |
| N2 | DMEM/F-12 | 1× N2 (Thermo Fisher Scientific, 17502048); 1× non-essential amino acids; 20 μg mL<sup>-1</sup> insulin (Sigma-Aldrich, I9278); 1× penicillin-streptomycin (Thermo Fisher Scientific, 15140122); plus doxycycline during NGN2 induction. | NGN2 neuronal forward programming; viral controls additionally receive puromycin. |
| HLI | HepatoZYME-SFM (Thermo Fisher Scientific, 17705021) | 1% penicillin-streptomycin; 2% MEM non-essential amino acids; 0.5% chemically defined lipid concentrate (Gibco, 11905031); 2 μg mL<sup>-1</sup> human insulin; 30 μg mL<sup>-1</sup> Optiferrin. Benchmark condition uses 2% lipid concentrate. | Hepatic differentiation after day 2. |

Continued on next page

| Medium | Basal medium | Supplements / final concentrations | Use |
| --- | --- | --- | --- |
| EMT post-casting medium, direct hiPSC | hE6 | 2 g L<sup>-1</sup> 6-aminocaproic acid (Sigma-Aldrich, A2504); 1 µg mL<sup>-1</sup> doxycycline; 2 µM thiazovivin. | First 24 h after casting GERALT directly from hiPSCs. |
| EMT post-casting medium, pre-induced | SKGM | 2 g L<sup>-1</sup> 6-aminocaproic acid; 1 µg mL<sup>-1</sup> doxycycline; 2 µM thiazovivin. Myoblast condition omits doxycycline. | First 24 h after casting pre-induced GERALT or myoblasts. |
| EMT differentiation medium | High-glucose DMEM | 2% horse serum; 1× B27; 1× non-essential amino acids; 10 ng mL<sup>-1</sup> IGF1; 10 ng mL<sup>-1</sup> HGF; 10 µM SB431542; 2 g L<sup>-1</sup> 6-aminocaproic acid; 0.5% penicillin–streptomycin; doxycycline maintained for GERALT only. | Long-term culture of engineered muscle tissues. |

**Supplementary Table 2 | Composition and experimental use of cell culture media**

Basal media, supplements and final concentrations used for hiPSC maintenance, pathway-perturbation experiments, lineage forward programming, myoblast expansion and fusion, suspension culture and engineered muscle tissue generation. The principal experimental application of each formulation is indicated.

| Plasmid name | Purpose | Addgene accession |
| --- | --- | --- |
| pR26-Bst_CAG-Tet-On-3G | *hROSA26* donor plasmid encoding Tet-On 3G under the CAG promoter. | 260616 |
| pSpCas9n(BB)_R26-L | SpCas9 nickase/gRNA plasmid targeting the left *hROSA26* site for paired-nickase genome editing. | 255612 |
| pSpCas9n(BB)_R26-R | SpCas9 nickase/gRNA plasmid targeting the right *hROSA26* site for paired-nickase genome editing. | 255613 |
| pCLYBL-Neo_CAG-NANOG_2A_mCherry | *CLYBL* donor plasmid encoding CAG-driven human *NANOG*-P2A-T2A-mCherry. | 260617 |
| pZT-C13-L1 | Left TALEN expression plasmid targeting the *CLYBL* locus. | 62196 |
| pZT-C13-R1 | Right TALEN expression plasmid targeting the *CLYBL* locus. | 62197 |
| pAAV-Puro_TRE-EGFP-(miR-F) | *AAVS1* donor backbone for doxycycline-inducible EGFP expression followed by a 3'-UTR miR-F cloning scaffold; used to generate the CD151-, CD46-, RLuc- and *NANOG*-targeting miR-F validation vectors. | 260618 |
| pAAV-Puro_TRE-MYOD1-(miR-F-NANOG) | *AAVS1* donor plasmid encoding doxycycline-inducible human *MYOD1* followed in the 3' UTR by a miR-F artificial microRNA targeting *NANOG*, enabling coupled myogenic induction and *NANOG* silencing. | 260619 |
| pAAV-Puro_TRE-(intron-miR-F-NANOG)-MYOD1 | *AAVS1* donor plasmid encoding doxycycline-inducible human *MYOD1* with an intronic miR-F artificial microRNA targeting *NANOG*; lead GERALT MYOD1 configuration. | 260620 |
| pAAV-Puro_TREb-EGFP-(miR-F-NANOG)_MYOD1 | *AAVS1* donor plasmid with a bidirectional doxycycline-inducible TRE promoter driving EGFP-miR-F-NANOG on one side and human *MYOD1* on the opposite side. | 260621 |
| pAAV-Puro_TRE-(intron-miR-F-NANOG)-NGN2 | *AAVS1* donor plasmid encoding doxycycline-inducible human *NEUROG2/NGN2* with an intronic miR-F artificial microRNA targeting *NANOG*, enabling coupled neuronal induction and *NANOG* silencing. | 260622 |
| pAAV-Puro_TRE-(intron-miR-F-NANOG)-HNF1A-FOXA3-HNF6-RORC | *AAVS1* donor plasmid encoding doxycycline-inducible human *HNF1A-FOXA3-HNF6-RORC* with an intronic miR-F artificial microRNA targeting *NANOG*. | 260623 |
| pZFN-AAVS1_ELD | Left ZFN expression plasmid targeting the *AAVS1* locus with obligate-heterodimer FokI. | 159297 |
| pZFN-AAVS1_KKR | Right ZFN expression plasmid targeting the *AAVS1* locus with obligate-heterodimer FokI. | 159298 |

**Supplementary Table 3 | Plasmids and genome-engineering constructs deposited for this study**

Donor and nuclease plasmids deposited with Addgene for generation of the renewal and differentiation circuits, with their experimental purpose and accession number. Additional source and intermediate constructs used for cloning are described in Supplementary Note 2.

| Locus | PCR type | Primer | Sequence (5' to 3') | Annealing temp. (°C) | Extension time | Expected amplicon size |  |  |
| --- | --- | --- | --- | --- | --- | --- | --- | --- |
|  |  |  |  |  |  | Control gDNA | Targeted allele | Donor plasmid |
| *hROSA26* | WT locus | F<br>R | GAGAAGAGGCTGTGCTTCGG<br>ACAGTACAAGCCAGTAATGGAG | 63 | 3 min | 2186 bp | No band | No band |
|  | 5' INT | F<br>R | GAGAAGAGGCTGTGCTTCGG<br>AAGACCGCGAAGAGTTTGTC | 60 | 1 min 30 s | No band | 1274 bp | No band |
|  | 3' INT | F<br>R | GAGAATAGCAGGCATGCTG<br>ACAGTACAAGCCAGTAATGGAG | 60 | 1 min 30 s | No band | 1020 bp | No band |
|  | 5' BB | F<br>R | CGTTGTAAAACGACGCCAG<br>AAGACCGCGAAGAGTTTGTC | 60 | 1 min 30 s | No band | No band | 1035 bp |
|  | 3' BB | F<br>R | GAGAATAGCAGGCATGCTG<br>TGACCATGATTACGCCAAGC | 60 | 1 min 30 s | No band | No band | 919 bp |
| *CLYBL* | WT locus | F<br>R | TAAGTGACCCCTGGCGAGAC<br>AGATGGAGCAGTGGATGACA | 65 | 2 min | 1728 bp | No band | No band |
|  | 5' INT | F<br>R | TAAGTGACCCCTGGCGAGAC<br>AAGACTTCCTCTGCCCTCTC | 65 | 2 min | No band | 962 bp | No band |
|  | 3' INT | F<br>R | TCCAGGACGGAGTCAGTGAG<br>AGATGGAGCAGTGGATGACA | 60 | 2 min | No band | 907 bp | No band |
|  | 5' BB | F<br>R | CACAGGAAACAGCTATGACC<br>AAGACTTCCTCTGCCCTCTC | 60 | 2 min | No band | No band | 980 bp |
|  | 3' BB | F<br>R | TCCAGGACGGAGTCAGTGAG<br>TTTTCCCAGTCACGACGTTG | 65 | 2 min | No band | No band | 874 bp |
| *AAVS1* | WT locus | F<br>R | CTGTTTCCCCTTCCCAGGCAGGTCC<br>TGCAGGGGAACGGGCTCAGTCTGA | 65 | 2 min | 1692 bp | No band | No band |
|  | 5' INT | F<br>R | CTGTTTCCCCTTCCCAGGCAGGTCC<br>TCGTCGCGGGTGGCGAGGCGCACCG | 65 | 1 min 30 s | No band | 994 bp | No band |
|  | 3' INT | F<br>R | GAATTCGAGCTCGGTACC<br>TGCAGGGGAACGGGCTCAGTCTGA | 65 | 1 min 30 s | No band | 1168 bp | No band |
|  | 5' BB | F<br>R | GCTTGCTGTAAAGCGGATGC<br>TCGTCGCGGGTGGCGAGGCGCACCG | 60 | 1 min 30 s | No band | No band | 1424 bp |
|  | 3' BB | F<br>R | GAATTCGAGCTCGGTACC<br>ATGCTTCCGGCTCGTATGTT | 55 | 1 min 30 s | No band | No band | 1227 bp |

**Supplementary Table 4 | Genotyping PCR primers, amplification conditions and expected products**

Primer sequences and PCR conditions used to assess unmodified loci, 5' and 3' targeted-integration junctions and donor-backbone integration at *hROSA26*, *CLYBL* and *AAVS1*. Expected amplicon sizes are reported for control genomic DNA, correctly targeted clones and donor-plasmid controls. INT, targeted-integration junction PCR; BB, donor-backbone PCR.

| Seed target | Sequence (5' to 3') |
| --- | --- |
| Human *CD46* | AGCGCAGCCAAGCAGTTTCTTTCAAATAGTGAAGCCACAGATGTATTTGAAAGAAACTGCTTGGCTA<br>GGCATAGCCAAGCAGTTTCTTTCAAATACATCTGTGGCTTCACTATTTGAAAGAAACTGCTTGGCTG |
| Human *CD151* | AGCGCGCCTCAAGTACCTGCTGTTTATAGTGAAGCCACAGATGTATAAACAGCAGGTACTTGAGGCA<br>GGCATGCCTCAAGTACCTGCTGTTTATACATCTGTGGCTTCACTATAAACAGCAGGTACTTGAGGCG |
| Renilla luciferase | AGCGCAGGAATTATAATGCTTATCTATAGTGAAGCCACAGATGTATAGATAAGCATTATAATTCCTA<br>GGCATAGGAATTATAATGCTTATCTATACATCTGTGGCTTCACTATAGATAAGCATTATAATTCCTG |
| Human *NANOG* | AGCGCTCCGACTGTAAAGAATCTTCATAGTGAAGCCACAGATGTATGAAGATTCTTTACAGTCGGAT<br>GGCAATCCGACTGTAAAGAATCTTCATACATCTGTGGCTTCACTATGAAGATTCTTTACAGTCGGAG |

**Supplementary Table 5 | miR-F hairpin oligonucleotide sequences**

Oligonucleotide sequences used to generate miR-F artificial microRNAs targeting human *CD46*, *CD151* and *NANOG*, together with the Renilla luciferase control hairpin. Sequences are shown in the 5'-to-3' orientation.

| Target / assay | Primer | Primer sequence (5' to 3') or assay identifier |
| --- | --- | --- |
| *ACTA1* | F | CCCGCCCAGAAACTAGACAC |
|  | R | GTCGCCCACGTAGGAATCTT |
| *AFP* | F | TGCGGCCTCTTCCAGAAACT |
|  | R | TAATGTCAGCCGCTCCCTCG |
| *ALB* | F | CCTTTGGCACAATGAAGTGGGTAACC |
|  | R | CAGCAGTCAGCCATTTCACCATAG |
| *APOA1* | F | AGACAGCGGCAGAGACTATG |
|  | R | CCAGTTGTCAAGGAGCTTTAGG |
| *CYP3A4* | F | TGTGCCTGAGAACACCAGAG |
|  | R | GTGGTGGAAATAGTCCCGTG |
| *DPPA4* | F | ATGGAGAAGGCAAAAGGCAA |
|  | R | GGGTTGTCAGTGTGCTCTGC |
| *GBX2* | F | GTTCCACTGCAAAAAGTACCTCT |
|  | R | GGGACGACGATCTTAGGGTTC |
| *HMBS/PBGD* | F | GGAGCCATGTCTGGTAACGG |
|  | R | CCACGCGAATCACTCTCATCT |
| *HNFI1A* | F | TGGCCATGGACACGTACAG |
|  | R | GCTGCTTGAGGGTACTTCTG |
| *HPRT1/HPRT* | F | TGACACTGGCAAAACAATGCA |
|  | R | GGTCCTTTTCACCAGCAAGCT |
| *LEFTY1* | F | CGCCAGGAGATGTACATTGA |
|  | R | CTTGATGCTGACGATCATGG |
| *MAP2* | F | AGACTGCAGCTCTGCCTTTAG |
|  | R | AGGCTGTAAGTAAATCTTCCTCC |
| *MYBPC2* | F | CCTATTGCACCCACGAGTGA |
|  | R | CTCCGGTCGGGAGATTCTTG |
| *MYH3* | F | TGCTGCATACCCAGAACACC |
|  | R | CCCTGCTGGCATCTTCTACC |
| *MYOG* | F | TGGGCGTGTAAGGTGTGTAA |
|  | R | CGATGTACTGGATGGCACTG |
| *NANOG, endogenous* | F | ATGCAACCTGAAGACGTGTGA |
|  | R | ACCATACTCCACCCTCCATGA |
| *NEUROG2/NGN2* | F | TGTTCGTCAAATCCGAGACCT |
|  | R | CGATCCGAGCAGCACTAACA |
| *NODAL* | F | TGAGCCAACAAGAGGATCTG |
|  | R | TGGAAAATCTCAATGGCAAG |
| *PAX6* | F | CTTTGCTTGGGAAATCCGAG |
|  | R | AGCCAGGTTGCGAAGAACTC |
| *POU3F2/BRN2* | F | ACCCGCTTTATCGAAGGCAA |
|  | R | CCTCCATAACCTCCCCCAGA |
| *POU5F1/OCT4* | F | AGTGAGAGGCAACCTGGAGA |
|  | R | ACACTCGGACCACATCCTTC |
| *RPLP0* | F | GGCGTCCTCGTGGAAGTGAC |
|  | R | GCCTTGCGCATCATGGTGTT |
| *SERPINA1/A1AT* | F | CCACCGCCATCTTCTTCCTGCCTGA |
|  | R | GAGCTTCAGGGGTGCCTCCTCTG |
| *SLC17A6/VGLUT2* | F | GTAGACTGGCAACCACCTCC |
|  | R | CCATTCCAAAGCTTCCGTAGAC |
| *SOX1* | – | QIAGEN QuantiTect Primer Assay, QT00215299 |
| *SYN* | F | ACCTCGGGACTCAACACCTCGG |
|  | R | GAACCACAGGTTGCCGACCCAG |
| *TNNI1* | F | ACCATGCCGGAAGTCGAGAG |
|  | R | TAGCGCACCTTCTCAGCCTC |
| *TTN* | F | GTAAAAAGAGCTGCCCCAGTGA |
|  | R | GCTAGGTGGCCCAGTGCTACT |

Continued on next page

| Target / assay | Primer | Primer sequence (5' to 3') or assay identifier |
| --- | --- | --- |
| *Transgene-specific assays* |  |  |
| HFHR transgene, *FOXA3*-T2A junction | F | GCCTCTATTCCCGCTCTTTGC |
|  | R | ATTCTCCTCCACGTCACCGC |
| *NANOG*<sub>exo</sub> transgene | F | GGGCAACGTGCTGGTTATT |
|  | R | CAGTCTCCGTGTGAGGCAT |

**Supplementary Table 6 | RT-qPCR primers and commercial primer assays**

Primer sequences and commercial primer-assay identifiers used for RT-qPCR analysis of endogenous genes and engineered transgenes. Official human gene symbols are given first, followed by aliases used in the manuscript where applicable. The endogenous and transgenic *NANOG* assays are specific for the 3' UTR and 5' UTR of these transcripts, respectively. HFHR-specific assays target junctions within the polycistronic *HNF1A*-P2A-*FOXA3*-T2A-*ONECUT1*/*HNF6*-E2A-*RORC* transgene. Primer sequences are shown in the 5'-to-3' orientation.

| Target / antibody | Host | Clone /<br>clonality | Conjugate | Manufacturer | Catalogue no. | Working dilution /<br>concentration |
| --- | --- | --- | --- | --- | --- | --- |
| **Flow cytometry** |  |  |  |  |  |  |
| *Primary antibodies* |  |  |  |  |  |  |
| CD151 | Mouse | 50-6 | APC | BioLegend | 350405 | 1:50 |
| CD46 | Mouse | TRA-2-10 | APC | BioLegend | 352405 | 1:50 |
| NANOG | Mouse | N31-355 | Alexa Fluor<br>647 | BD Biosciences | 561300 | 1:50 |
| OCT4 | Mouse | 40/Oct-3 | PE | BD Biosciences | 560186 | 1:15 |
| *Isotype controls* |  |  |  |  |  |  |
| Mouse IgG1κ isotype<br>control | Mouse | MOPC-21 | APC | BioLegend | 400121 | Matched to test antibody |
| NANOG isotype control | Mouse | MOPC-21 | Alexa Fluor<br>647 | BD Biosciences | 557714 | Matched to test antibody |
| OCT4 isotype control | Mouse | MOPC-21 | PE | BD Biosciences | 556650 | Matched to test antibody |
| **Western blotting** |  |  |  |  |  |  |
| *Primary antibodies* |  |  |  |  |  |  |
| NANOG | Goat | Polyclonal | – | R&D<br>Systems/Bio-Techne | AF1997 | 1 $\mu\text{g mL}^{-1}$ |
| VCL/vinculin | Mouse | V284 | – | Sigma-Aldrich | SAB4200080 | 0.01 $\mu\text{g mL}^{-1}$ |
| *Secondary antibodies* |  |  |  |  |  |  |
| Goat anti-mouse IgG<br>(H+L) | Goat | Polyclonal | HRP | Bio-Rad | 1706516 | 1:3,000 |
| Mouse anti-goat IgG | Mouse | Monoclonal | HRP | Santa Cruz<br>Biotechnology | sc-2354 | 0.4 $\mu\text{g mL}^{-1}$ |
| **Immunofluorescence** |  |  |  |  |  |  |
| *Primary antibodies* |  |  |  |  |  |  |
| ACTN2/sarcomeric<br>α-actinin | Mouse | EA-53 | – | Abcam | ab9465 | 1:100 |
| ACTN2/α-actinin 2 | Rabbit | 2285D | – | R&D<br>Systems/Bio-Techne | MAB9830 | 2.5 $\mu\text{g mL}^{-1}$ |
| ALB/albumin | Goat | Polyclonal | – | Bethyl Laboratories | A80-229A | 1:100 |
| HNF1A | Mouse | 2 | – | Santa Cruz<br>Biotechnology | sc-135939 | 2 $\mu\text{g mL}^{-1}$ |
| HNF4A | Rabbit | EPR3648 | – | Abcam | ab92378 | 1:150 |
| MAP2 | Chicken | Polyclonal | – | Abcam | ab5392 | 1:5,000 |
| MYOD1/MyoD1 | Rabbit | EPR6653-131 | – | Abcam | ab133627 | 1:250 |
| NANOG | Goat | Polyclonal | – | R&D<br>Systems/Bio-Techne | AF1997 | 2 $\mu\text{g mL}^{-1}$ |
| Sarcomeric myosin heavy<br>chain (MHC) | Mouse | MF20 | – | DSHB | MF20 | 1 $\mu\text{g mL}^{-1}$ |
| SOX2 | Rabbit | Polyclonal | – | Abcam | ab97959 | 1:150 |
| TUBB3/β-tubulin III | Mouse | SDL3D10 | – | Sigma-Aldrich | T8660 | 1:1,000 |
| *Secondary antibodies* |  |  |  |  |  |  |
| Donkey anti-goat IgG<br>(H+L) | Donkey | Polyclonal | Alexa Fluor<br>488 | Thermo Fisher<br>Scientific/Invitrogen | A11055 | 4 $\mu\text{g mL}^{-1}$ |
| Donkey anti-goat IgG<br>(H+L) | Donkey | Polyclonal | Alexa Fluor<br>568 | Thermo Fisher<br>Scientific/Invitrogen | A11057 | 4 $\mu\text{g mL}^{-1}$ |
| Donkey anti-rabbit IgG<br>(H+L) | Donkey | Polyclonal | Alexa Fluor<br>488 | Thermo Fisher<br>Scientific/Invitrogen | A21206 | 4 $\mu\text{g mL}^{-1}$ |
| Donkey anti-rabbit IgG<br>(H+L) | Donkey | Polyclonal | Alexa Fluor<br>546 | Thermo Fisher<br>Scientific/Invitrogen | A10040 | 4 $\mu\text{g mL}^{-1}$ |
| Donkey anti-rabbit IgG<br>(H+L) | Donkey | Polyclonal | Alexa Fluor<br>647 | Thermo Fisher<br>Scientific/Invitrogen | A31573 | 4 $\mu\text{g mL}^{-1}$ |
| Goat anti-chicken IgY<br>(H+L) | Goat | Polyclonal | Alexa Fluor<br>647 | Thermo Fisher<br>Scientific/Invitrogen | A21449 | 4 $\mu\text{g mL}^{-1}$ |
| Goat anti-mouse IgG<br>(H+L) | Goat | Polyclonal | Alexa Fluor<br>488 | Thermo Fisher<br>Scientific/Invitrogen | A11001 | 4 $\mu\text{g mL}^{-1}$ |
| Goat anti-mouse IgG<br>(H+L) | Goat | Polyclonal | Alexa Fluor<br>568 | Thermo Fisher<br>Scientific/Invitrogen | A11004 | 4 $\mu\text{g mL}^{-1}$ |

Continued on next page

| Target / antibody | Host | Clone /<br>clonality | Conjugate | Manufacturer | Catalogue no. | Working dilution /<br>concentration |
| --- | --- | --- | --- | --- | --- | --- |
| Goat anti-rabbit IgG<br>(H+L) | Goat | Polyclonal | Alexa Fluor<br>488 | Thermo Fisher<br>Scientific/Invitrogen | A11008 | 4 $\mu\text{g mL}^{-1}$ |

**Supplementary Table 7 | Antibodies used for flow cytometry, immunoblotting and immunofluorescence**

Primary antibodies, isotype controls and secondary antibodies used in this study, including host species, clone or clonality, conjugate, manufacturer, catalogue number and working dilution or concentration. Within each application, primary antibodies and controls or secondary antibodies are listed separately and ordered alphabetically by target or antibody name. Official protein symbols are given first, followed by aliases used in the manuscript where applicable.

| Submitted ID | Description | Cell line | Medium | Stage | Rep. | Raw pairs | Trimmed pairs | Retained (%) | R1 adapter (%) | R2 adapter (%) | Unique (%) | Multi-map (%) | Too short (%) | Mis-match (%) | Assigned (%) | Genes detected |
| --- | --- | --- | --- | --- | --- | --- | --- | --- | --- | --- | --- | --- | --- | --- | --- | --- |
| **E-MTAB-17410** |  |  |  |  |  |  |  |  |  |  |  |  |  |  |  |  |
| c16_TGF_8d_D1 | ET parental WTC11 hiPSC line | ET | E8 | D8 | R1 | 59590979 | 59542513 | 100% | 38% | 34% | 83% | 5% | 13% | 0% | 73% | 29287 |
| c118_TGF_8d_D1 | NANOG<sub>exo</sub> cassette | ETN<sub>2</sub> | E8 | D8 | R1 | 37136873 | 37121337 | 100% | 38% | 36% | 83% | 5% | 12% | 0% | 72% | 26213 |
| c16_SB_8d_D1 | ET parental WTC11 hiPSC line | ET | E7 + SB | D8 | R1 | 50806844 | 50764223 | 100% | 38% | 35% | 83% | 5% | 12% | 0% | 70% | 28276 |
| c118_SB_8d_D1 | NANOG<sub>exo</sub> cassette | ETN<sub>2</sub> | E7 + SB | D8 | R1 | 47185418 | 47138753 | 100% | 37% | 35% | 83% | 5% | 12% | 0% | 73% | 27349 |
| c16_TGF_8d_D2 | ET parental WTC11 hiPSC line | ET | E8 | D8 | R2 | 55032124 | 54991734 | 100% | 38% | 35% | 83% | 5% | 12% | 0% | 74% | 28547 |
| c118_TGF_8d_D2 | NANOG<sub>exo</sub> cassette | ETN<sub>2</sub> | E8 | D8 | R2 | 55988082 | 55940794 | 100% | 38% | 36% | 83% | 6% | 12% | 0% | 71% | 28032 |
| c16_SB_8d_D2 | ET parental WTC11 hiPSC line | ET | E7 + SB | D8 | R2 | 61228746 | 61176610 | 100% | 38% | 36% | 83% | 5% | 12% | 0% | 74% | 28464 |
| c118_SB_8d_D2 | NANOG<sub>exo</sub> cassette | ETN<sub>2</sub> | E7 + SB | D8 | R2 | 63854035 | 63801600 | 100% | 38% | 36% | 81% | 6% | 13% | 0% | 71% | 26713 |
| c16_TGF_8d_D3 | ET parental WTC11 hiPSC line | ET | E8 | D8 | R3 | 51325973 | 51291828 | 100% | 38% | 35% | 82% | 5% | 13% | 0% | 74% | 27955 |
| c118_TGF_8d_D3 | NANOG<sub>exo</sub> cassette | ETN<sub>2</sub> | E8 | D8 | R3 | 52525744 | 52483835 | 100% | 38% | 36% | 83% | 6% | 11% | 0% | 73% | 27216 |
| c16_SB_8d_D3 | ET parental WTC11 hiPSC line | ET | E7 + SB | D8 | R3 | 61445948 | 61395814 | 100% | 37% | 35% | 83% | 5% | 11% | 0% | 74% | 27349 |
| c118_SB_8d_D3 | NANOG<sub>exo</sub> cassette | ETN<sub>2</sub> | E7 + SB | D8 | R3 | 50062175 | 50025647 | 100% | 38% | 36% | 83% | 5% | 12% | 0% | 73% | 27683 |
| c16_TGF_ct_T1 | ET parental WTC11 hiPSC line | ET | E8 | D0 | R1 | 52740019 | 52685001 | 100% | 37% | 36% | 83% | 6% | 12% | 0% | 75% | 27406 |
| c118_TGF_ct_T1 | NANOG<sub>exo</sub> cassette | ETN<sub>2</sub> | E8 | D0 | R1 | 48490053 | 48448759 | 100% | 38% | 37% | 83% | 5% | 12% | 0% | 74% | 27452 |
| c16_SB_24h_T1 | ET parental WTC11 hiPSC line | ET | E7 + SB | 24 h / D1 | R1 | 52341215 | 52300958 | 100% | 38% | 36% | 82% | 5% | 14% | 0% | 75% | 27113 |
| c118_SB_24h_T1 | NANOG<sub>exo</sub> cassette | ETN<sub>2</sub> | E7 + SB | 24 h / D1 | R1 | 53462326 | 53422156 | 100% | 38% | 37% | 81% | 5% | 13% | 0% | 72% | 27463 |
| c16_TGF_ct_T2 | ET parental WTC11 hiPSC line | ET | E8 | D0 | R2 | 63048710 | 63002507 | 100% | 38% | 35% | 82% | 5% | 14% | 0% | 75% | 27945 |
| c118_TGF_ct_T2 | NANOG<sub>exo</sub> cassette | ETN<sub>2</sub> | E8 | D0 | R2 | 39485582 | 39455637 | 100% | 37% | 36% | 82% | 4% | 14% | 0% | 73% | 26784 |
| c16_SB_24h_T2 | ET parental WTC11 hiPSC line | ET | E7 + SB | 24 h / D1 | R2 | 58846513 | 58801457 | 100% | 37% | 35% | 83% | 4% | 12% | 0% | 76% | 27686 |
| c118_SB_24h_T2 | NANOG<sub>exo</sub> cassette | ETN<sub>2</sub> | E7 + SB | 24 h / D1 | R2 | 61242864 | 61191586 | 100% | 38% | 37% | 83% | 5% | 13% | 0% | 73% | 28282 |
| c16_TGF_ct_T3 | ET parental WTC11 hiPSC line | ET | E8 | D0 | R3 | 54504614 | 54461914 | 100% | 37% | 34% | 82% | 5% | 13% | 0% | 74% | 28039 |
| c118_TGF_ct_T3 | NANOG<sub>exo</sub> cassette | ETN<sub>2</sub> | E8 | D0 | R3 | 51655443 | 51617046 | 100% | 38% | 36% | 82% | 4% | 14% | 0% | 72% | 28594 |
| c16_SB_24h_T3 | ET parental WTC11 hiPSC line | ET | E7 + SB | 24 h / D1 | R3 | 48826809 | 48790266 | 100% | 38% | 35% | 84% | 5% | 12% | 0% | 75% | 27838 |
| c118_SB_24h_T3 | NANOG<sub>exo</sub> cassette | ETN<sub>2</sub> | E7 + SB | 24 h / D1 | R3 | 45901411 | 45864675 | 100% | 38% | 36% | 81% | 5% | 14% | 0% | 71% | 27877 |
| **Summary (median)** |  |  |  |  |  | 52632881.5 | 52584418 | 100% | 38% | 36% | 83% | 5% | 12% | 0% | 73% | 27762 |
| **E-MTAB-17408** |  |  |  |  |  |  |  |  |  |  |  |  |  |  |  |  |
| GER_3_C15_R1 | GERALT MYOD1, 3'-UTR anti-NANOG miR-F | ETN<sub>2</sub>MR<sub>N-UTR</sub> | Myogenic + TET | D7 | R1 | 61472578 | 61466354 | 100% | 63% | 62% | 54% | 2% | 44% | 0% | 49% | 27573 |
| GER_3_C15_R2 | GERALT MYOD1, 3'-UTR anti-NANOG miR-F | ETN<sub>2</sub>MR<sub>N-UTR</sub> | Myogenic + TET | D7 | R2 | 49809770 | 49801774 | 100% | 63% | 63% | 54% | 2% | 44% | 0% | 48% | 26883 |

Continued on next page

| Submitted ID | Description | Cell line | Medium | Stage | Rep. | Raw pairs | Trimmed pairs | Retained (%) | R1 adapter (%) | R2 adapter (%) | Unique (%) | Multi-map (%) | Too short (%) | Mis-match (%) | Assigned (%) | Genes detected |
| --- | --- | --- | --- | --- | --- | --- | --- | --- | --- | --- | --- | --- | --- | --- | --- | --- |
| GER_Bi_C11_R1 | GERALT MYOD1, bidirectional anti-*NANOG* miR-F | ETN<sub>2</sub>MR<sub>N</sub>-BID | Myogenic + TET | D7 | R1 | 43505644 | 43502923 | 100% | 59% | 58% | 61% | 5% | 34% | 0% | 54% | 26992 |
| GER_Bi_C11_R2 | GERALT MYOD1, bidirectional anti-*NANOG* miR-F | ETN<sub>2</sub>MR<sub>N</sub>-BID | Myogenic + TET | D7 | R2 | 44163757 | 44160313 | 100% | 60% | 60% | 59% | 3% | 38% | 0% | 53% | 27079 |
| GER_CAG_C114_R1 | GERALT MYOD1, intronic anti-*NANOG* miR-F | ETN<sub>2</sub>MR<sub>N</sub>-INT | Myogenic + TET | D7 | R1 | 37562216 | 37559957 | 100% | 64% | 63% | 53% | 2% | 44% | 0% | 48% | 26217 |
| GER_CAG_C114_R2 | GERALT MYOD1, intronic anti-*NANOG* miR-F | ETN<sub>2</sub>MR<sub>N</sub>-INT | Myogenic + TET | D7 | R2 | 51502353 | 51499145 | 100% | 65% | 64% | 51% | 2% | 47% | 0% | 46% | 27876 |
| OPTIOX_1 | OPTi-OX MYOD1 | ETM | Myogenic + TET | D7 | R1 | 80941998 | 80934791 | 100% | 60% | 59% | 60% | 3% | 38% | 0% | 53% | 30119 |
| OPTIOX_3 | OPTi-OX MYOD1 | ETM | Myogenic + TET | D7 | R2 | 66122623 | 66116440 | 100% | 63% | 62% | 56% | 5% | 39% | 0% | 50% | 29426 |
| OPTIOX_4 | OPTi-OX MYOD1 | ETM | Myogenic + TET | D7 | R3 | 49605300 | 49602031 | 100% | 61% | 60% | 56% | 8% | 36% | 0% | 50% | 26018 |
| hiPSCs_1 | Undifferentiated hiPSC | ETM | E8 | D0 | R3 | 78989820 | 78987295 | 100% | 57% | 55% | 64% | 3% | 34% | 0% | 57% | 28729 |
| hiPSCs_2 | Undifferentiated hiPSC | ETM | E8 | D0 | R2 | 84022514 | 84019473 | 100% | 59% | 57% | 61% | 3% | 36% | 0% | 54% | 28935 |
| hiPSCs_3 | Undifferentiated hiPSC | ETM | E8 | D0 | R1 | 40966336 | 40964817 | 100% | 62% | 62% | 52% | 4% | 43% | 0% | 46% | 26670 |
| Summary (median) |  |  |  |  |  | 50656061.5 | 50650459.5 | 100% | 61.5% | 61% | 56% | 3% | 38.5% | 0% | 50% | 27326 |

**Supplementary Table 8 | Bulk RNA-sequencing sample metadata and quality-control metrics**
Sample identity, experimental condition, replicate identifier and sequencing quality-control metrics for the E-MTAB-17410 and E-MTAB-17408 bulk RNA-sequencing datasets. Submitted sample names are retained unchanged. Summary rows report the median across all samples within each dataset.

| Sample ID | Description | Cell line | Medium | Stage | Rep. | Cells | Mean reads/cell | Median genes/cell | Genes detected | Median counts/cell |
| --- | --- | --- | --- | --- | --- | --- | --- | --- | --- | --- |
| **10x Genomics scRNA-seq – E-MTAB-17411** |  |  |  |  |  |  |  |  |  |  |
| cl6_WT_P2_E8_n1 | ET parental WTC11 hiPSC line | ET | E8 | P3 | R1 | 1330 | 14680 | 2643 | 30931 | 5296 |
| cl6_WT_P2_E8_n2 | ET parental WTC11 hiPSC line | ET | E8 | P3 | R2 | 1228 | 16354 | 2833 | 31060 | 5907 |
| cl6_WT_P2_E8_n3 | ET parental WTC11 hiPSC line | ET | E8 | P3 | R3 | 1275 | 15106 | 2670 | 30961 | 5362 |
| c118_OVER_P2_E8_n1 | *NANOG*<sub>exo</sub> cassette | ETN<sub>2</sub> | E8 | P3 | R1 | 914 | 17741 | 2708 | 29759 | 5606 |
| c118_OVER_P2_E8_n2 | *NANOG*<sub>exo</sub> cassette | ETN<sub>2</sub> | E8 | P3 | R2 | 954 | 16975 | 2675 | 29629 | 5656 |
| c118_OVER_P2_E8_n3 | *NANOG*<sub>exo</sub> cassette | ETN<sub>2</sub> | E8 | P3 | R3 | 900 | 20569 | 3004 | 29683 | 6618 |
| cl6_WT_P2_E6_n1 | ET parental WTC11 hiPSC line | ET | E7 | P3 | R1 | 1141 | 13350 | 2525 | 29520 | 4759 |
| cl6_WT_P2_E6_n2 | ET parental WTC11 hiPSC line | ET | E7 | P3 | R2 | 1367 | 13457 | 2526 | 30038 | 4813 |
| cl6_WT_P2_E6_n3 | ET parental WTC11 hiPSC line | ET | E7 | P3 | R3 | 1162 | 13359 | 2550 | 29684 | 4867 |
| c118_OVER_P12_E6_n1 | *NANOG*<sub>exo</sub> cassette | ETN<sub>2</sub> | E7 | P10 | R1 | 1038 | 19932 | 3167 | 30752 | 7148 |
| c118_OVER_P12_E6_n2 | *NANOG*<sub>exo</sub> cassette | ETN<sub>2</sub> | E7 | P10 | R2 | 1203 | 16839 | 2823 | 30564 | 6049 |
| c118_OVER_P12_E6_n3 | *NANOG*<sub>exo</sub> cassette | ETN<sub>2</sub> | E7 | P10 | R3 | 1145 | 16168 | 2754 | 30374 | 5904 |
| **PIPseq scRNA-seq – E-MTAB-17414** |  |  |  |  |  |  |  |  |  |  |
| GERALTd4 | GERALT MYOD1 | ETN<sub>2</sub>MR<sub>N-INT</sub> | hE6 + dox | D4 | R1 | 900 | 5030 | 149 | 16524 | 1641 |
| GERALTd7 | GERALT MYOD1 | ETN<sub>2</sub>MR<sub>N-INT</sub> | hE6 + dox | D7 | R1 | 1508 | 33477 | 3854 | 26885 | 14449 |
| LONZA pre | Primary myoblasts | HSMM | SkGM-2 | D0 | R1 | 2342 | 25429 | 2995 | 26370 | 7847 |
| LONZA post | Primary myotubes | HSMM | HSMM fusion medium | D4 | R1 | 1478 | 37967 | 4103 | 25428 | 13672 |
| OPTi-OX d4 | OPTi-OX MYOD1 | ETM | hE6 + dox | D4 | R1 | 1087 | 34585 | 393 | 20555 | 540 |
| OPTi-OX d7 | OPTi-OX MYOD1 | ETM | hE6 + dox | D7 | R1 | 775 | 67765 | 4471 | 25813 | 17200 |
| WT pre | hiPSC DD myoblasts | WTC11 | SKGM | D0 | R1 | 1195 | 33508 | 210 | 19803 | 278 |
| WT d4 | hiPSC DD myotubes | WTC11 | Myoblast fusion medium | D4 | R1 | 1320 | 50862 | 4627 | 25625 | 17133 |

###### Supplementary Table 9 | Single-cell RNA-sequencing sample metadata and quality-control metrics

Sample identity, cell line, culture condition, passage or differentiation stage, replicate identifier and sequencing quality-control metrics for the 10x Genomics and PIPseq single-cell RNA-sequencing datasets. Submitted sample identifiers are retained unchanged and may therefore contain legacy condition and passage labels; the Medium and Stage columns report the corresponding experimental conditions used in the manuscript. The final column reports median UMIs per cell for E-MTAB-17411 and median molecules per cell for E-MTAB-17414. Metrics are reported after the stated sample-processing and cell-filtering procedures.

| Signature | No. genes | Genes | References | Dataset(s) |
| --- | --- | --- | --- | --- |
| Cell cycle S phase | 26 | *MCM5, PCNA, TYMS, FEN1, MCM2, MCM4, RRM1, UNG, GINS2, MCM6, CDCA7, DTL, PRIM1, UHRF1, HELLS, RFC2, RPA2, NASP, RAD51, CHEK1, ORC6, MCM3, MCM7, MLF1IP, BRIP1, E2F8* | Seurat S.Score gene set; Whitfield et al. 2002 (PMID: 12058064); Trapnell et al. 2014 (PMID: 24658644) | E-MTAB-17408; E-MTAB-17414 |
| Cell cycle G2/M | 49 | *HMGB2, CDK1, NUSAP1, UBE2C, BIRC5, TPX2, TOP2A, NDC80, CKS2, NUF2, CKS1B, MKI67, TMPO, CENPF, TACC3, SMC4, CCNB2, CKAP2L, CKAP2, AURKB, BUB1, KIF11, ANP32E, TUBB4B, GTSE1, KIF20B, HJURP, CDCA3, CDC20, TTK, CDC45, CDC6, EXO1, TIPIN, DSCC1, BLM, CASP8AP2, USP1, CLSPN, POLA1, CHAF1B, BRCA1, EXOSC2, USP16, POLD3, MSH2, ATAD2, RAD51AP1, RRM2* | Seurat G2M.Score gene set; Tirosh et al. 2016 (PMID: 27124452); Trapnell et al. 2014 (PMID: 24658644) | E-MTAB-17408; E-MTAB-17414 |
| Ribosome tone | 19 | *RPLP0, RPL3, RPL5, RPL7, RPL11, RPL13A, RPL23A, RPS3, RPS6, RPS8, RPS12, RPS18, RPS27, NCL, FBL, NPM1, EIF4EBP1, MYC, DDIT4* | Murgia et al. 2021 (PMID: 34727990); Chemello et al. 2011 (PMID: 21364935) | E-MTAB-17408; E-MTAB-17414 |
| Fiber type I (slow) | 18 | *MYH7, TNNT1, TNNT1, MYL2, ATP2A2, CKMT2, MB, COX7A1, COX6A2, PPARGC1A, SDHA, CS, SLC25A4, MYBPC1, MYL3, PDK4, ESRR, NDUFA10* | Schiaffino & Reggiani 2011 (PMID: 22013216); Murgia et al. 2021 (PMID: 34727990); Chemello et al. 2011 (PMID: 21364935); Schiaffino et al. 2025 (PMID: 39134381) | E-MTAB-17408; E-MTAB-17414 |
| Fiber type IIa (fast) | 17 | *MYH2, TNNT2, TNNT3, MYL1, ATP2A1, ACTN3, TPM2, PVALB, LDHA, PKM, PFKM, ENO1, PGAM2, ALDOA, CKM, DMD, CASQ1* | Murgia et al. 2021 (PMID: 34727990); Chemello et al. 2011 (PMID: 21364935); Schiaffino et al. 2025 (PMID: 39134381); Bloomberg & Quadrilatero 2012 (PMID: 22530000) | E-MTAB-17408; E-MTAB-17414 |
| Fiber type IIx (fast) | 12 | *MYH1, ACTN3, MYLK2, TNNT3, TNNT2, ATP2A1, TPM2, CKM, LDHA, ENO1, PFKM, MYH4* | Murgia et al. 2021 (PMID: 34727990); Uhlen et al. 2015 (Human Protein Atlas; PMID: 25613900) | E-MTAB-17408; E-MTAB-17414 |
| Developmental / embryonic | 9 | *MYH3, MYH8, MYL4, TNNT1, TNNT1, CHRNG, NEFM, NES, CCND2* | Schiaffino et al. 2011 (PMID: 22013216); Schiaffino et al. 2025 (PMID: 39134381); Chal et al. 2015 (PMID: 26237517); Xi et al. 2020 (PMID: 32396864); Hicks et al. 2018 (PMID: 29255171) | E-MTAB-17408; E-MTAB-17414 |
| Sarcomere core | 20 | *TTN, NEB, ACTN2, TCAP, MYBPC1, MYBPC2, MYL1, MYL2, TNNT1, TNNT2, TNNT1, TNNT3, TPM1, TPM2, MYOM1, MYOM2, CSRP3, ANKRD1, FHL1, MYBPH* | Cowling et al. 2008 (PMID: 19075112); Miller et al. 2003 (PMID: 14583192); GO:0030017 (sarcomere); GO:0030016 (myofibril) | E-MTAB-17408; E-MTAB-17414 |
| Calcium excitation-contraction | 14 | *RYR1, CACNA1S, ATP2A1, ATP2A2, CASQ1, CASQ2, SLN, PLN, TRDN, STAC3, JPH1, CAMK2A, CAMK2B, CAV3* | Rall 2024 (PMID: 39116389) | E-MTAB-17408; E-MTAB-17414 |
| Oxidative mitochondrial | 18 | *SDHA, SDHB, UQCRC1, NDUFA9, NDUFS1, COX4I1, COX6A2, ATP5F1A, ATP5MC1, CKMT2, SLC25A4, PPARGC1A, TFAM, CS, ACADM, NDUFB8, CYCS, PDHA1* | Lin et al. 2002 (PMID: 12181572); Murgia et al. 2021 (PMID: 34727990); Chemello et al. 2011 (PMID: 21364935); Schiaffino et al. 2025 (PMID: 39134381) | E-MTAB-17408; E-MTAB-17414 |
| Glycolytic fast | 15 | *HK2, PFKM, ALDOA, TPI1, GAPDH, PGK1, PGAM2, ENO1, PKM, LDHA, PDK1, SLC2A4, PFKFB3, ENO3, MYL1* | Schiaffino & Reggiani 2011 (PMID: 22013216); Murgia et al. 2021 (PMID: 34727990); Chemello et al. 2011 (PMID: 21364935); Schiaffino et al. 2025 (PMID: 39134381) | E-MTAB-17408; E-MTAB-17414 |
| Satellite cell core | 12 | *PAX7, VCAM1, ITGA7, CXCR4, MYF5, CDH15, SDC4, EGFR, NCAM1, SPRY1, MUSTN1, NOS1* | De Micheli et al. 2020 (PMID: 32624006); GEO: GSE143704; Yin et al. 2013 (PMID: 23303905) | E-MTAB-17408; E-MTAB-17414 |
| Myoblast proliferation cycle | 13 | *MYOD1, MYF5, MKI67, TOP2A, PCNA, CCNB1, CDK1, E2F1, NUSAP1, HMGB2, UBE2C, AURKB, RRM2* | Trapnell et al. 2014 (PMID: 24658644); Nalbandian et al. 2022 (PMID: 35459735); Hicks et al. 2018 (PMID: 29255171) | E-MTAB-17408; E-MTAB-17414 |

Continued on next page

| Signature | No. genes | Genes | References | Dataset(s) |
| --- | --- | --- | --- | --- |
| Myotube differentiation & fusion | 10 | *MYOG, MEF2C, MYMK, MYMX, MYH3, MYH8, CKM, DES, DNM2, ACTA1* | Chal et al. 2015 (PMID: 26237517); Sampath et al. 2018 (PMID: 29386054); Hicks et al. 2018 (PMID: 29255171) | E-MTAB-17408; E-MTAB-17414 |
| Myogenic regulators axis | 7 | *PAX3, PAX7, MYF5, MYOD1, MYOG, MEF2C, MYF6* | Weintraub et al. 1991 (PMID: 1846704); Rudnicki et al. 1993 (PMID: 8269513) | E-MTAB-17408; E-MTAB-17414 |
| Neuromuscular junction | 13 | *AGRN, LRP4, MUSK, DOK7, RAPSN, COLQ, CHRNA1, CHRNB1, CHRND, CHRNE, CHRNG, ACHE, UTRN* | Burden et al. 2018 (PMID: 29415504) | E-MTAB-17408; E-MTAB-17414 |
| Muscle ECM / basement membrane | 18 | *LAMA2, LAMC1, COL6A1, COL6A2, COL6A3, COL1A1, COL1A2, COL3A1, FN1, SPP1, LUM, DCN, DAG1, ITGA7, THY1, COL4A1, COL4A2, COL5A1* | De Micheli et al. 2020 (PMID: 32624006; GEO: GSE143704) | E-MTAB-17408; E-MTAB-17414 |
| Fibro-adipogenic progenitors | 12 | *PDGFRA, COL1A1, COL3A1, DCN, LUM, CXCL14, THY1, GPC6, FBN1, DPT, POSTN, PCOLCE* | De Micheli et al. 2020 (PMID: 32624006; GEO: GSE143704); Negroni et al. 2022 (PMID: 36200044); Zhao et al. 2024 (PMID: 39286484) | E-MTAB-17408; E-MTAB-17414 |
| Atrophy / stress | 12 | *FBXO32, TRIM63, DDIT4, GADD45A, ATF4, FOXO3, PSMA1, PSMB1, SESN1, SESN3, EIF4EBP1, PPP1R15A* | Bodine et al. 2001 (PMID: 11679633); Gomes et al. 2001 (PMID: 11717410) | E-MTAB-17408; E-MTAB-17414 |
| Pluripotency core | 9 | *POU5F1, NANOG, SOX2, PRDM14, DPPA2, DPPA4, ZFP42, DNMT3A, DNMT3B* | Theunissen et al. 2016 (PMID: 27424783); Nichols & Smith 2009 (PMID: 19497275) | E-MTAB-17408; E-MTAB-17414 |
| Naive pluripotency | 8 | *KLF17, KLF4, TFCP2L1, DNMT3L, DPPA3, TBX3, ESRRB, SUSD2* | Rostovskaya et al. 2022 (PMID: 34870831); Pastor et al. 2016 (PMID: 26853856) | E-MTAB-17408; E-MTAB-17414 |
| Mesoderm specification | 13 | *MESP1, PDGFRA, KDR, ISL1, HAND1, HAND2, GATA4, TBX5, NKX2-5, MEF2C, FGF8, EOMES, TBXT* | Chan et al. 2013 (PMID: 23896987); Chan et al. 2013 (PMID: 23642367); Loh et al. 2016 (PMID: 27419872); Bulger et al. 2024 (PMID: 38411343) | E-MTAB-17408; E-MTAB-17414 |
| Early myogenic commitment | 9 | *PAX3, PAX7, MYF5, MYOD1, MET, ITGA7, FGFR4, LBX1, MYOG* | Chal et al. 2015 (PMID: 26237517); Hicks et al. 2018 (PMID: 29255171); Hicks et al. 2023 (PMID: 37919520) | E-MTAB-17408; E-MTAB-17414 |
| *Additional signatures used for E-MTAB-17414* |  |  |  |  |
| Active myotube fusion | 8 | *MYOG, MYMK, MYMX, MEF2C, DNM2, ACTA1, CKM, DES* | Millay et al. 2013 (PMID: 23868259); Leikina et al. 2018 (PMID: 30197239); Chen, You, Wang et al. 2020 (PMID: 31642939) | E-MTAB-17414 |
| Nascent post-fusion myotube | 12 | *ANKRD1, MYBPH, FHL1, CDKN1A, TNNT2, TNNT3, BIN1, COX6A2, CRYAB, F13A1, PTGDS, SIRT2* | Cowling et al. 2008 (PMID: 19075112); Halevy et al. 1995 (PMID: 7863327) | E-MTAB-17414 |
| Satellite cell core (hiPSC) | 5 | *ITGA7, MYF5, CDH15, NCAM1, SPRY1* | De Micheli et al. 2020 (PMID: 32624006; GEO: GSE143704); Yin et al. 2013 (PMID: 23303905) | E-MTAB-17414 |
| Oxidative stress / antioxidant | 8 | *NQO1, HMOX1, TXNIP, FTH1, MT1E, MT2A, SLC7A11, SRXN1* | Ma 2013 (PMID: 23294312) | E-MTAB-17414 |
| Slow-twitch myofiber (NOVO) | 11 | *TNNT1, MYL3, TPM3, MYBPC1, ATP2A2, TPM2, MYL6B, MYH7, TNNI1, TNNC1, MYL2* | bioRxiv 2026.01.16.698404; van der Wal et al. 2023 (PMID: 37774701) | E-MTAB-17414 |
| Fast-twitch myofiber (NOVO) | 10 | *TPM1, MYBPC2, ATP2A1, MYH2, MYL1, TNNI2, TNNC2, TNNT3, MYLPF, ACTN3* | bioRxiv 2026.01.16.698404; van der Wal et al. 2023 (PMID: 37774701) | E-MTAB-17414 |
| Skeletal muscle, fiber-type independent (NOVO) | 12 | *TPM4, VIM, DES, OBSCN, ACTN2, RYR1, DMD, TMOD1, ACTA1, CKM, NEB, TTN* | bioRxiv 2026.01.16.698404; van der Wal et al. 2023 (PMID: 37774701) | E-MTAB-17414 |
| Genes not expressed in mature skeletal muscle (NOVO) | 7 | *MYH8, MYH6, MYH3, MYL6, MYL4, TNNT2, MYOG* | bioRxiv 2026.01.16.698404; van der Wal et al. 2023 (PMID: 37774701) | E-MTAB-17414 |

Continued on next page

| Signature | No. genes | Genes | References | Dataset(s) |
| --- | --- | --- | --- | --- |
| Glycogen metabolism (NOVO) | 12 | *GYS1, PYGM, PHKA1, PHKB, PHKG1, GBE1, PPP1R3A, PPP1R3C, GYG1, PGM1, UGP2, STBD1* | bioRxiv 2026.01.16.698404 | E-MTAB-17414 |
| AMPK energy sensing (NOVO) | 9 | *PRKAA1, PRKAA2, PRKAB1, PRKAB2, PRKAG1, PRKAG3, STK11, CAB39, CAB39L* | bioRxiv 2026.01.16.698404 | E-MTAB-17414 |
| $\beta_2$-adrenergic response (NOVO) | 10 | *ADRB2, PPARGC1A, NR4A1, NR4A2, NR4A3, PGK1, CREB1, LIPE, RPS6, GYS1* | bioRxiv 2026.01.16.698404 | E-MTAB-17414 |
| Insulin / GLUT4 signaling (NOVO) | 10 | *SLC2A4, SLC2A1, SLC2A3, INSR, IRS1, IRS2, PIK3R1, AKT2, TBC1D4, RAB10* | bioRxiv 2026.01.16.698404 | E-MTAB-17414 |
| Exercise training adaptation (NOVO) | 6 | *SLC2A4, HK2, PDHA1, PPARGC1A, NR4A3, VEGFA* | bioRxiv 2026.01.16.698404 | E-MTAB-17414 |
| Hypoxia / glucose / fatigue stress (NOVO) | 9 | *HIF1A, SLC16A1, SLC16A3, PDK1, LDHA, PFKFB3, VEGFA, CA9, BNIP3* | bioRxiv 2026.01.16.698404 | E-MTAB-17414 |
| Tissue maturation, 3D vs monolayer (NOVO) | 5 | *MYBPC1, MYL1, MYH2, SDHA, DMD* | bioRxiv 2026.01.16.698404 | E-MTAB-17414 |

**Supplementary Table 10 | Curated gene signatures used for transcriptomic analyses**

Gene sets used to characterize cell-cycle state, pluripotency, developmental and myogenic states, muscle-fiber properties, metabolism and stress responses in the E-MTAB-17408 bulk RNA-sequencing and E-MTAB-17414 single-cell RNA-sequencing datasets. For each signature, the number and identity of constituent genes, supporting references and dataset of application are reported.
